# CmP signaling network unveils novel biomarkers for triple negative breast cancer in African American women

**DOI:** 10.1101/2021.05.24.445510

**Authors:** Johnathan Abou-Fadel, Brian Grajeda, Xiaoting Jiang, Alyssa-Marie D. Cailing-De La O, Esmeralda Flores, Akhil Padarti, Muaz Bhalli, Alexander Le, Jun Zhang

## Abstract

Breast cancer is the most commonly diagnosed cancer worldwide and remains the second leading cause of cancer death. While breast cancer mortality has steadily declined over the past decades through medical advances, an alarming disparity in breast cancer mortality has emerged between African American women (AAW) and Caucasian American women (CAW); and new evidence suggests more aggressive behavior of triple-negative breast cancer (TNBC) in AAW may contribute to racial differences in tumor biology and mortality. Progesterone (PRG) is capable of exerting its cellular effects through either its classic, non-classic or combined responses through binding to either classic nuclear PRG receptors (nPRs) or non-classic membrane PRG receptors (mPRs), warranting both pathways an equally important status in PRG-mediated signaling. In our previous report, we demonstrated that the CCM signaling complex (CSC) consisting of CCM1, CCM2, and CCM3 proteins can couple both nPRs and mPRs signaling cascades to form a CSC-mPRs-PRG-nPRs (CmPn) signaling network in nPR positive(+) breast cancer cells. In this report, we furthered our research by establishing the CSC-mPRs-PRG (CmP) signaling network in nPR(-) breast cancer cells, demonstrating that a common core mechanism exists, regardless of nPR(+/-) cell type. This is the first report stating that inducible expression patterns exist between CCMs and major mPRs in TNBC cells. Furthermore, we firstly show mPRs in TNBC cells are localized in the nucleus and participate in nucleocytoplasmic shuttling in a coordinately synchronized fashion with CCM proteins under steroid actions, following the same cellular distribution as other well-defined steroid hormone receptors. Finally, for the first time, we deconvoluted the CmP signalosome by using multi-omics approaches, which helped us understand key factors within the CmP network, and identify 21 specific biomarkers with potential clinical applications associated with AAW-TNBC tumorigenesis. These novel biomarkers could have immediate clinical implications to dramatically improve health disparities among AAW-TNBCs.

## Introduction

Breast cancer is the most commonly diagnosed cancer worldwide and remains the second leading cause of cancer death in the United States (Allicock et al., 2013; Boyer-Chammard et al., 1999). While breast cancer mortality has steadily declined over the past three decades through advances in medical care and technology (Wells et al., 2014), an alarming Black-White disparity in breast cancer mortality has emerged in the U.S. As compared to Caucasian American Women (CAW) with breast cancer, African American Women (AAW) are less likely to be diagnosed at an early stage (Baquet et al., 2008), face more aggressive forms of the disease (Dignam, 2000; Newman, 2014; Newman, 2017), and experience a higher mortality rate (Baquet et al., 2008). Additionally, despite the lower lifetime incidence of breast cancer, AAW diagnosed with breast cancer are 38% more likely to die from the disease than age-matched and prognosis-matched CAW (Srivastava et al., 2019; Wells et al., 2014). Overall, AAW face disproportionately higher breast cancer mortality rates than that of CAW (Allicock et al., 2013; Newman, 2014; Newman, 2017; Wells et al., 2014). In addition to key social, political, and economic factors that impact the biological expression of breast cancer in AAW and CAW, mounting evidence indicates the presence of intrinsic molecular factors that contribute to racial disparities in breast cancer mortality. AAW were independently associated with poorer survival and worse prognosis than age- and clinically-matched CAW, suggesting that breast cancer is a more biologically aggressive disease in AAW under equal treatment (Bauer et al., 2007). Studies found that poorer rates of breast cancer survival are associated with an increased frequency of triple-negative breast cancer (TNBC), especially in young AAW (Baquet et al., 2008; Chen and Li, 2015; Sturtz et al., 2014). This suggests that a higher prevalence of TNBC and a lower prevalence of luminal A tumors could contribute to the poor prognosis of young AAW with breast cancer. TNBC is a subtype of breast cancer that is defined by the absence of estrogen, progesterone, and HER-2 receptors [ER(-), nPR(-), HER-2(-)] (Carey et al., 2006). While TNBC accounts for only ∼15% of all breast cancers, it exhibits the most aggressive metastatic behavior (Chacon and Costanzo, 2010; Perou et al., 2000; Sharma, 2018) and limited targeted therapies exist for TNBCs (Goldhirsch et al., 2011). AAW with TNBCs have worse clinical outcomes than that of CAW (Lund et al., 2009; Newman, 2014), suggesting that the higher prevalence and more aggressive biology of TNBC in AAW (Danforth, 2013) may contribute to racial differences in tumor biology (Beverly et al., 1987; Chen et al., 1994; Joslyn, 1995; Joslyn, 2002; Shiao et al., 1997). Overall, the biological basis of the four major disparities of breast cancer in AAW include, 1) early age of onset, 2) rapidly advancing stage of the disease, 3) aggressive histological changes, and 4) decreased survival, cannot be an oversight. Although the racial disparity, in both prevalence of and risks of breast cancer between AAW and CAW, may be attributable to social forces to some degree (Gerend and Pai, 2008; Hall et al., 2005; Ma et al., 2017; McCullough et al., 2005; Newman and Kaljee, 2017), decreased survival for AAW with TNBC remain the same after controlling for socioeconomic factors, treatment delay, and breast cancer receptor expression (Lund et al., 2009). This provides strong evidence that after controlling for environment/treatment disparities, biological differences seem to significantly contribute to decreased survival of AAW with TNBC.

Progesterone (PRG), a sex steroid hormone, is capable of exerting its cellular effects through either its classic, non-classic or combined responses through binding to either classic nuclear PRG receptors (nPRs) or non-classic membrane PRG receptors (mPRs), warranting both pathways an equally important status in PRG-mediated signaling. In our previous report, we demonstrated that the CCM signaling complex (CSC) consisting of CCM1, CCM2, and CCM3 can couple both nPRs and mPRs signaling cascades to form a CSC-mPRs-PRG-nPRs (CmPn) signaling network in nPR positive(+) breast cancer cells (Abou-Fadel et al., 2020b). Our previous data support that the CSC plays an important role during breast cancer tumorigenesis by coupling classic nPRs and non-classic mPRs (PAQRs) under PRG-induced actions (Abou-Fadel et al., 2020b).

Among TNBCs, over 70% of TNBCs are basal phenotype breast cancers (BPBC), one of the most aggressive TNBC cancers; mPRs role in tumorigenesis of BPBC is of great interest since BPBC derived cells do not express nPRs (Zuo et al., 2010). In this report, we will utilize two BPBC lines, the most commonly studied basal A (BaA) MDA-MB-468 (MB468) and basal B (BaB) MDA-MB-231(MB231) cells, both only expressing mPRs (Dressing et al., 2012; Dressing and Thomas, 2007; Pang and Thomas, 2011) (Suppl. Fig. 1A), to investigate the roles and relationships of key players of newly defined CSC-mPRs-PRG (CmP) network upon steroid actions. More importantly, MB468 is a well-known AAW-derived TNBC cell line, while MB231 is a CAW-derived TNBC cell line. Previous observations suggested that there is a biological basis for racial disparities in breast cancer mortality under the actions of sex hormones (Beverly et al., 1987; Chen et al., 1994; Joslyn, 1995; Joslyn, 2002; Shiao et al., 1997). Therefore, we hypothesized that there is a distinct molecular regulatory mechanism of the CmP signaling network under sex hormone actions during tumorigenesis, of AAW-derived TNBCs, providing a biological basis for racial disparities in breast cancer mortality.

The overall objective of this project is to understand the molecular regulatory mechanisms of racial disparities in breast cancer mortality concerning breast cancer tumorigenesis under sex hormone actions for the first time through molecular and cellular manipulation of cultured TNBC cell lines *in-vitro*. We also aim to systematically and comparatively analyze the dynamic role of the CmP network by deconvoluting the CmP signalosome utilizing multi-omics approaches. Indeed, our data demonstrated that a common core mechanism exists among nPR(-) breast cancer cells, termed the CmP signaling network, regardless of nPR(+/-) cell type, which partially overlaps with the CmPn signaling network in nPR(+) breast cancer cells under steroid actions. These findings indicate a more essential role of the CSC on the stability of mPRs (PAQRs) in nPR(-) cells under steroid actions. This is the first report stating that inducible expression patterns exist between CCMs and major mPRs in multiple nPR(-) cells. Furthermore, mPRs in TNBCs are localized in the nucleus and participate in nucleocytoplasmic shuttling in a coordinately synchronized fashion with CCM proteins under steroid actions, following the same cellular distribution as other well-defined steroid hormone receptors. Twenty-one potential biomarkers associated with altered expression in AAW-TNBC cells/tissues, consistent at both the transcriptional and translational levels, were also identified through our systems biology approach, which is quite significant for future AAW-TNBC therapeutic strategies. The knowledge gained from this project will have a great impact on deciphering the molecular and cellular mechanisms underlying racial disparities in breast cancer mortality through the CmP signaling network in AAW-TNBCs, which could establish a solid foundation for future therapeutic strategies.

## Results

### Distinct responses to combined sex steroid (PRG+MIF) actions through non-classic progesterone signaling in nPR(-) cells

Two TNBC/BPBC lines, MB468 and MB231 cells, are nPR(-)/mPR(+) (Dressing et al., 2012; Dressing and Thomas, 2007; Pang and Thomas, 2011) (Suppl. Fig. 1A). Therefore, both TNBC/BPBC lines are also known as good nPR(-) breast cancer cell models to examine the cellular roles and relationship between the CSC and mPRs within the CmP network upon combined steroid actions. Human leukemia-derived cell line, Jurkat cell, was defined as only bearing mPRs without any other steroid receptor (Areia et al., 2015; Dosiou et al., 2008), making it a great model to investigate the cellular function of mPRs. A normal nPR(-) breast cell line, MCF10A (10A), was also used as a control. Firstly, we investigated the relative expression levels of total mPRs (PAQRs) with combined steroid (PRG+MIF) treatment on these nPR(-) cell lines. Interestingly, significantly increased protein expression of major mPRs were observed (Fig. 1A), in contrast to previous nPR(+) data (decreased mPRs in T47D, no change in 293T cells) (Abou-Fadel et al., 2020a; Abou-Fadel et al., 2020b). Increased protein expression of PAQR7 and PAQR8 induced by PRG alone has been reported in MB231 cells (Pang and Thomas, 2011), but this is the first report stating that combined steroid treatment can also induce overexpression of all major mPRs in multiple nPR(-) cell lines. Overexpression of all major mPRs induced with combined steroid treatment in 10A, a normal breast cell line, also indicates that this treatment-induced overexpression of mPRs is likely not associated with breast cancer tumorigenesis. Hormone-induced autologous receptor down-regulation is a common property of most steroid hormones through decreasing the half-life of occupied receptors leading to their more rapid decay, which includes classic PRG receptors (nPRs). PRG bound-nPRs will down-regulate their levels by inhibiting transcription of the nPR gene. Similar autologous down-regulation of mPRs were also observed in nPR(+) T47D cells (Abou-Fadel et al., 2020b), making our current findings a novel and interesting phenomena in nPR(-) cells.

**Fig. 1.**
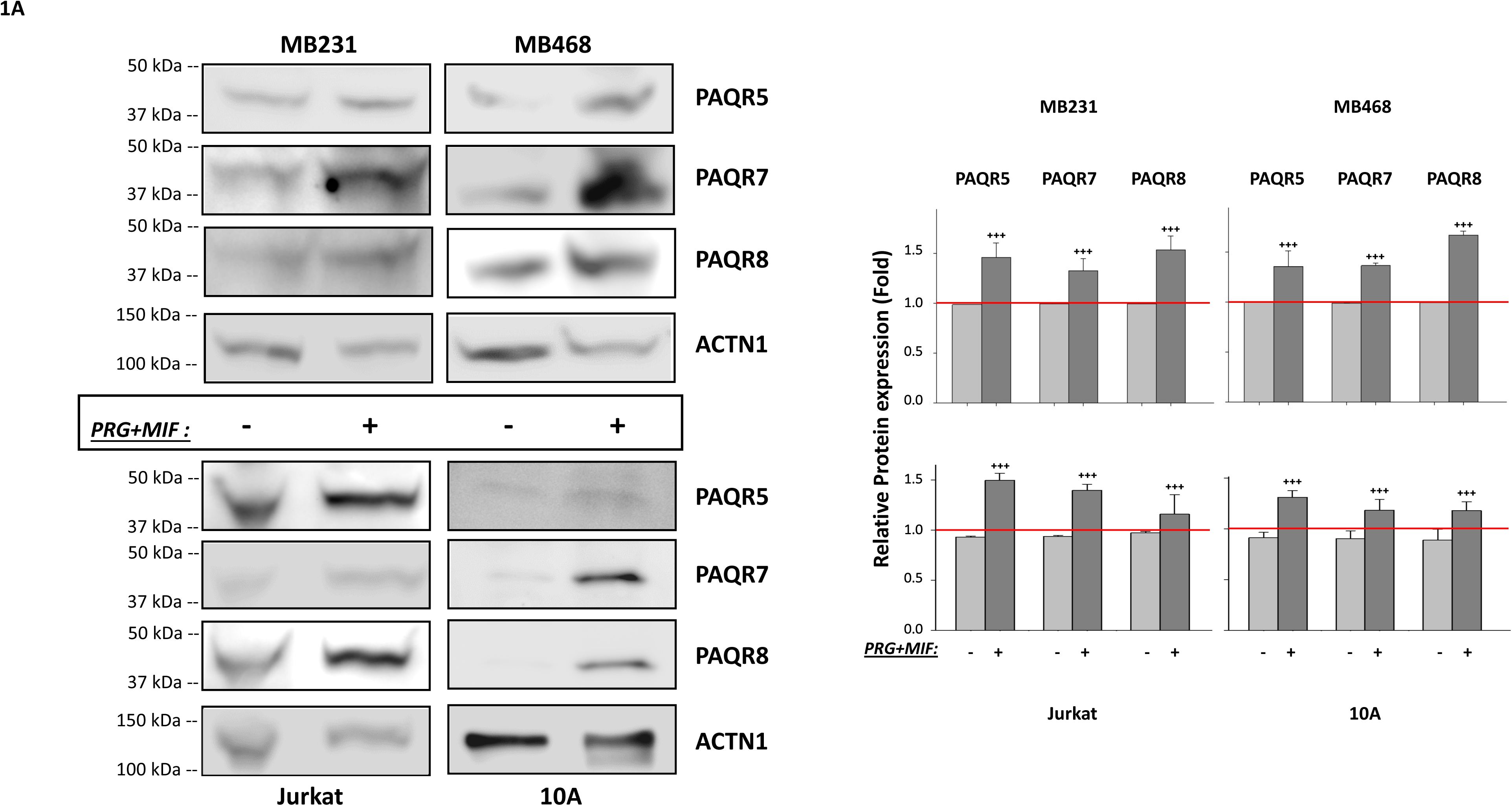
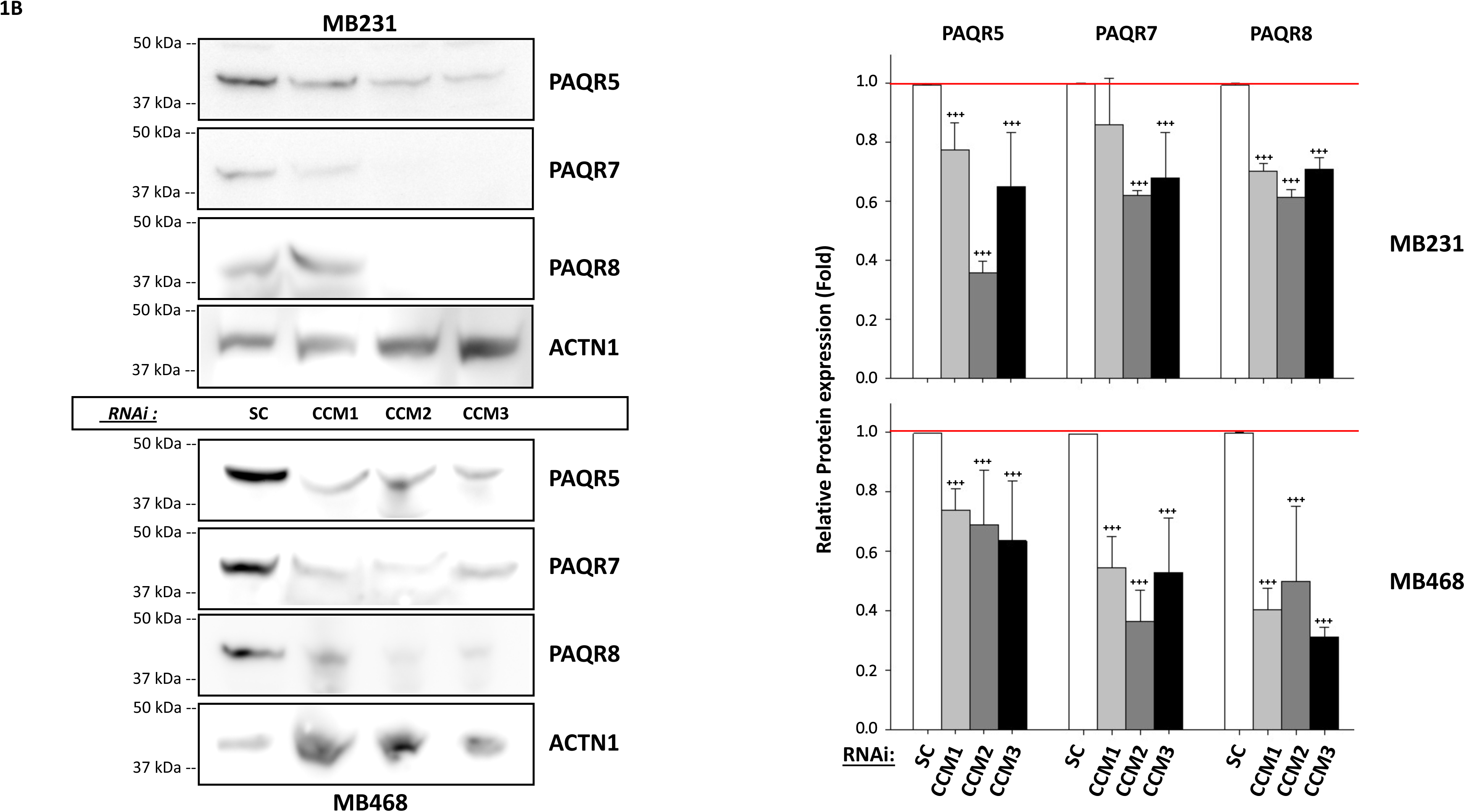
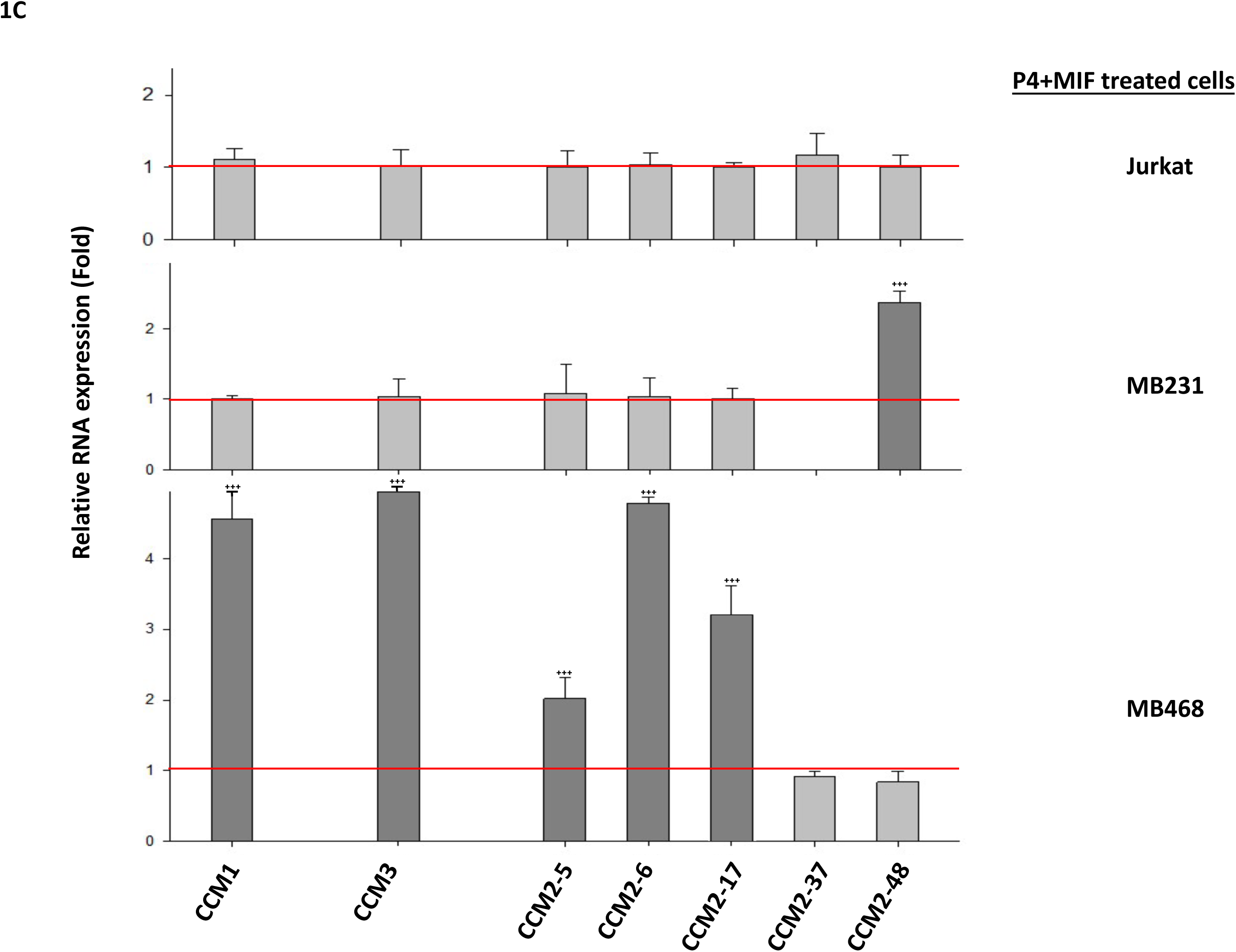
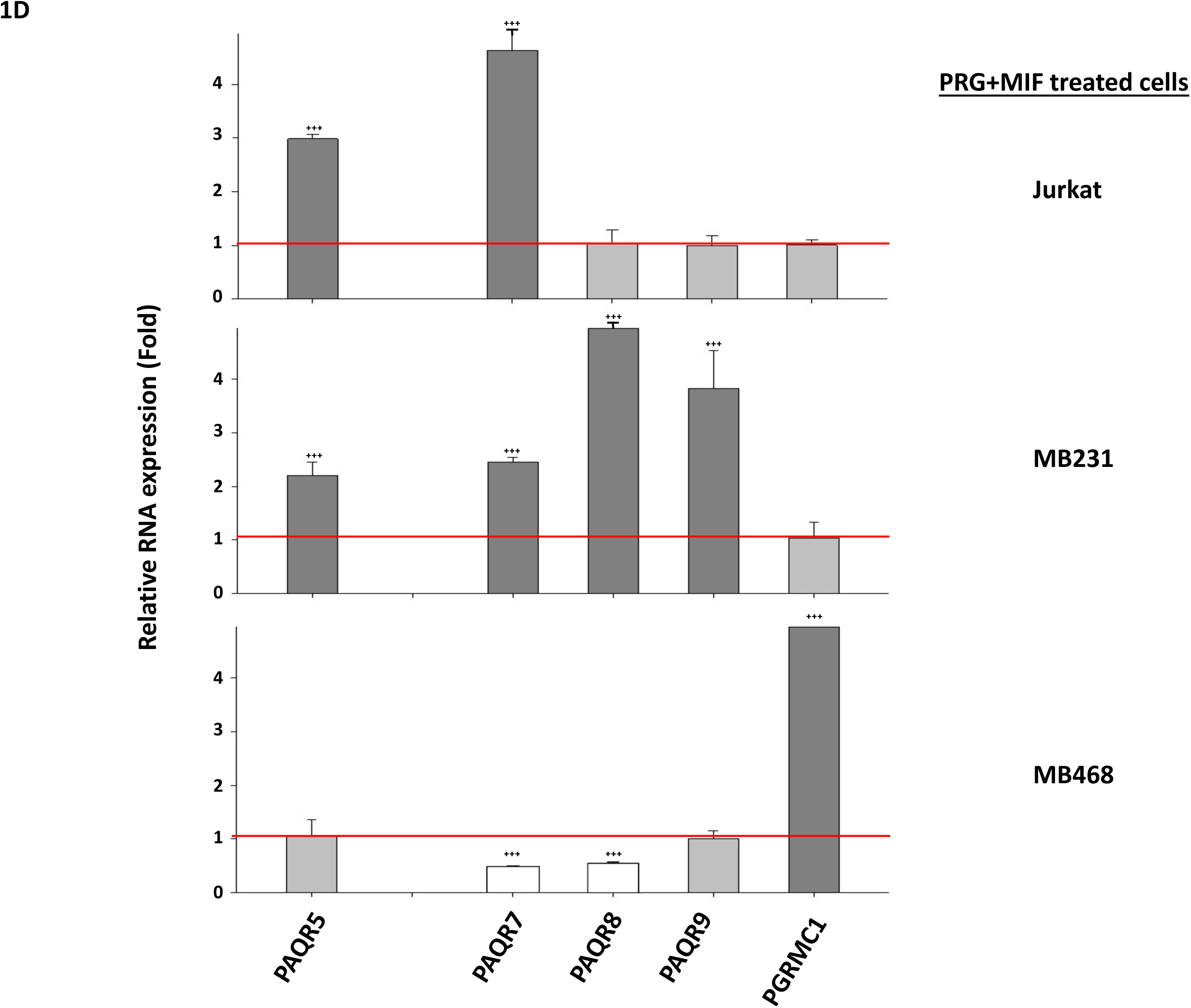
Expression levels of key factors of the CCM/mPR (CmP) network in nPR(-) cells are modulated by sex steroids. The expression profiles of key factors of the CmP network [all three *CCMs (1, 2, 3)* and major mPRs (*PAQR 5, 7, 8, 9*)] were investigated among different nPR(-) cell lines, 48 hrs after combined steroid treatment (PRG+MIF, 20 µM each). **A**. Protein expression levels of major mPRs (PAQR 5, 7, 8) are enhanced by combined steroid treatment. After combined steroid treatment for 48 hrs, western blots demonstrated dramatically increased expression levels of major mPR proteins [PAQR 5 (mPRγ), PAQR7 (mPRα), and PAQR8 (mPRβ)] in 4 different nPR(-) cell lines, MDA-MB-231(MB231), MDA-MB-468 (MB468), JURKAT (Jurkat) and MCF10A (10A) cells, compared to vehicle control (VEH) (left panels), and the increased protein expression levels are statistically significant (right panel). The relative expression levels of mPR proteins were measured through quantification of band intensities and normalized against α-actinin (ACTN1) and vehicle control (VEH) (n=3). **B**. Protein expression levels of major mPRs [PAQR 5 (mPRγ), PAQR7 (mPRα), and PAQR8 (mPRβ)] are suppressed by silencing *CCM1, CCM2,* and *CCM3* genes in 2 triple-negative breast cancer (TNBC) cell lines, MB468 and MB231. After silencing any one of three CCMs (CCM1, CCM2 or CCM3) for 48 hrs, decreased protein expression levels of major mPRs (PAQR 5, 7, 8) were visualized in both TNBC cell lines (left panel), and the decreased relative protein expression levels are statistically significant (right panel). The relative protein expression level of mPRs (PAQR 5, 7, 8) were measured through quantification of band intensities from three different experiments and normalized against α-actinin (ACTN1) followed by scramble control (SC) treatment (right panel red line) (n=3). **C**. RNA expression levels of CCMs (*CCM1, CCM2* isoforms, *CCM3*) genes are influenced by combined sex steroid treatments. After combined steroid treatment for 48 hrs, the RNA expression levels of all three CCMs (1, 2 or 3) genes in 3 different nPR(-) cell lines, MB231, MB468, and Jurkat were measured, in comparison to vehicle controls (red line). The RNA expression pattern of CCMs (1, 2 or 3) genes in MB468 are opposite to that of the other two nPR(-) cell lines, MB231 and Jurkat cells. Significantly increased RNA expression of *CCMs (1, 2, 3)* genes was observed in MB468 cells while no RNA expression changes of *CCMs (1, 2, 3)* genes was seen in either MB231 or Jurkat cells. The relative RNA expression changes of CCMs (1, 2 or 3) genes were measured by qPCR (Fold changes). The relative RNA expression changes of CCMs (1, 2 or 3) genes in MB231, MB468, and Jurkat cells were represented with bar plots where gray bars represent no change, and dark gray bars represent increased relative RNA expression, (triplicates per experiment, n=3). **D**. RNA expression levels of membrane progesterone receptors [mPRα (PAQR7), mPRβ (PAQR8), mPRγ (PAQR5), mPRε (PAQR9) and PGRMC1] are influenced by combined steroid treatments. After combined steroid treatment for 48 hrs, RNA expression level of major mPRs (PAQR 5, 7, 8, 9, PGRMC1) in 3 different nPR(-) cell lines, MB231, MB468, and Jurkat cells were measured, in comparison to vehicle control (red line). Similarly, the RNA expression pattern of major mPRs (*PAQR 5, 7, 8, 9, PGRMC1*) genes in MB468 were totally opposite to that of the other two nPR(-) cell lines, MB231 and Jurkat cells. Significantly increased RNA expression of major mPRs (PAQR5, 7, 8, 9) genes (except PGRMC1) was observed in MB231 while Jurkat cells displayed only significant increased expression in PAQR5,7. Decreased expression of PAQR7,8 and increased RNA expression of PGRMC1 was observed in MB468 cells under combined steroid actions. The relative RNA expression changes of major mPRs (PAQR 5, 7, 8, 9, PGRMC1) in 3 different nPR(-) cell lines were measured by qPCR (Fold changes) and represented with bar plots where the white bars represent significantly decreased expression, while light gray bars represent no change and dark gray bars represent significantly increased expression, compared to vehicle controls (VEH) (triplicates per experiment, n=3). In all bar plots, red line is the control baseline for fold change measurements (-/+). **, *** above bar indicates P ≤ 0.01 or 0.001 for paired *t*-test, respectively.

### Common relationships among the CmP signaling network in nPR(-) cells

After silencing *CCM (1, 2, 3)* genes respectively in both nPR(-) AAW-TNBC and CAW-TNBC cells, a significantly decreased protein expression of major mPRs (PAQR5, 7, 8) was observed compared to SC controls, especially in *CCM2*-KD TNBC cells (Fig. 1B, left panel). This decreased protein expression of major mPRs (PAQR5, 7, 8) caused by CCMs deficiency is in concordance with our data from nPR(+) T47D cells (Abou-Fadel et al., 2020b) but much more significant (Fig. 1B, right panel). These data suggest that a common core mechanism exists, termed the CmP signaling network, regardless of nPR(+/-) cell types. Furthermore, it also indicates a more essential role of the CSC on the stability of non-classic mPRs in nPR(-) cells under steroid actions.

### New subtypes of TNBC cells are based on the role and expression patterns of the CSC and mPRs

The expression levels of all CCMs (1, 2, 3) proteins are very low in the two nPR(-) TNBC cells compared to nPR(+) cells, 293T and various endothelial cells (Abou-Fadel et al., 2020a; Abou-Fadel et al., 2020b), and as a result, the effects of combined steroid actions on the protein expression of CCM3 in these nPR(-) cells were not detected (Suppl. Fig. 1B); however, given the impact of knocking down any member of the CSC and its impact on mPRs expression (Fig. 1B), suggests that the effect of combined steroid actions on the CSC through mPRs is effective even with minimal CSC expression in this group of nPR(-) cells. Increased protein expression of major mPRs (mPRα/β) under PRG treatment has been reported in MB231 cells (Pang and Thomas, 2011). Increased RNA expression levels of CCMs were observed under combined steroid actions for 48 hrs in AAW-TNBC cells only, while unchanged RNA expression of *CCMs* genes was observed in the other two nPR(-) cells (Fig. 1C). Increased RNA expression levels of mPRs were observed under combined steroid actions for 48 hrs in CAW-TNBC and nPR(-) Jurkat cells, while decreased/unchanged RNA expression of *mPRs* genes was observed AAW-TNBC cells (Fig. 1D). Interestingly, the RNA expression pattern of *mPRs* and *CCMs* genes in AAW-TNBC cells is opposite, yielding unchanged and/or decreased RNA expression of major *mPRs* genes and increased RNA expression of *CCMs* genes with combined steroid treatment (Figs. 1C, 1D). These data suggest that RNA profiling of both *mPRs* and *CCMs* genes can be used as potential biomarkers to distinguish different subtypes among TNBC cells, defined here as type-2A (CAW-TNBCs) and type-2B (AAW-TNBCs).

## TNBC cell performance is determined by the CmP network under steroid actions

### TNBC cell performance is determined by the CmP signaling network

Based on our previous findings, under sterol actions, the intricate balance among key players of the CmPn signaling network is achieved through a balance between the negative effects of PRG/MIF signaling via mPRs and the positive effects of PRG through nPRs signaling in nPR(+) breast cancer cells (Abou-Fadel et al., 2020b; Abou-Fadel et al., 2020c). Our goal is to investigate the more fragile relationship within the CmP signaling network on the motility of nPR(-) breast cancer cells by tipping the delicate balance of the CmP network under steroid actions. Two TNBC cell lines, CAW-TNBC and AAW-TNBC cells, were treated with combined steroids and cell migration, wound healing and transwell invasion were measured *in-vitro*. CAW-TNBC cells displayed no change of cell migration with PRG, compared to vehicle controls, while MIF appeared to enhance cell migration and the combined steroid effect was in-between the individual responses, both effects seen at earlier stages only (Fig. 2A, upper panel). Although PRG has been reported to decrease serum starvation-induced cell death in AAW-TNBC cells via mPRα (Dressing et al., 2012), our data show that AAW-TNBC cells displayed no effect in cell migration with PRG, compared to vehicle controls, while MIF as well as the combined steroids, appeared to enhance cell migration at earlier stages only (Fig. 2A, lower panel), similar to our data obtained in CAW-TNBC cells. Similar trends in wound healing experiments were observed. CAW-TNBC cells displayed unchanged wound closure ability with PRG and combined steroids, while MIF appeared to enhance wound closure significantly, compared to vehicle controls (Fig. 2B, upper panel, Suppl. Fig. 2A). AAW-TNBC cells displayed no effect in wound closure with PRG, compared to vehicle controls, while MIF as well as the combined steroids, appeared to enhance wound closure ability (Fig. 2B, lower panel, Suppl. Fig. 2B). Interestingly, using cell invasion assays, trends of decreased cell invasion for both TNBC cell lines under combined steroid actions were shown, compared with vehicle controls, however, without any statistical significance (Fig. 2C). These findings are in line with previous data that sex steroids can have actions directly on mPRs to direct overall cell performance of nPR(-) breast cancer cells and even their metastatic potential (Valadez-Cosmes et al., 2016; Xie et al., 2012; Zuo et al., 2010).

**Fig. 2.**
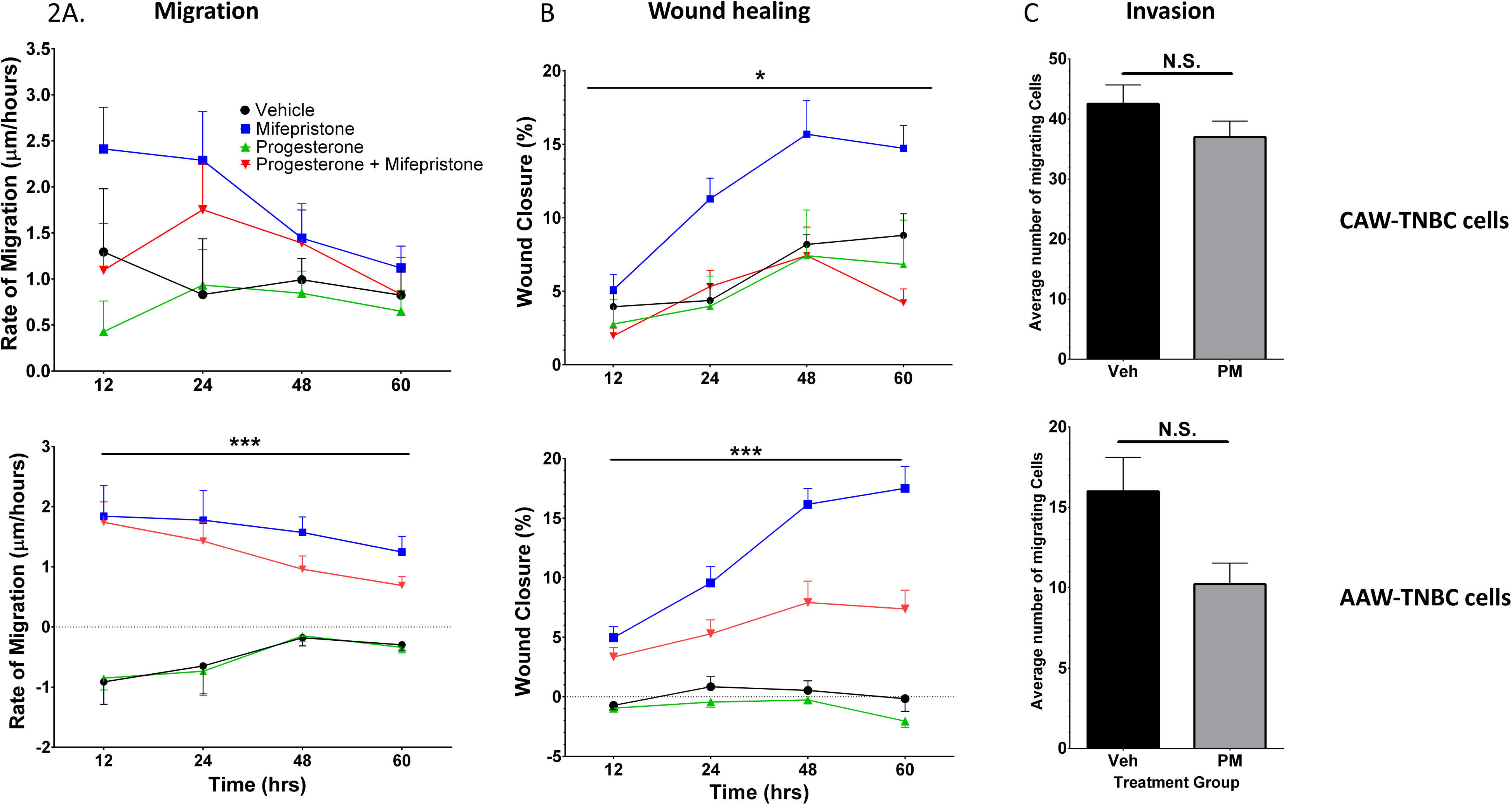
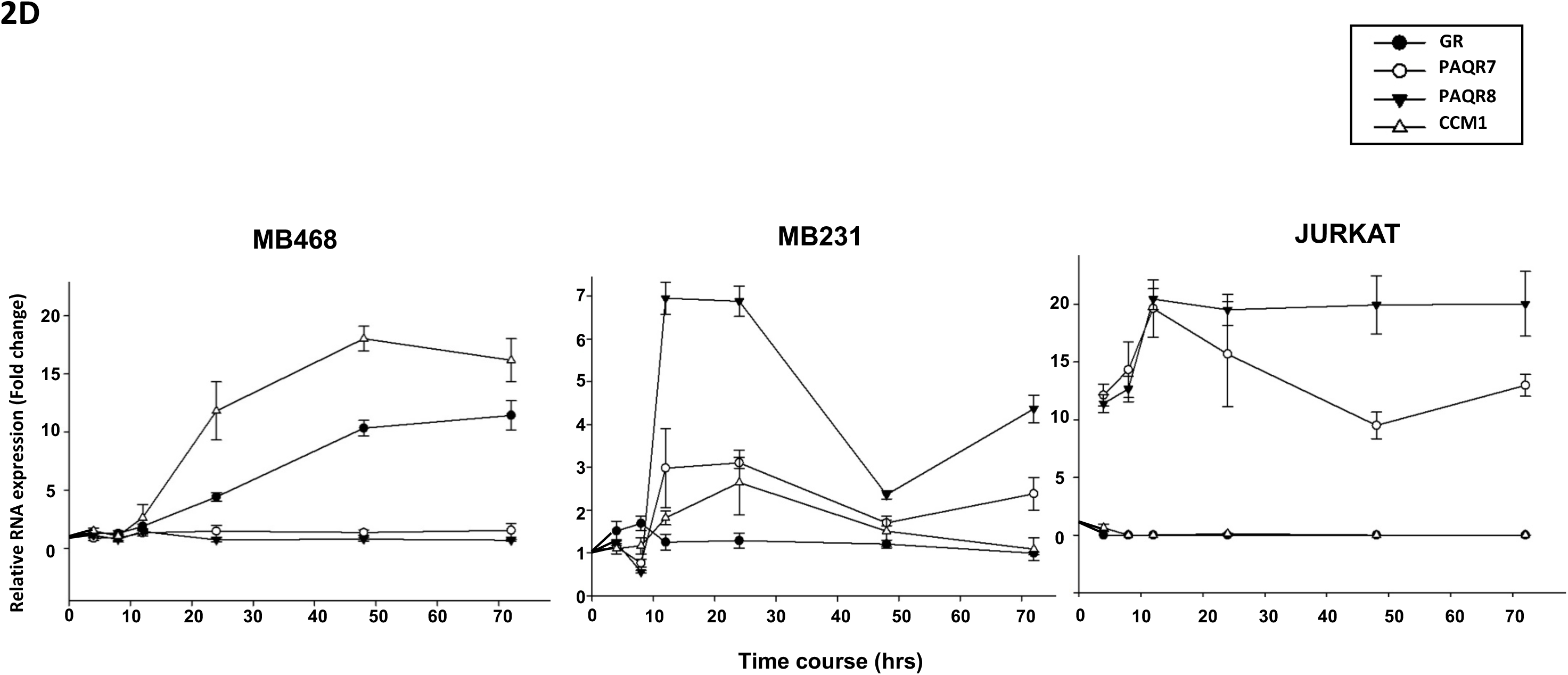
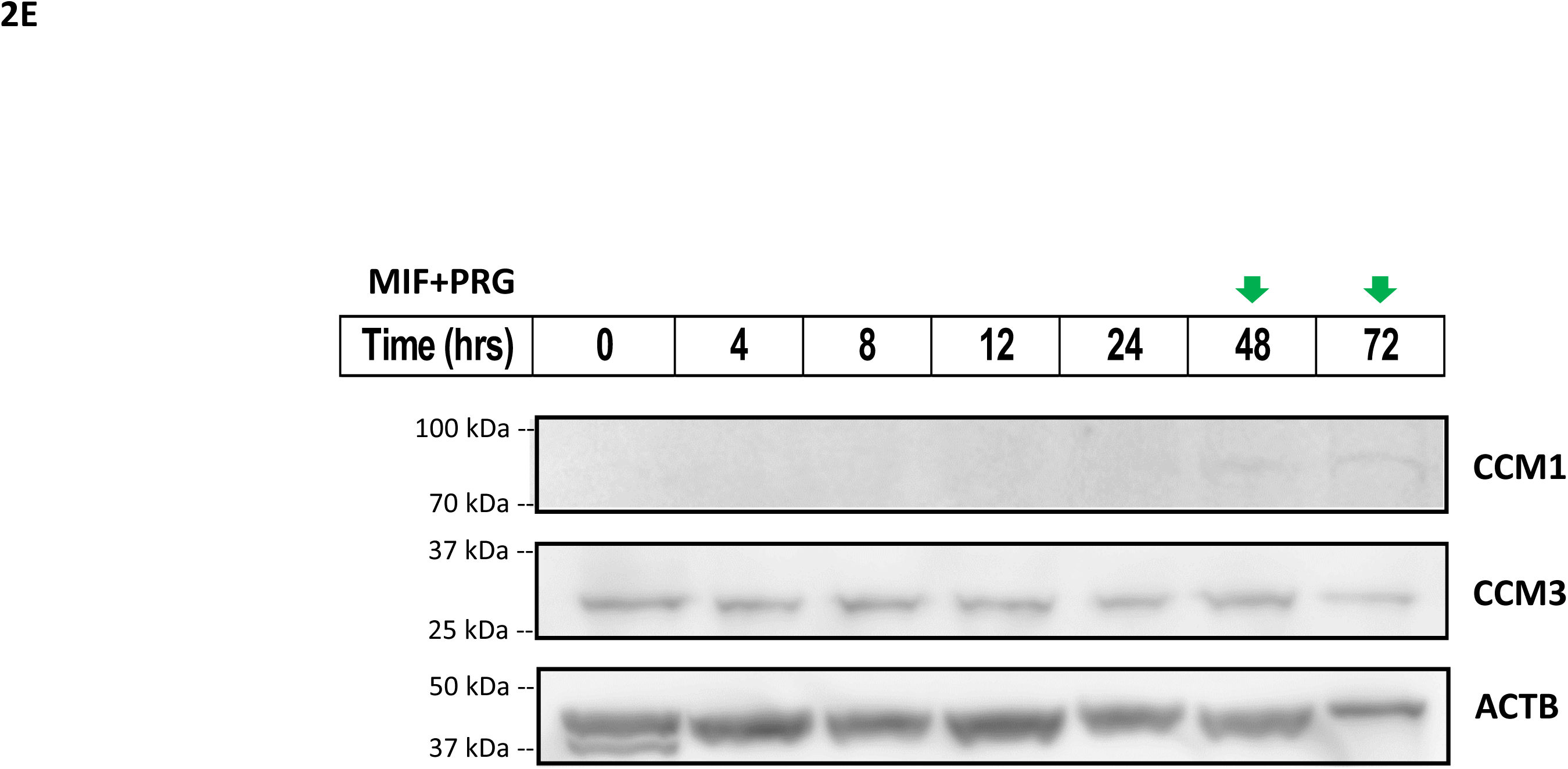
Tumorigenic assessment of two TNBC cells under combined steroid treatment. Two TNBC cells, AAW-TNBC and CAW-TNBC cells, were treated in triplicates in FBS-free medium containing vehicle control (EtOH, DMSO), Mifepristone (MIF, 20 µM), progesterone (PRG, 20 µM), or combined steroids (MIF+PRG, 20 μM each). **A**. CAW-TNBC cells displayed no major change in cell migration with PRG, compared to vehicle controls, while MIF appeared to enhance cell migration. The combined steroid effect was in-between the individual responses (Fig. 2A, upper panel). AAW-TNBC cells displayed no effect in cell migration with PRG, compared to vehicle controls, while MIF as well as the combined steroids, enhanced cell migration. **B**. Similar trends in wound healing experiments were observed. CAW-TNBC cells displayed no change in wound closure with PRG and PRG+MIF, while MIF appeared to enhance wound closure significantly, compared to vehicle controls (Fig. 2B, upper panel). AAW-TNBC cells displayed no effect in wound closure with PRG, compared to vehicle controls, while MIF as well as the combined steroids, appeared to enhance wound closure ability. **C**. In cell invasion assays, trends of decreased cell invasion for both TNBC cell lines under combined steroid actions were shown, compared with vehicle controls, without statistical significance (triplicates per experiment, n=3). **D**. Temporal RNA expression patterns of 4 key CmP network genes, *CCM1*, *GR* (glucocorticoid receptor), *mPRα* (PAQR7) and *mPRβ* (PAQR8) genes, were demonstrated in a time course analysis of the response to combined steroid treatments. Combined steroid treatment can induce RNA expression of both *CCM1* and *GR* genes in MB468 cells while no change was observed in the RNA expression of both *PAQR7/PAQR8* genes (Fig. 2D, left panel). In contrast, in both nPR(-) MB231 and Jurkat cells, a sharp induced RNA expression of both *PAQR7* and *PAQR8* genes was seen under combined steroid actions but not for *CCM1* or *GR* genes (Fig. 2D, middle and right panels), although there is a slight early induction in CCM1 gene at 24 hrs. No RNA expression was detected for both *AR* and *nPRs* genes in all three nPR(-) cells (data not shown). **E**. Sex-steroid-inducible expression of CCM1 protein in MB468 cells. The inducible expression of CCM1 protein in MB468 cells treated with combined steroid hormones was discovered in a time course analysis where green arrows demonstrate peak expression for CCM1 while no significant changes were detected for CCM3. For all panels, Relative RNA expression levels were measured through RT-qPCR (triplicates per experiment, n=3). In all bar plots, red line is the control baseline for fold change measurements (-/+).*, **, *** above bar indicates P ≤ 0.05, 0.01 or 0.001, respectively, for paired *t*-test.

### The inducible expression patterns of key factors in the CmP signaling network can be observed under combined steroid actions

The distinct RNA expression patterns between CAW-TNBC and AAW-TNBC cells are novel and interesting (Figs. 1C, 1D). We followed these phenomena temporally and found validating pieces of evidence for these opposite expression patterns. RNA expression of major *mPRs* (*PAQR7, PAQR8*) genes in type-2A cells can be drastically induced by combined steroid actions (Fig. 2D, middle and right panels), compared to no response in AAW-TNBC (Type-2B) cells (Fig. 2D, left panel), while RNA expression of *CCM1* gene in AAW-TNBC cells can be significantly induced by sex steroids (Fig. 2D, left panel), compared to little or no response in Type-2A cells (Fig. 2D, middle and right panels). These opposite temporal RNA expression data, seen among Type-2A and Type-2B cells, agree with RT-qPCR data at 48 hrs treatment (see Figs. 1C-D). Increased RNA expression levels of Glucocorticoid Receptor (*GR)* gene in AAW-TNBC cells under steroid actions is quite interesting (Fig. 2D, left panel), however, the same inducible RNA expression enhancement of *GR* gene under steroid actions was also observed in nPR(+) breast cancer cells (T47D, MCF7) with opposite cell performances (Abou-Fadel et al., 2020b), suggesting that this inducible GR might be caused by the strong binding affinity of MIF to GR, which is unrelated to tumorigenesis. Previous studies that the activation of GR signaling results in growth inhibition of various types of tumors, also suggest less relevance of activated GR in tumorigenesis (Ide et al., 2018). Finally, sex steroid-induced CCM1 expression was further confirmed at the protein level through time-course Western blots (Fig. 2E), validating our novel findings that sex steroid actions can induce distinct expression patterns of key factors (CCM1, mPRs) within the CmP signaling network between type-2A and type-2B TNBC cells.

## mPRs are nucleocytoplasmic shuttling proteins

### Temporal and spatial cellular compartmentation of key factors within the CmP signaling network

Since mPRs were initially identified as a group of cell surface and membrane-bound PRG receptors (Zhu et al., 2003b) with seven predicted membrane-spanning motifs (Smith et al., 2008; Zhu et al., 2003a; Zhu et al., 2003b), functioning as putative G-protein coupled receptors (GPCRs) (Zhu et al., 2003a), membrane-impermeable PRG-BSA conjugate techniques (Zheng et al., 1996) have been widely used to measure PRG-mPRs signaling with a sole focus on its non-genomic actions in various organisms and cell types (Josefsberg Ben-Yehoshua et al., 2007; Kasubuchi et al., 2017; Pang et al., 2015; Tischkau and Ramirez, 1993; Xie et al., 2015). It has been reported that there is very low expression of PAQR7 in MB231 cells (Xie et al., 2015; Xie et al., 2012), therefore we performed cellular localization of PAQR8 along with other key factors (CCM1/3) in the CmP network in both CAW/AAW-TNBC cells. Under our observations, initially, CCM1 protein is equally distributed in the nucleus in both TNBC cells, however, after combined steroid actions both nuclear and cytoplasmic increased expression of CCM1 protein is visualized in AAW-TNBC cells (Fig. 3A1, left panel), while CCM1 protein expression not only increases as well but starts to concentrate along the nuclear membrane (peri-nucleus) in CAW-TNBC cells (Fig. 3A1, right panel). In contrast, CCM3 protein is distributed dominantly in the nucleus initially and combined steroid-enhanced expression of CCM3 proteins are visualized more towards the cytosol (more dramatic in AAW-TNBC cells) (Fig. 3A2). Intriguingly, PAQR8 is clearly localized within the nucleus, and remains unchanged under combined steroid actions in AAW-TNBC cells (Fig. 3A3, left panel), while PAQR8 protein is dominantly distributed within the nucleus (likely associated with the nucleolus), and becomes more evenly distributed in both nucleus and cytosol after combined steroid actions in CAW-TNBC cells (Fig. 3A3, right panel). It was not a total surprise for us to observe nuclear localization of PAQR8 in TNBC cells, since we have observed a similar phenomenon in Luminal-A T47D cells (Abou-Fadel et al., 2020b). There have been contradicting reports of cellular localization of mPRs in either the cytoplasmic membrane or cytoplasmic compartmentations (mainly rough ERs) in several cell types, including MB231 cells (Krietsch et al., 2006; Zhu et al., 2003a), while cellular compartmentations of mPRs on MB468 cells have yet to be investigated. Our data clearly show that PAQR8 is capable of nuclear sub-compartmentation in both TNBC cells. Furthermore, we also observed nuclear localization of PAQR5 (mPRγ) and PAQR7 (mPRα) in both TNBC cell lines utilizing IHC techniques (Suppl. Fig 3A). Together, these data support the notion that mPRs are nuclear proteins in TNBC cells.

Next, we examined the temporal expression patterns under combined steroid actions, demonstrating steroid-induced expression of CCM1 again in AAW-TNBC cells (Fig. 3B1), in line with our previous data (Figs. 1D, 2D, 2E). Interestingly, there is an early surge (non-genomic) expression of CCM3 protein in AAW-TNBC cells (Fig. 3B2), which is unrelated to the genomic induced expression of CCM3 (increased RNA expression at earlier timepoints, Fig. 2D), and opposite to the data observed in luminal A T47D breast cancer cells (Abou-Fadel et al., 2020b). The underlined mechanism of this fluctuation of CCM3 protein in AAW-TNBC cells is still unknown. A visual subtle decrease of PAQR8 in AAW-TNBC cells under steroid actions was also observed (Fig. 3B3). Interestingly, steroid-induced expression of CCM1 was observed again in CAW-TNBC cells (Fig. 3C1), collaborating with our previous data (Fig. 2D). Likewise, steroid-induced expression of CCM3 (Fig. 3C2) and PAQR8 in CAW-TNBC cells (Fig. 3C3) were also visualized, in line with our previous data (Figs. 1D, 2D). The major discovery from these observations is the temporal and spatial regulation of protein expression of key players of the CmP network under steroid actions in both TNBC cells.

### Cytoplasmic-nuclear trafficking of key players of the CmP network

Another novel finding associated with our data is the cytoplasmic-nuclear trafficking of mPRs. Utilizing a larger sample size, we carefully examined the temporally subcellular localization of key factors (CCM1/3, PAQR8) in the CmP network under steroid actions in both TNBC cells and not only found PAQR8 inside the nucleus (Fig. 3B-3, 3C-3) but also found the temporally dynamic expression patterns of PAQR8 in the nucleus and cytosol in both TNBC cells (Fig. 3D, upper panels). Cytoplasmic-nuclear trafficking of CCM1 has been well studied by our group (Liu et al., 2010; Liu et al., 2011; Zhang et al., 2004; Zhang et al., 2005; Zhang et al., 2008; Zhang et al., 2011), therefore, we use CCM1 as a positive control to examine the possible cytoplasmic-nuclear trafficking of PAQR8 by measuring the expressional ratio of nucleus/cytosol along the time-course. We found that both CCM1 and CCM3 proteins are predominantly in the nucleus in AAW-TNBC cells (Fig. 3D, middle and lower left panels), while CCM1 and CCM3 show distinct nucleo-cytoplasmic shuttling capabilities with a more equal subcellular compartmentation in CAW-TNBC cells (Fig. 3D, middle and lower right panels). Intriguingly, alternative fluctuations of the expressional ratio of nucleus/cytosol along the time-course are only seen in both CCM1 and PAQR8 in CAW-TNBC cells with the same temporal and spatial trends observed after 4 hrs, indicating the coordinately synchronized cytoplasmic-nuclear trafficking activities of endogenous CCM1 and PAQR8 proteins only in CAW-TNBC cells under steroid actions (Fig. 3D, middle and upper right panels). The observed phenomena suggest that PAQR8 nucleocytoplasmic shuttling might be CCM1-dependent and recent data showed that many proteins constantly shuttle between the cytoplasm and the nucleus without an obvious *bona fide* classic NLS and/or NES signaling site (Fu et al., 2018), however, our bioinformatics simulation data suggested the possible existence of one NLS and two NES signaling sites within PAQR8 (Suppl. Fig. 3B), further strengthening our observations of nuclear-cytoplasmic shuttling of PAQR8 in both TNBC cells.

**Fig. 3:**
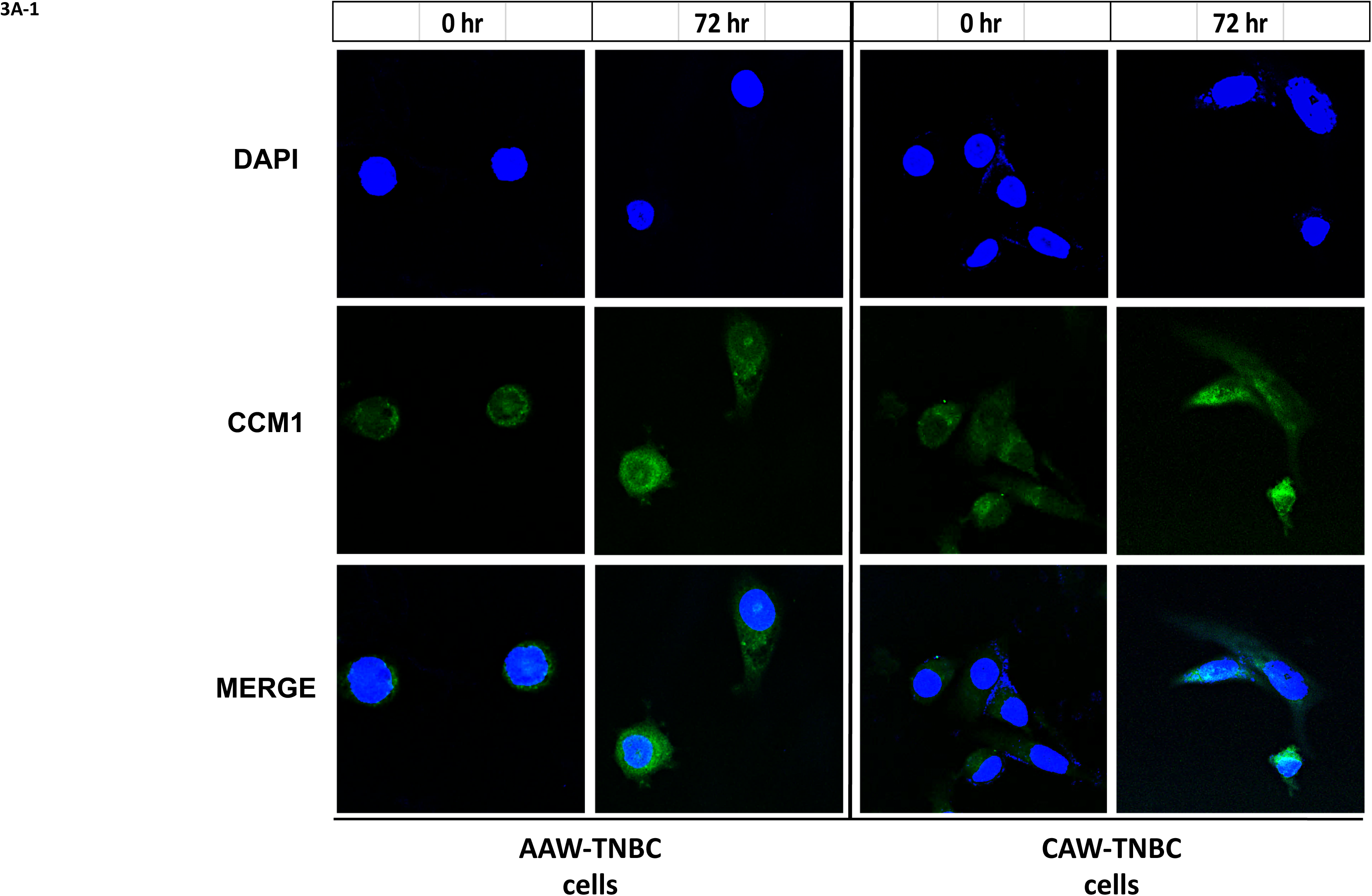
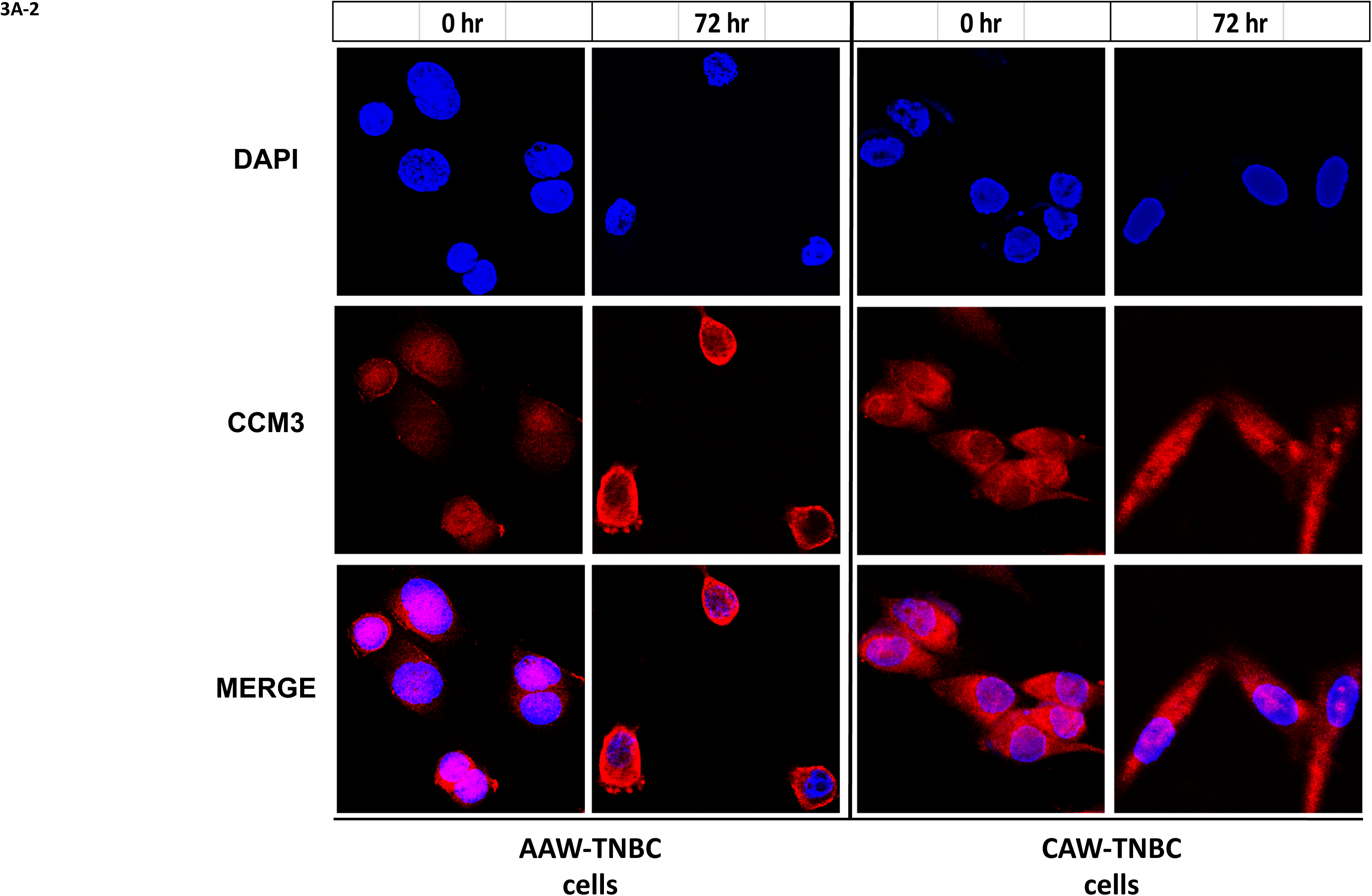
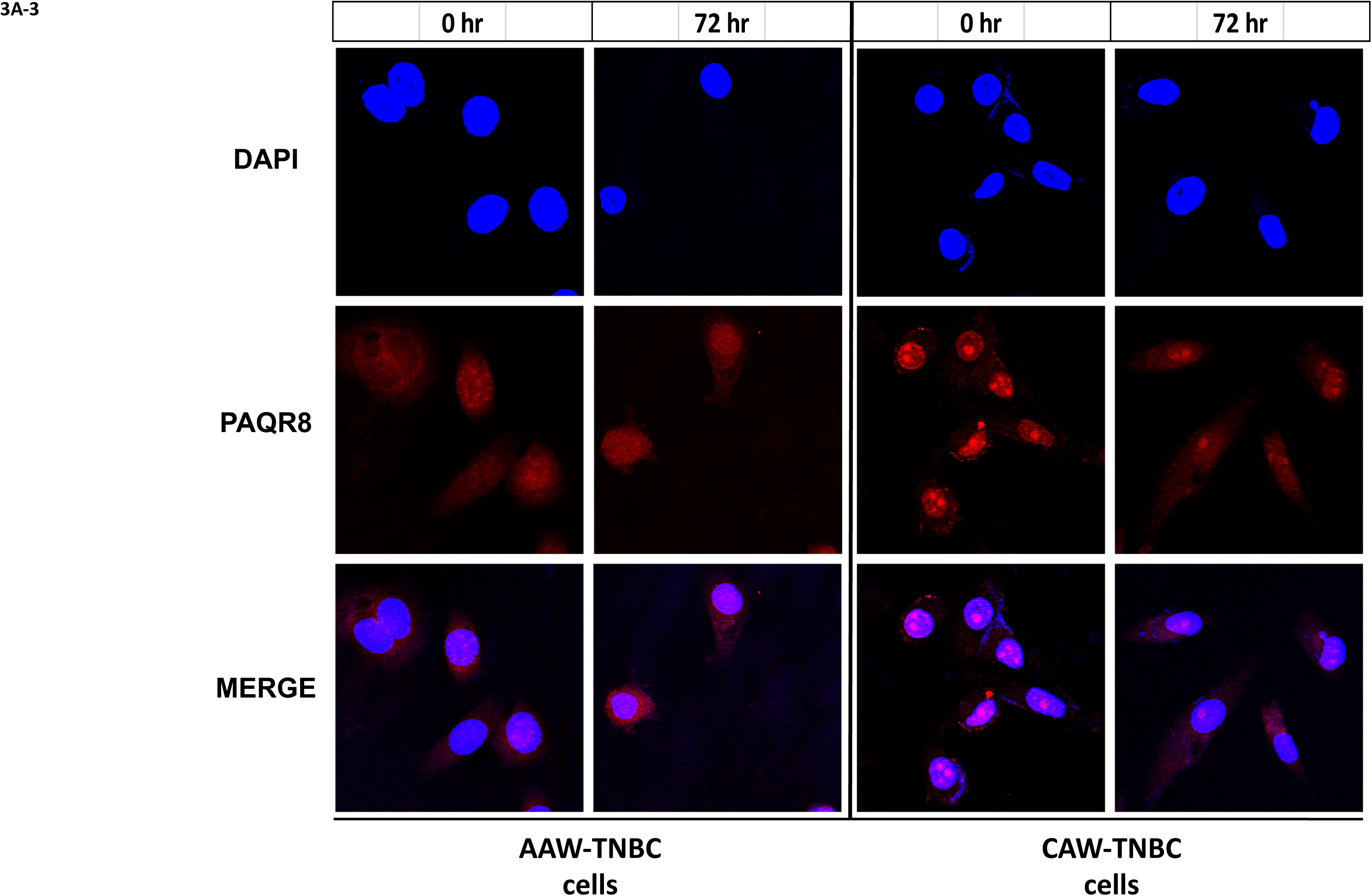
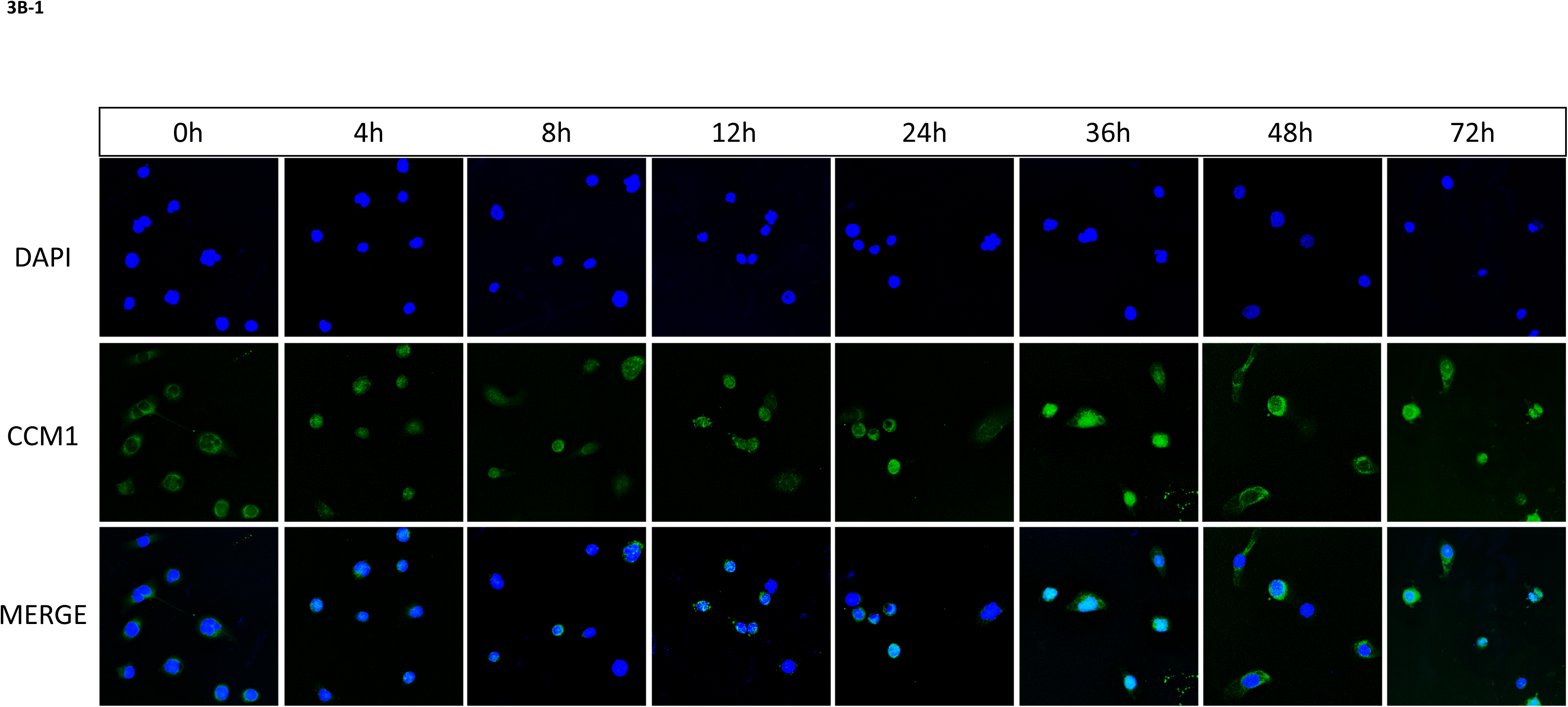
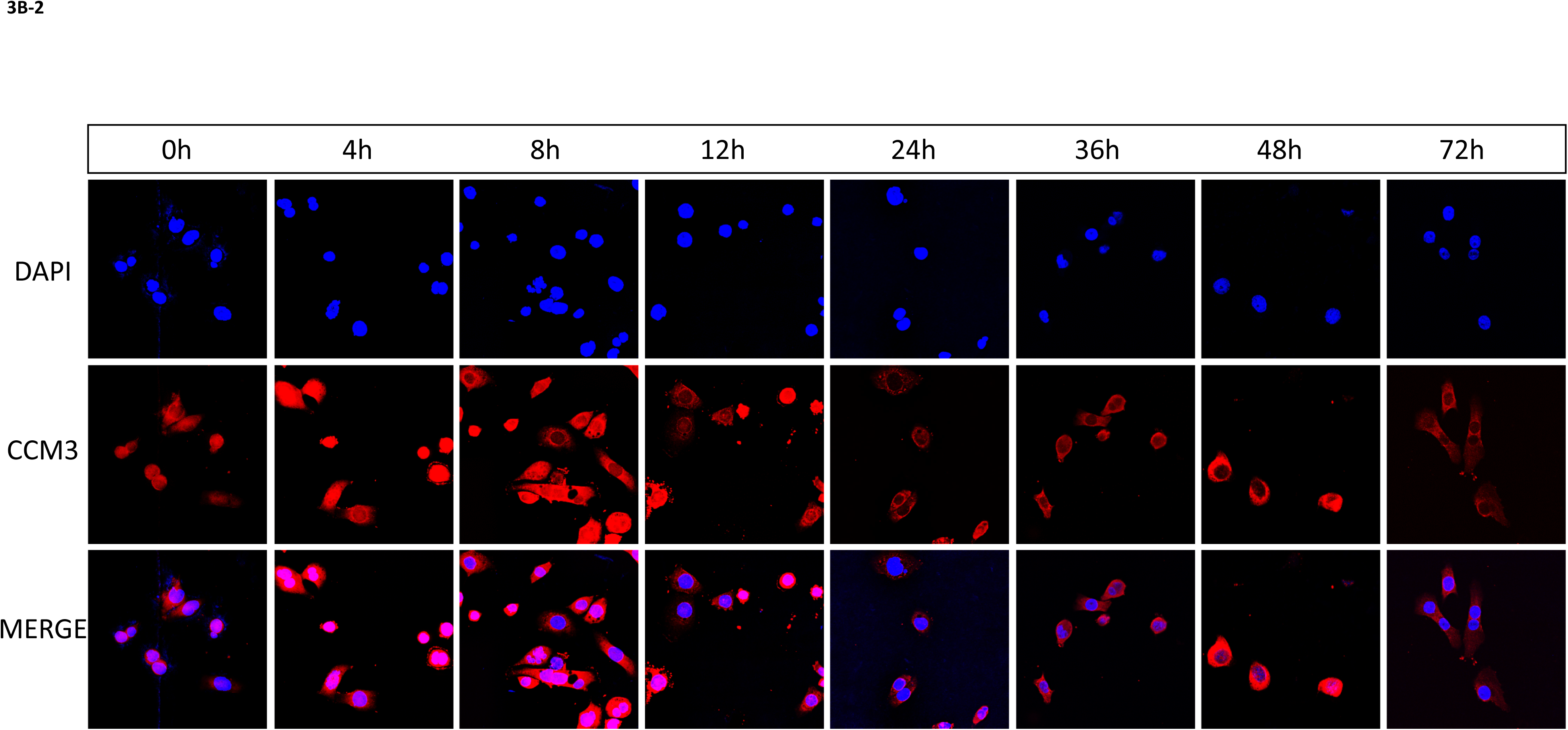
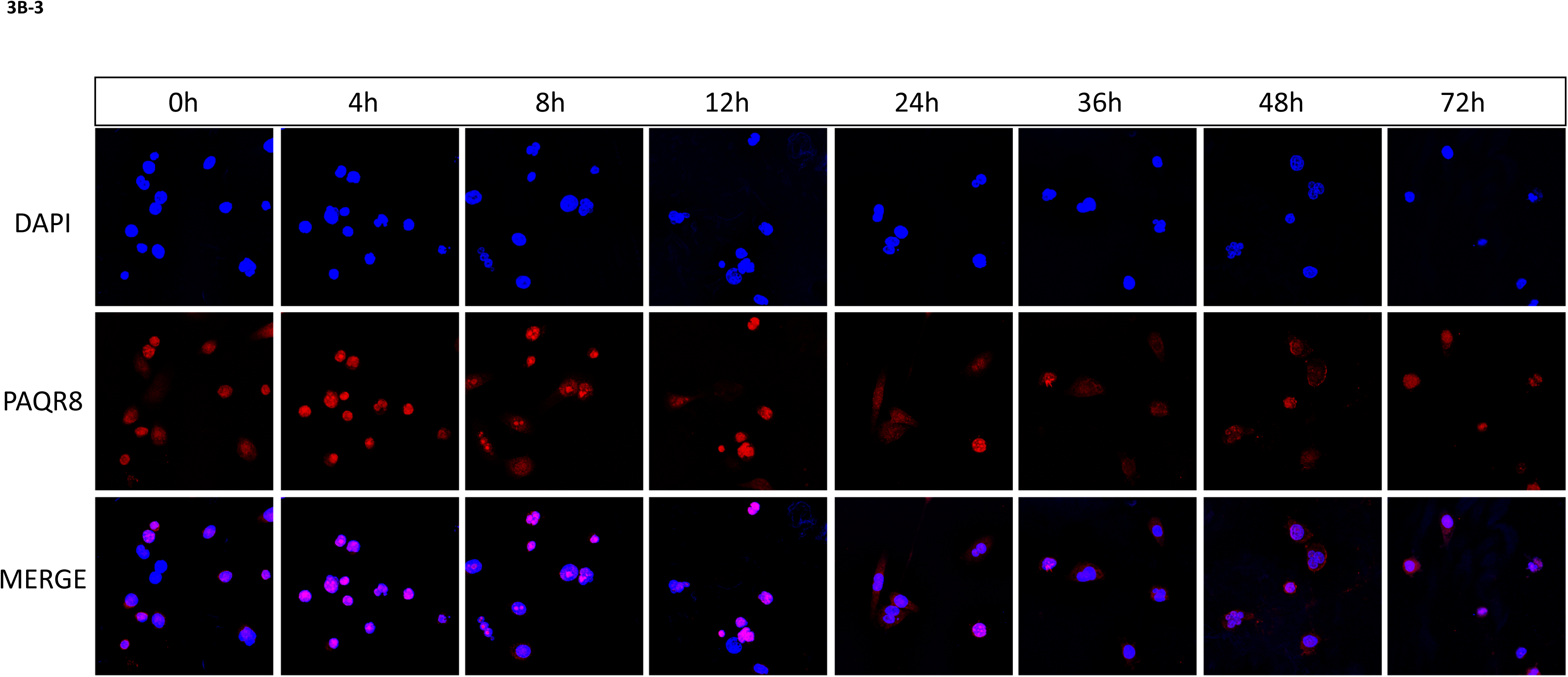
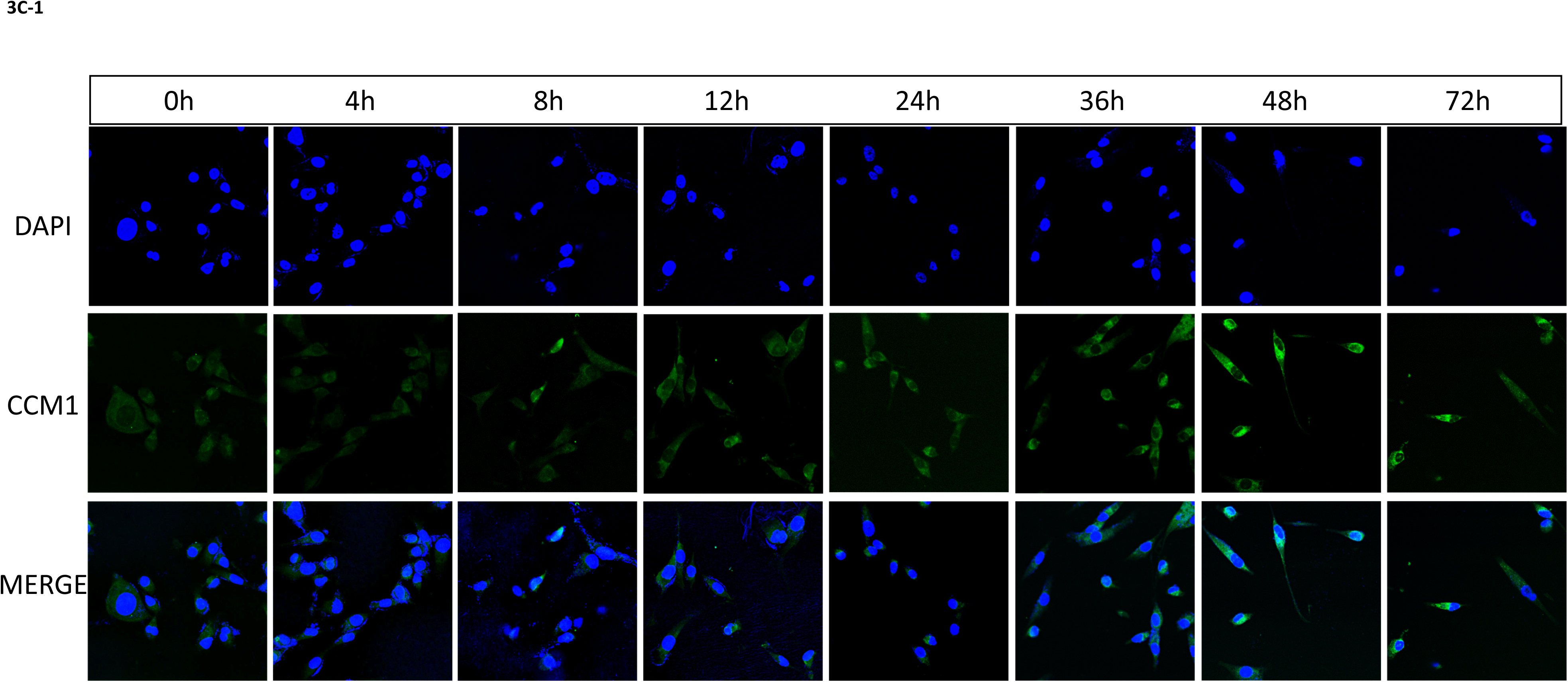
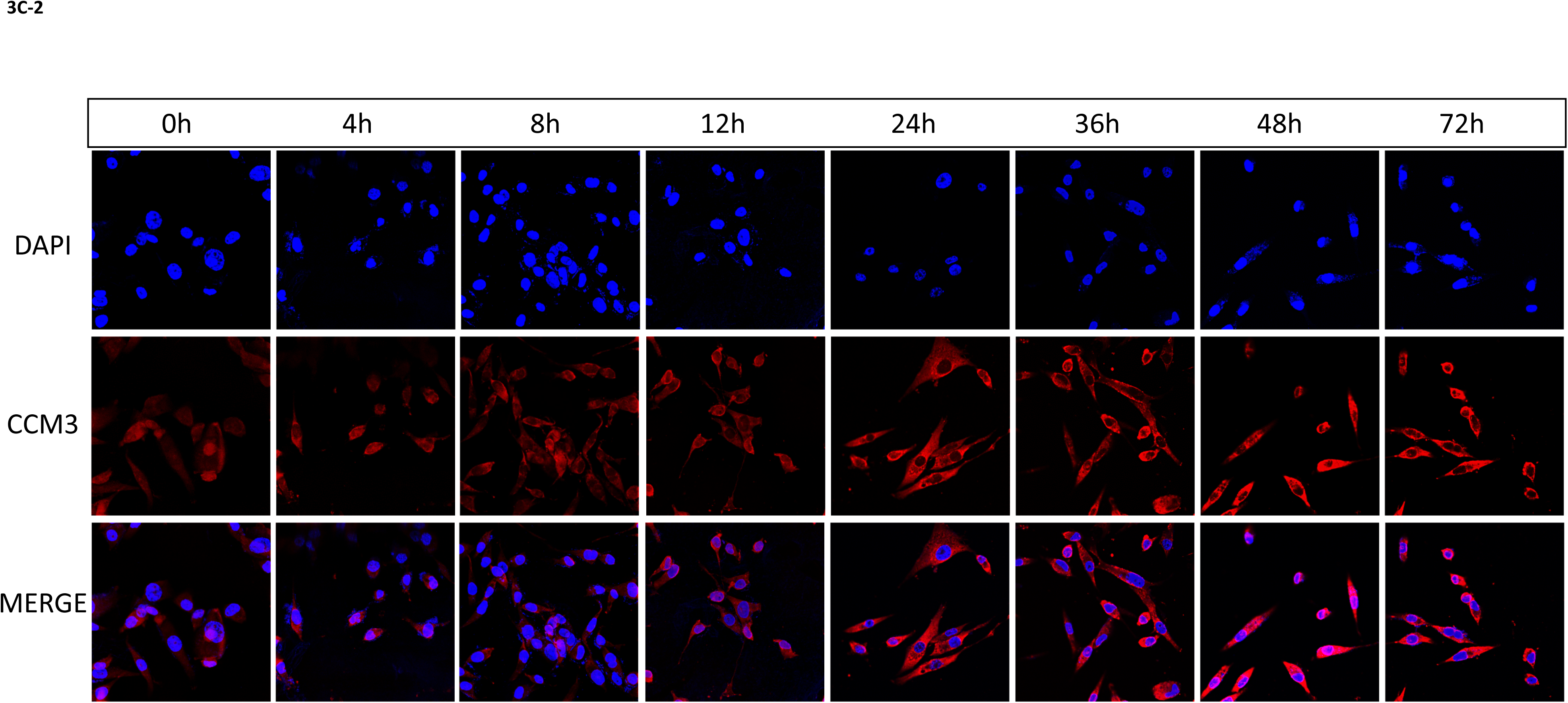
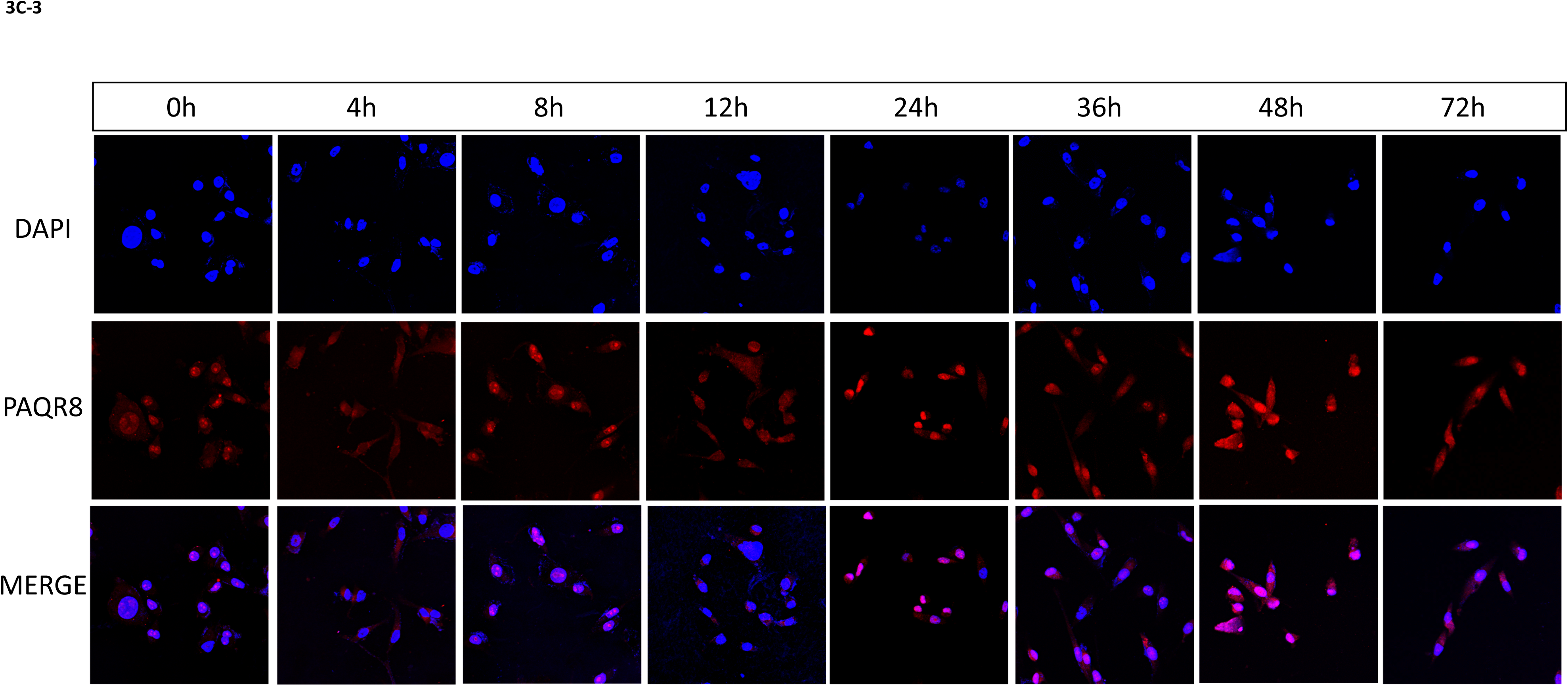
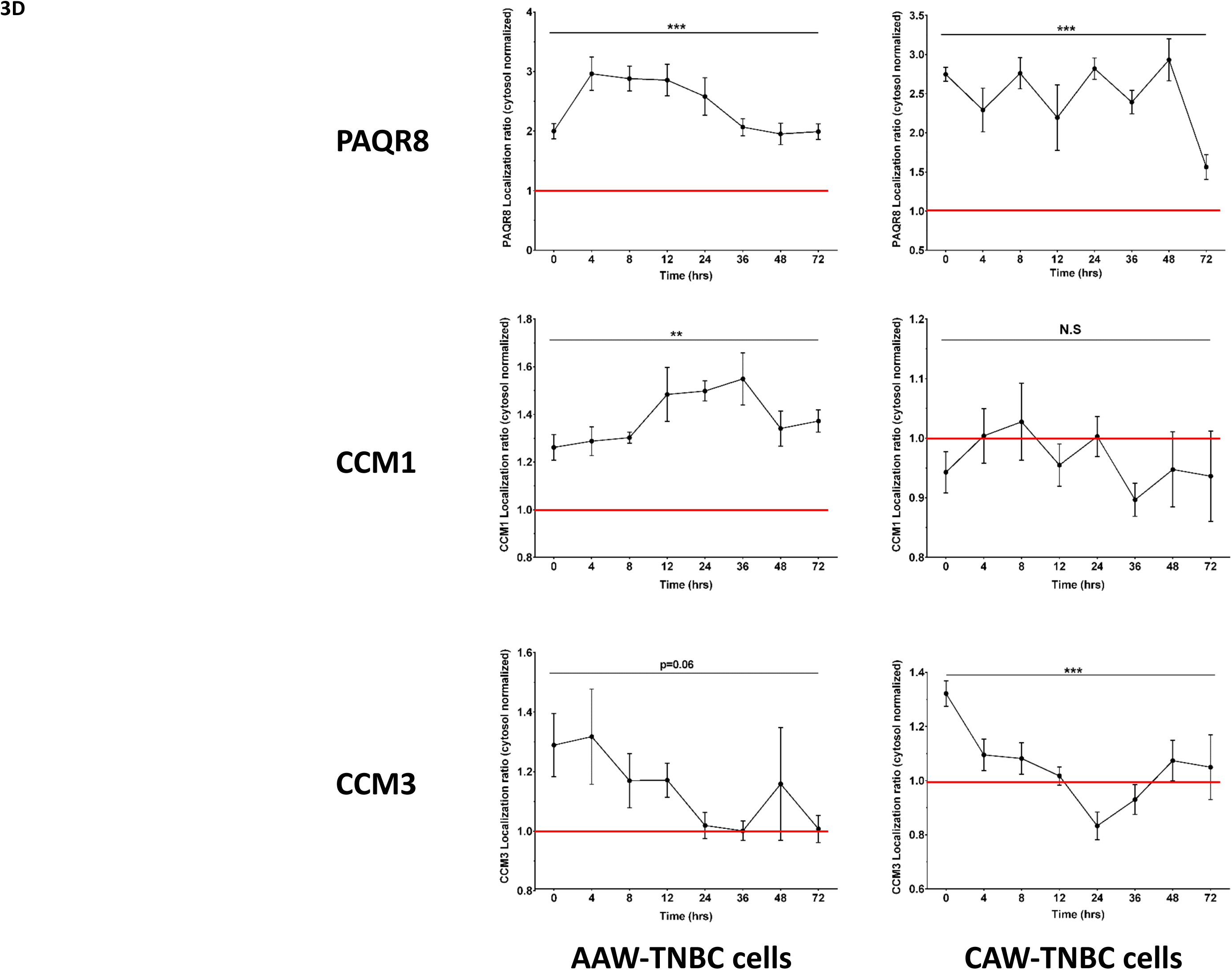
Modulation of key factors of the CmP network in two TNBC cells under combined steroid actions. CAW-TNBC and AAW-TNBC cells were treated with combined steroids (MIF+PRG, 20µM each) for 72 hrs. **A-1**. Relative expression of CCM1 protein. IF approaches revealed significant modulation in the relative intensity of CCM1 staining between TNBC cells treated with MIF+PRG. Both CAW-TNBC and AAW-TNBC cells are capable of altering localization of CCM1 proteins in and out of the nucleus and demonstrated increased relative CCM1 expression inside the nucleus at 72 hrs, although the majority of staining resides in the peri-nuclear area (cytosol). The relative expression of CCM1 is also greater at 72 hrs compared to 0 hrs. **A-2**. Relative expression of CCM3 protein. IF approaches revealed modulation in the relative intensity of CCM3 staining in AAW-TNBC cells treated with MIF+PRG. Staining revealed significant changes in cellular localization of CCM3 in both TNBC cells, but only showed a significant increase in CCM3 intensity between 0 hrs and 72 hrs for AAW-TNBC cells. **A-3**. Relative expression of PAQR8 protein. IF approaches revealed significant modulation in the localization of PAQR8 staining between representative CAW-TNBC cells treated with MIF+PRG. Interestingly, CAW-TNBC cells showed more PAQR8 staining inside the nucleus at 0 hrs followed by altering localization of PAQR8 back to the cytosol at 72 hrs. AAW-TNBC cells did not show much change in localization or expression of PAQR8 at 72 hrs compared to 0 hrs. **B-1**. Temporal modulation of CCM1 protein in AAW-TNBC cells. AAW-TNBC cells were treated with combined steroids for 0-72 hrs. Relative expression of total CCM1 proteins in AAW-TNBC cells was performed. IF approaches revealed significant modulation in the relative intensity of CCM1 staining between representative TNBC cells treated with MIF+PRG over the specified time course. Relative CCM1 expression demonstrated an increased intensity observed at 36hrs throughout the remainder of the time course. **B-2.** Temporal modulation of CCM3 protein in AAW-TNBC cells. Relative expression of total CCM3 proteins in AAW-TNBC cells was performed. IF approaches revealed modulation in the relative intensity of CCM3 staining between AAW-TNBC cells treated with MIF+PRG over the specified time course. Overall quantification demonstrated significant modulation of relative CCM3 expression with an increased intensity observed at 4hrs, followed by a steady decrease back to normal expression after 24h. **B-3.** Temporal modulation of PAQR8 protein in AAW-TNBC cells. Relative expression of total PAQR8 proteins in AAW-TNBC cells was performed. IF approaches revealed modulation in the relative intensity of PAQR8 staining between AAW-TNBC cells treated with MIF+PRG over the specified time course. Relative PAQR8 expression demonstrated an increased intensity observed at 4hrs, followed by a steady decrease back down to normal expression by 36h. **C-1**. Temporal modulation of CCM1 protein in CAW-TNBC cells. CAW-TNBC cells were treated with combined steroids for 0-72 hrs. Relative expression of total CCM1 proteins in CAW-TNBC cells was performed. IF approaches revealed significant modulation in the relative intensity of CCM1 staining between CAW-TNBC cells treated with MIF+PRG over the specified time course. Interestingly, increased expression in CCM1 was observed after 8 hours throughout the remainder of the time course. **C-2.** Temporal modulation of CCM3 protein in CAW-TNBC cells. Relative expression of total CCM3 proteins in CAW-TNBC cells was performed. IF approaches revealed modulation in the relative intensity of CCM3 staining between CAW-TNBC cells treated with MIF+PRG over the specified time course. Overall quantification demonstrated significant modulation of relative CCM3 expression with an increased intensity observed at 12hrs throughout the remainder of the time course. **C-3.** Temporal modulation of PAQR8 protein in CAW-TNBC cells. Relative expression of total PAQR8 proteins in CAW-TNBC cells was performed. IF approaches revealed modulation in quantification in the relative intensity of PAQR8 staining between CAW-TNBC cells treated with MIF+PRG over the specified time course. Relative PAQR8 expression demonstrated an increased intensity observed at 24hrs, followed by a steady decrease back down to normal expression by 72hrs. **3D.** Sub-cellular localization of key factors of the CmP network in two TNBC cells under combined steroid treatment. Using NIKON elements analysis tools, localization of PAQR8 was quantified using binary operations to identify the relative ratio of expressed PAQR8 that was localized inside the nucleus (overlaps of DAPI/mCHERRY) compared to expressed PAQR8 that was localized in the cytosol (unique mCHERRY without overlapping DAPI signal). Interestingly, the increased expression observed in total PAQR8 in both CAW-TNBC and AAW-TNBC cells appears to localize specifically inside the nucleus, and even when levels have decreased back down to those levels observed at 0h, the ratio decreases back towards the cytosol, but is still predominantly located in the nucleus (top panels). Localization of CCM1 was quantified using binary operations to identify the relative ratio of expressed CCM1 that was localized inside the nucleus (overlaps of DAPI/GFP) compared to expressed CCM1 that was localized in the cytosol (unique GFP without overlapping DAPI signal). Interestingly, increased expression in nuclear CCM1 in AAW-TNBC cells after 8 hours was observed and the expression appears to still be predominantly located in the nucleus throughout the time course while the opposite trend was observed for CCM1 expression in CAW-TNBC cells, where CCM1 is localized in the cytosol for most timepoints (middle panels). Localization of CCM3 was performed using binary operations to identify the relative ratio of expressed CCM3 that was localized inside the nucleus (overlaps of DAPI/mCHERRY) compared to expressed CCM3 that was localized in the cytosol (unique mCHERRY without overlapping DAPI signal). Interestingly, expression of CCM3 in AAW-TNBC cells appears to localize specifically inside the nucleus at the earlier time points (0-12 hrs), and then localizes back into the cytosol closer to a 1:1 ratio, (with the exception of 48 hrs) (bottom left panel). Expression of CCM3 in CAW-TNBC cells appears to localize specifically inside the nucleus at the earlier time points (0-12 hrs), shuttles back into the cytosol at 24-36 hrs, and then shuttles back into the nucleus for the remainder of the experiment. Red line indicates a 1:1 ratio of proteins in nucleus: cytosol. CCM1 was quantified through ROI intensities using wavelength channels 488nm. PAQR8 and CCM3 was quantified through ROI intensities using wavelength channels 555nm. Data were acquired from 8 sets of independent images and normalized against its respective internal controls using 408nm wavelength for DAPI (nuclear) and background staining subtraction. For each replicate, Region of Interest (*ROI*) i*ntensities* were automatically quantified (over 1000 times/per area). In all bar plots, ** and *** above any bar graph indicate P ≤ 0.01 and P ≤ 0.001, respectively using One-way ANOVA (n=8).

## Our novel findings are further validated by Omics data

### Dissecting the CmP network in AAW-derived TNBC cells using omics approaches

To define key altered pathways, potential partners involved in the CmP signaling network, and potential biomarkers, we examined the expression patterns of AAW-TNBC cells under combined steroid actions, at both the transcriptional and translational levels using high throughput omic approaches, including RNAseq and LC-MS/MS. Since RNA expression of *CCM1* gene in AAW-TNBC cells can be significantly induced by sex steroids (Fig. 2D, left panel), compared to no or little response in Type 2A (MB231/Jurkat) cells (Fig. 2D, middle and right panels), we filtered out our RNAseq and proteomics data for AAW-TNBC cells by removing similarly altered Differentially Expressed Genes/Proteins (DEGs/DEPs) shared between Type 2B and Type 2A cells to evaluate AAW-TNBC specific DEGs/DEPs.

### Key signaling cascades identified within the CmP network in AAW-TNBC cells using RNAseq

Among identified DEGs (Suppl. Table. 4A), we were able to visualize hierarchical clustering (Fig. 4A, top panel), and found similar temporal patterns in both intersection and union of DEGs (Figs. 4A, bottom panel). Identification of up and down-regulated genes at each time point revealed that combined steroid actions have an overall stronger up-regulation on AAW-TNBC specific DEGs (Fig. 4B, top panel). Further stratification demonstrated 1300+ genes temporally up-regulated and 700+ genes temporally down-regulated under combined steroid actions yielding a great foundation of potential biomarkers to distinguish type-2B (AAW-derived) from other TNBCs (Fig. 4B, bottom panel). Pathways functional enrichment revealed several major altered pathways including the most affected signaling pathway which was the Gonadotropin-releasing hormone (GnRH) signaling pathway (Fig. 4C, top panel), responsible for regulating mammalian reproduction, including production of hormones in both men and women (Perrett and McArdle, 2013). Amplification of EGFR members is one of the most common genetic alterations associated with breast cancer (Filardo, 2002), and interestingly, EGF receptor signaling pathways was one of the top 10 affected pathways in AAW-TNBC cells under combined steroid actions (Fig. 4C, top panel), reaffirming the role of key players in the CmP network in AAW-TNBC progression. The other top signaling pathways altered in AAW-TNBC cells under steroid actions included Inflammation mediated by chemokines and cytokines, several angiogenesis-related pathways as well as heterotrimeric G-protein pathways (Fig. 4C). Together, these results further validate the crosstalk between the CSC and mPRs in the CmP signaling network involved in AAW-TNBC tumorigenesis. Finally, several key signaling cascades in tumorigenesis were frequently observed to be altered in AAW-TNBC cells under combined steroid actions, including apoptosis, p53, MAPK, WNT, RAS, and hormone signaling pathways, further validating involvement of the CmP network in AAW-TNBC progression (Fig. 4C)(Suppl. Figs 4A-E).

**Figure 4:**
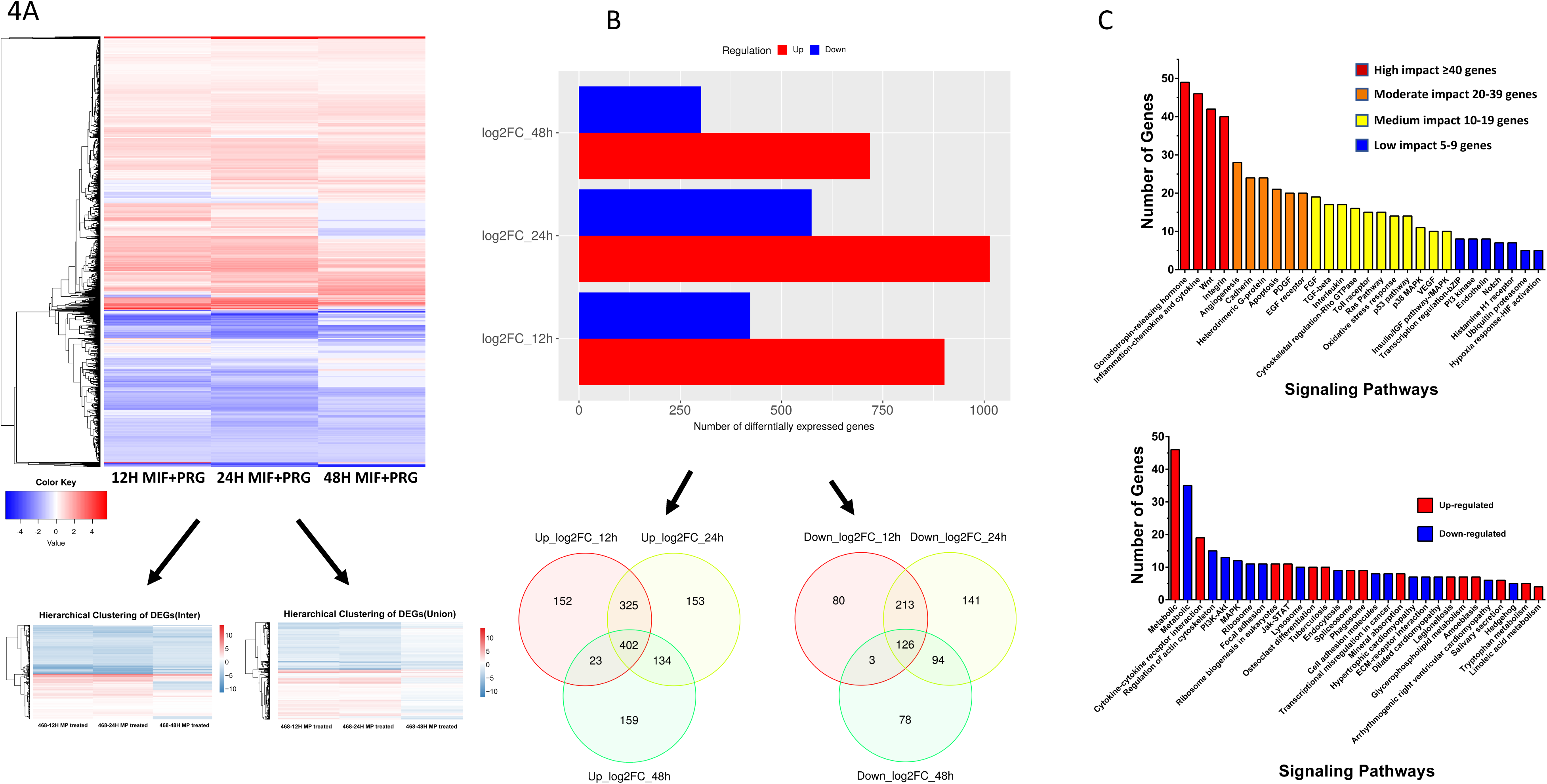
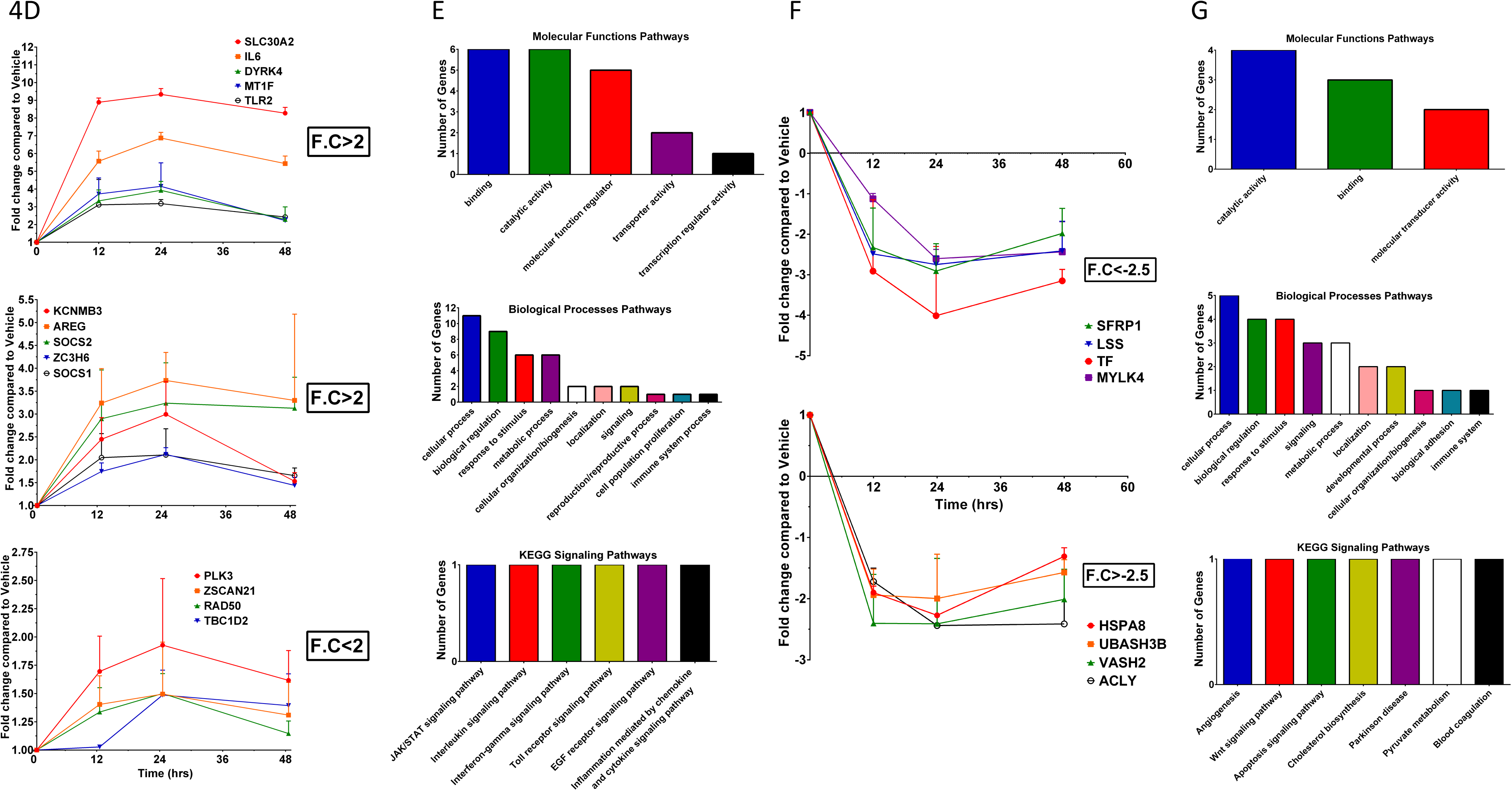
Differentially Expressed Genes (DEGs) among AAW-TNBC cells utilizing high-throughput RNA sequencing (RNAseq): AAW-TNBC cells were treated with MIF+PRG (20µM each) for 12, 24 and 48 hrs. Extracted significant DEGs were further filtered using corresponding MB231/Jurkat cells to obtain DEGs specific to AAW-TNBC cells. **A)** Cluster software and Euclidean distance matrixes were used for the hierarchical clustering analysis of the expressed genes (RNA) and sample program at the same time to generate the overall Heatmap of hierarchical clustering (top panel); intersection of DEGs *(bottom left panel),* or union of DEGs *(Bottom right panel)* of expression clustering scheme are also displayed; x axis represents each comparing sample and Y axis represents DEGs. **B)** Bioinformatics analysis was performed to identify significant DEGs, compared to vehicle controls *(Top panel)*. Significant DEGs were further stratified into independent Venn diagrams for both up and down regulated genes to identify overlaps between timepoints *(Bottom panels)***. C)** Pathway functional enrichment results for MIF+PRG treated cells compared to their respective vehicle controls were compiled using molecular functions, biological processes and signaling pathway data retrieved from the PANTHER (Protein ANalysis THrough Evolutionary Relationships) classification system (top panel). Additionally, signaling pathways were further stratified to identify up/down regulation within each signaling pathway using iDEP (Integrated Differential Expression and Pathway Analysis) program (bottom panel). For upper panel C, coloring indicates the potential impact on each signaling pathway (high (≥40 genes): red, moderate (20-39 genes): orange, medium (10-19 genes): yellow, low (5-9 genes): blue. For panels A, B and lower C, coloring indicates the log2 transformed fold change (high: red, low: blue). **D)** To delineate cellular partners sharing the same/reverse inducible expression patterns as CCM1 using high-throughput RNAseq. Transcriptomics analysis was temporally performed to identify consistently up-regulated genes (statistically significant), compared to vehicle controls, with peak Log2FC categories greater than 2.0 *(upper and middle panels) and* peak Log2FC categories less than 2.0 *(lower panel)*. **E)** Pathway functional enrichment results for corresponding genes shown in panel D were compiled using molecular functions, biological processes and signaling pathway data retrieved from the PANTHER classification system. **F)** Transcriptomics analysis was temporally performed to identify consistently down-regulated genes (statistically significant), compared to vehicle controls, with peak Log2FC categories less than −2.5 *(upper panel) and* peak Log2FC categories greater than −2.5 (lower panel). **G)** Pathway functional enrichment results for corresponding genes shown in panel F were compiled using molecular functions, biological processes and signaling pathway data retrieved from the PANTHER classification system. For all functional enrichment graphs, X axis represents the terms of pathways while y axis represents the number of identified DEGs involved in each pathway. For all panels, each timepoint was run with biological duplicates and technical triplicates for treated and vehicle samples.

To delineate cellular partners sharing the same inducible expression patterns as CCM1 (Fig. 2D), transcriptomics analysis was temporally performed to identify consistently up-regulated genes (Suppl. Table. 4B) (statistically significant), compared to vehicle controls (Fig. 4D). We identified 14 genes that were consistently up-regulated under sterol actions (Fig. 4D), indicating that steroid hormone treatment can induce RNA expression for a subset of genes, identical to the expression patterns seen for *CCM1* (Fig. 2D, left panel) in AAW-TNBC cells. Among the identified 14 AAW-TNBC specific DEGs, was *IL6*, a well-known player in key cancer pathways including folate metabolism and transcriptional dysregulation in cancer (Fig. 4D, upper panel). *AREG* was also discovered to be up-regulated (Fig. 4D, middle panel) and AREG-mediated activation of EGFR results in up-regulation of fibronectin, which is known to be a mediator of invasive capacity via interaction with integrin β1 in TNBCs (Kappler et al., 2015). *Rad50*, a component of the MRE11 complex, has an essential role in maintaining genomic integrity and preventing cells from malignancy (Heikkinen et al., 2006) (Fig. 4D, bottom panel). *PLK3* has been suggested as a novel independent prognostic marker in breast cancer due to PLK isoform overexpression being associated with disease progression (Weichert et al., 2005)(Fig. 4D, lower panel). Alterations in several key pathways included cell population proliferation, reproduction, and immune system processes (Fig. 4E). Additionally, these results validated altered pathways seen in the global RNAseq profiling of AAW-TNBC cells under combined steroid actions including JAK/STAT, Interleukin, EGFR, and inflammation-mediated by chemokine and cytokine signaling pathways (Figs. 4C, 4E). Next, transcriptomics analysis was temporally performed to identify consistently down-regulated genes (statistically significant), compared to vehicle controls (Fig. 4F, Suppl. Table. 4B). We identified 8 AAW-TNBC specific genes that were consistently down-regulated under sterol actions (Fig. 4F), including *SFRP1*, which deregulates activation of the WNT pathway when silenced and is an early event in breast tumorigenesis (Klopocki et al., 2004). Interestingly, *LSS* was also down-regulated, which is a known player in catalyzing the first step in the biosynthesis of steroid hormones. Additional altered signaling pathways included angiogenesis, WNT, and apoptosis in AAW-TNBC cells under steroid actions (Fig. 4G).

### Key signaling cascades within the CmP network in AAW-TNBC cells using LC-MS/MS approaches

Through Euclidean mapping of DEPs (Suppl. Table 5A), we were able to visualize proteomic regulation (Fig. 5A and B, top panels), and upon identifying up and down-regulated genes temporally, revealed that there is a greater propensity of upregulated proteins at 24 and 72 hrs when analyzed using *t*-test (Fig. 5A, middle panel) while there is a greater propensity of upregulated proteins at all three timepoints compared to down-regulated when analyzed using ANOVA methods (Fig. 5B, middle panel). Hormone-treated AAW-TNBC cells showed a total of 66 upregulated proteins using *t-test* analysis (Fig. 5A, bottom left panel) while 342 proteins were identified using ANOVA methods (Fig. 5B, bottom left panel). In contrast, 70 proteins were observed down-regulated using *t-test* analysis (Fig. 5A, bottom right panel) and 27 proteins using ANOVA statistics (Fig. 5B, bottom right panel). In complementation to potential DEG biomarkers (Fig. 4) identified, we now have an additional index of potential AAW-TNBC specific DEP biomarkers that may help limit the scope of commonly shared biomarkers in different sub-types of TNBCs. When proteins were functionally enriched, we noted that some pathways overlaid with the RNA enrichment, reinforcing the concept that these may play potential key roles in hormone signaling and factors in AAW-TNBCs (Fig. 5C both panels compare with Fig. 4C both panels, and Suppl. Figs 5A-D). Pathways functional enrichment revealed several major altered pathways (Fig. 5C, top panel) including the most affected signaling pathway which was the GnRH signaling pathway, similar to our RNA-seq data (Fig. 4C, top panel). Another one of the top 10 altered pathways included inflammatory chemokines and cytokines, which have a vast range of functionalities and may be directly responsible for the modulation of tumor growth by stimulating angiogenic factors from tumor cells (Balkwill, 2004). It has been shown that inflammation contributes to induction and metastasis of breast cancer as well as its homeostatic regulation (Klopocki et al., 2004). These molecules can prevent apoptosis, inducing proliferation of cancer cells (Esquivel-Velazquez et al., 2015), which is also one of the pathways affected in AAW-TNBC cells under steroid actions. Altered signaling pathways also included the Integrin pathway which helps with the process of cell adhesion, growth, division, proliferation, and apoptosis, and may be linked to induction of angiogenesis by activation of mitogen-activated protein kinases (Chow and Luster, 2014). Additionally, it has been shown that there is a relationship with the regulation of Integrin and IGF-IR in mitogenesis and prostate cancer development (Bergh et al., 2005). Integrin and GnRH are not only key regulators in both our RNA and proteomic pathways but have also been shown to mediate one another in ovarian cancer progression (Goel et al., 2005). This results from gonadotropins inducing expression of integrins and an overall increase in cell adhesion which is essential for invasion (Schiffenbauer et al., 2002). Modulation of the GnRH pathway is also seen in breast, prostate, and endometrium cancers (Schally, 1999). Wnt, another key pathway affected in AAW-TNBC cells under steroid actions, is responsible for signal transduction that occurs through cell surface receptors. Evidence associates that this pathway may be potentially connected to various types of cancers such as breast, prostate (Zimmerli et al., 2017), colorectal melanoma, and lung cancer (Anastas and Moon, 2013). Wnt has been classified as a proto-oncogene and its dysregulation can induce uncontrolled cell division producing tumors (Freese et al., 2010). Notably, activated secondary messengers like MAPK, through EGFR, can activate or couple to GPCRs (Filardo, 2002), all of which are dysregulated in our proteomic analysis of AAW-TNBC cells under steroid actions.

**Fig. 5.**
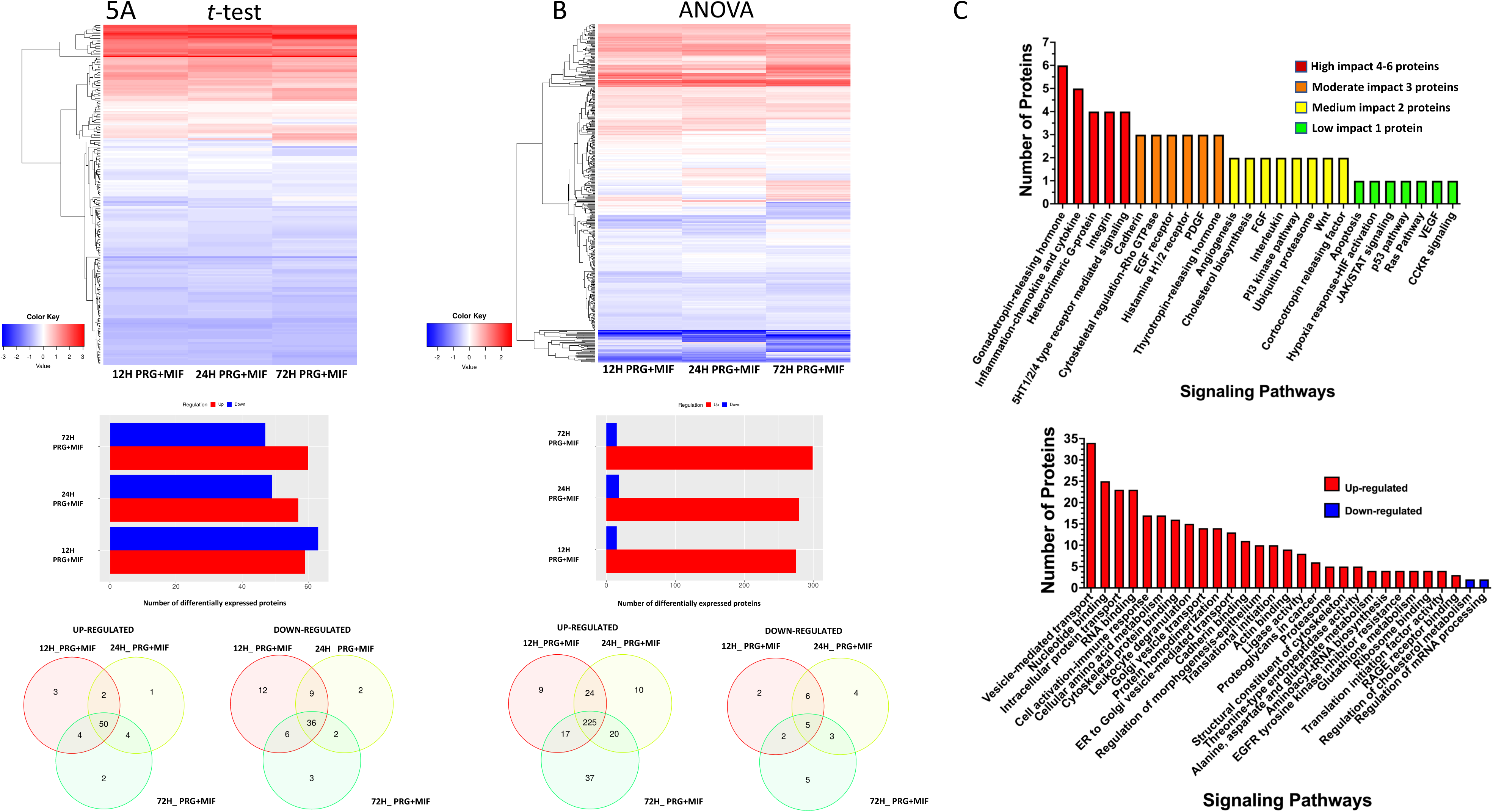
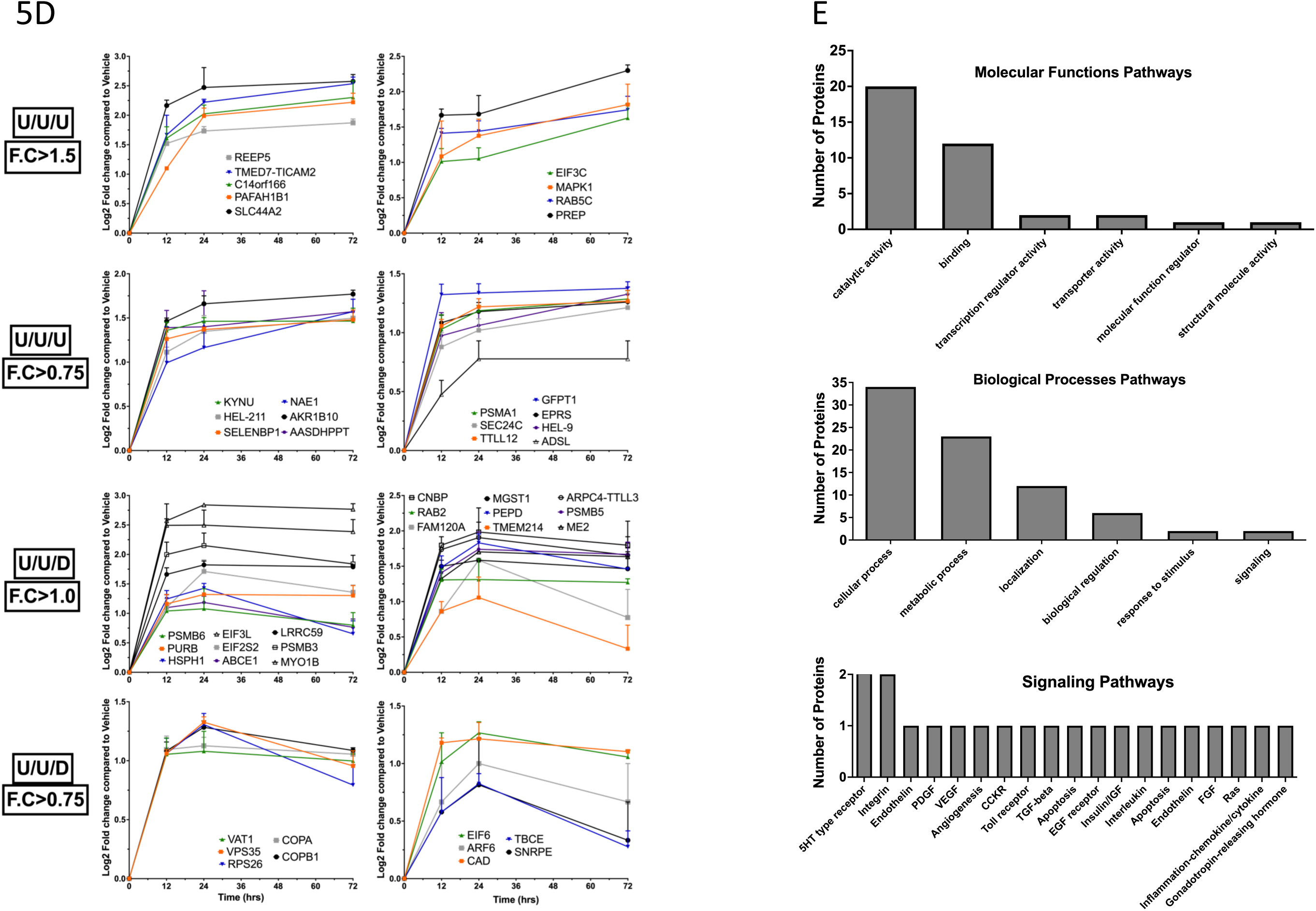
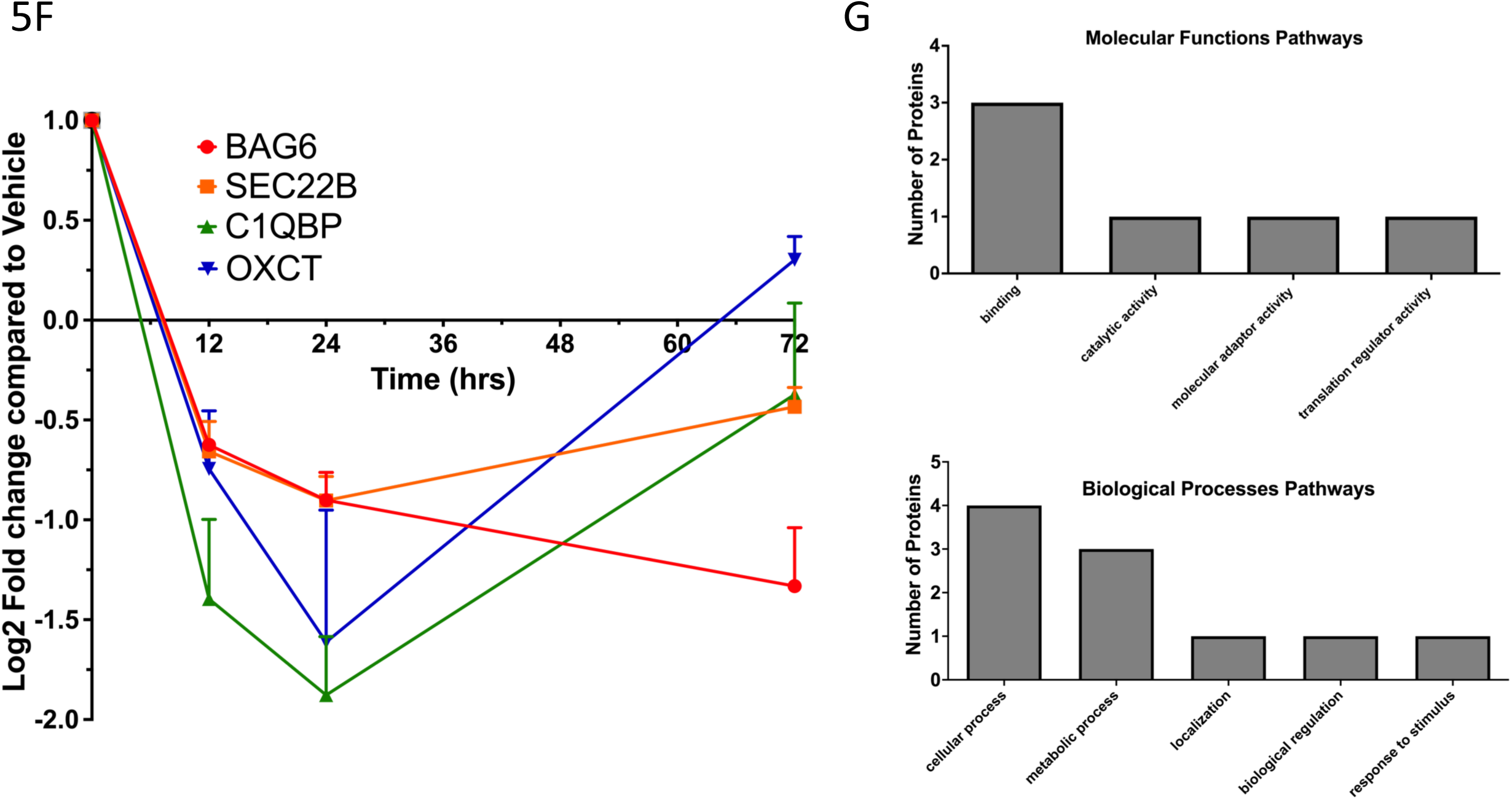
Differentially Expressed Proteins (DEPs) among AAW-TNBC cells utilizing high-throughput Proteomics: AAW-TNBC cells were treated with MIF+PRG (20µM each) for 12, 24 and 72 hrs. Extracted DEPs were analyzed using both *t-test* and ANOVA statistical analysis to increase identified DEPs, and were further filtered using corresponding MB231/Jurkat cells to obtain significant DEPs specific to AAW-TNBC cells. **A)** Cluster software and Euclidean distance matrixes were used for the hierarchical clustering analysis of the expressed proteins and sample program at the same time to generate the overall Heatmap of hierarchical clustering using *t-test* statistical analysis (top panel). Further analysis was performed to identify significant DEPs, compared to vehicle controls (middle panel). Significant DEPs were further stratified into independent Venn diagrams for both up and down regulated proteins to identify overlaps between timepoints (Bottom panels). **B)** Cluster software and Euclidean distance matrixes were used for the hierarchical clustering analysis of the expressed proteins and sample program at the same time to generate the overall Heatmap of hierarchical clustering using ANOVA statistical analysis (top panel); Further analysis was performed to identify significant DEPs, compared to vehicle controls (middle panel). Significant DEPs were further stratified into independent Venn diagrams for both up and down regulated proteins to identify overlaps between timepoints (Bottom panels)**. C)** Pathway functional enrichment results for MIF+PRG treated cells compared to their respective vehicle controls were compiled using molecular functions, biological processes and signaling pathway data retrieved from the PANTHER classification system (top panel). For upper panel C, coloring indicates the potential impact on each signaling pathway; high (4-6 proteins): red, moderate (3 proteins): orange, medium (2 proteins): yellow, low (1 protein): green. Additionally, signaling pathways were further stratified to identify up/down regulation within only signaling pathways using iDEP program (bottom panel). **D)** To delineate cellular partners sharing the same/reverse inducible expression patterns as CCM1 using high-throughput omics. Proteomics analysis was temporally performed to identify consistently up-regulated DEPs (statistically significant); up-regulated (u) trends included u/u/u with peak Log2FC categories greater than 1.5 *(top panels),* u/u/u with peak Log2FC categories greater than 0.75 *(second panels)*, u/u/d (d, down) with peak Log2FC categories greater than 1.0 (third panels), or u/u/d with peak Log2FC categories greater than 0.75 (fourth panels). **E)** Pathway functional enrichment results for corresponding proteins shown in panel D were compiled using molecular functions, biological processes and signaling pathway data retrieved from the PANTHER classification system. **F)** Proteomics analysis was performed to identify consistently down regulated DEPs (statistically significant). **G)** Pathway functional enrichment results for corresponding proteins shown in panel F were compiled using molecular functions and biological processes data retrieved from the PANTHER classification system. For all heatmaps, x axis represents each comparing sample and Y axis represents DEPs. For all functional enrichment graphs, X axis represents the terms of pathway while y axis represents the number of identified DEPs involved in each pathway. For all panels, each timepoint was run with biological triplicates and technical triplicates for treated and vehicle samples. For panels (A-B, and lower C), Coloring indicates the log2 transformed fold change (high: red, low: blue).

To delineate cellular partners sharing the same/reverse inducible expression patterns as CCM1 (Fig. 2D, left panel), as we did with our transcriptomics data, we sought to elucidate these partners at the translational level as well. We identified 50 consistently up-regulated proteins (Suppl. Table 5B) with similar expression patterns as CCM1 (Fig. 5, panel D). Some notable proteins included CNBP, which functions as a sterol-mediated transcriptional factor and overexpression has been linked with tumor cell biology, in which CNBP acts as a key transcriptional regulator of tumor-promoting target genes to control tumor cell biology (Lee et al., 2019). Additionally, 30 of the 50 consistently up-regulated proteins have been shown to be differentially expressed in numerous cancers including breast, cervical, liver, lung, and brain cancer, contributing to tumor growth, metastasis and high invasive activities (Suppl. Table 5C). We observed several of the previously altered signaling pathways (detected using DEGs) for these up-regulated DEPs (Fig. 5, panel D), including Integrin, Inflammation, toll receptor, and interleukin signaling pathways (Figs. 4E/5E). Finally, we identified only 4 DEPs with inverse expression patterns to CCM1 (Fig. 5F, Suppl. Table 5B). Interestingly, SNARE protein SEC22B has been shown to modulate many pathological and physiological processes including autophagy, vesicle trafficking, neurodegeneration, cancers as well as infectious diseases (Sun et al., 2020). SEC22B is closely related to tumorigenesis and has been confirmed in numerous cancers including lung, breast, prostate, and uterine cancers (Gao et al., 2013), and furthermore, SEC22B has been shown to fuse with NOTCH2 and was confirmed in aggressive breast cancer (Veeraraghavan et al., 2016). Interestingly, in hepato-carcinogenesis SEC22B interacts with VPS33B to regulate distribution of E-cadherin in hepatocytes related to the progression of cancers by increasing proliferation, invasion and metastasis (Wang et al., 2018). C1QBP was also identified in our analysis and has been established as having a pro-tumor role in breast cancer that could serve as a molecular target for cancer therapeutics (McGee et al., 2011). We also observed similar regulatory effects (seen at the transcriptional level) in the analysis of signaling pathways for these down-regulated DEPs (Fig. 5, panel F) including biological regulation, response to stimuli and localization pathways (Fig. 5, panel G).

Together, these results solidify the existence of a cellular relationship between the CSC and mPRs signaling at the translational level by establishing the CmP signaling network in AAW-TNBC cells. Our extensive omics analysis has also provided several new candidate biomarkers, at both the transcriptional and translational levels, that can be further investigated to determine their potential clinical applications for AAW-TNBCs.

### Key signaling cascades *within the CmP network in AAW-TNBC cells* identified using systems biology approaches

To look at the inter-relationship between RNA and protein expression from AAW-TNBC cells under steroid actions, we overlaid the regulation patterns and extrapolated the overlaps between the two datasets (Suppl. Table 6A). When looking at the enriched annotations we see pathways involved in the regulation of cancer and tumor factors still consistently present from the individual analyses performed. Key pathways including Integrin, cytokine-mediated responses, GnRH, WNT, and angiogenesis are still among the top 10 affected pathways in our multi-omics analyses (Fig. 6 panel A), further validating our independent observations at both the transcriptional and translational levels (Figs. 4-5). When reviewing expression patterns between RNA and protein analysis it should be noted that genes with the same regulation trends (averaged log2 Fold Changes) seen at both the transcriptional and translational levels (Bolded blue and red ID’s) are key AAW-TNBC specific candidates for potential biomarkers (Fig. 6 panel B), while genes displaying opposite expression at the transcriptional and translational levels (black colored ID’s) commonly reflect genes that are capable of undergoing potential feedback auto-regulation in AAW-TNBC cells under steroid actions (Fig. 6 panel B and Suppl. Table 6B). Up-regulated genes illustrate a significant impact on essential cancer pathways including Integrin, cytoskeletal regulation by Rho GTPase, angiogenesis, VEGF, GNRH, G-protein, EGF, PDGF and Cadherin signaling pathways (Fig. 6 panel B). Among the up-regulated genes, ARF6 has been shown to be involved in tumor invasion and metastasis when overexpressed (Li et al., 2017). Additionally, several of the up-regulated DEGs/DEPs have been shown to be differentially expressed in numerous cancers including breast, cervical, liver, lung, and brain cancer, contributing to tumor growth, metastasis and high invasive activities (Suppl. Table 5C).

**Figure 6:**
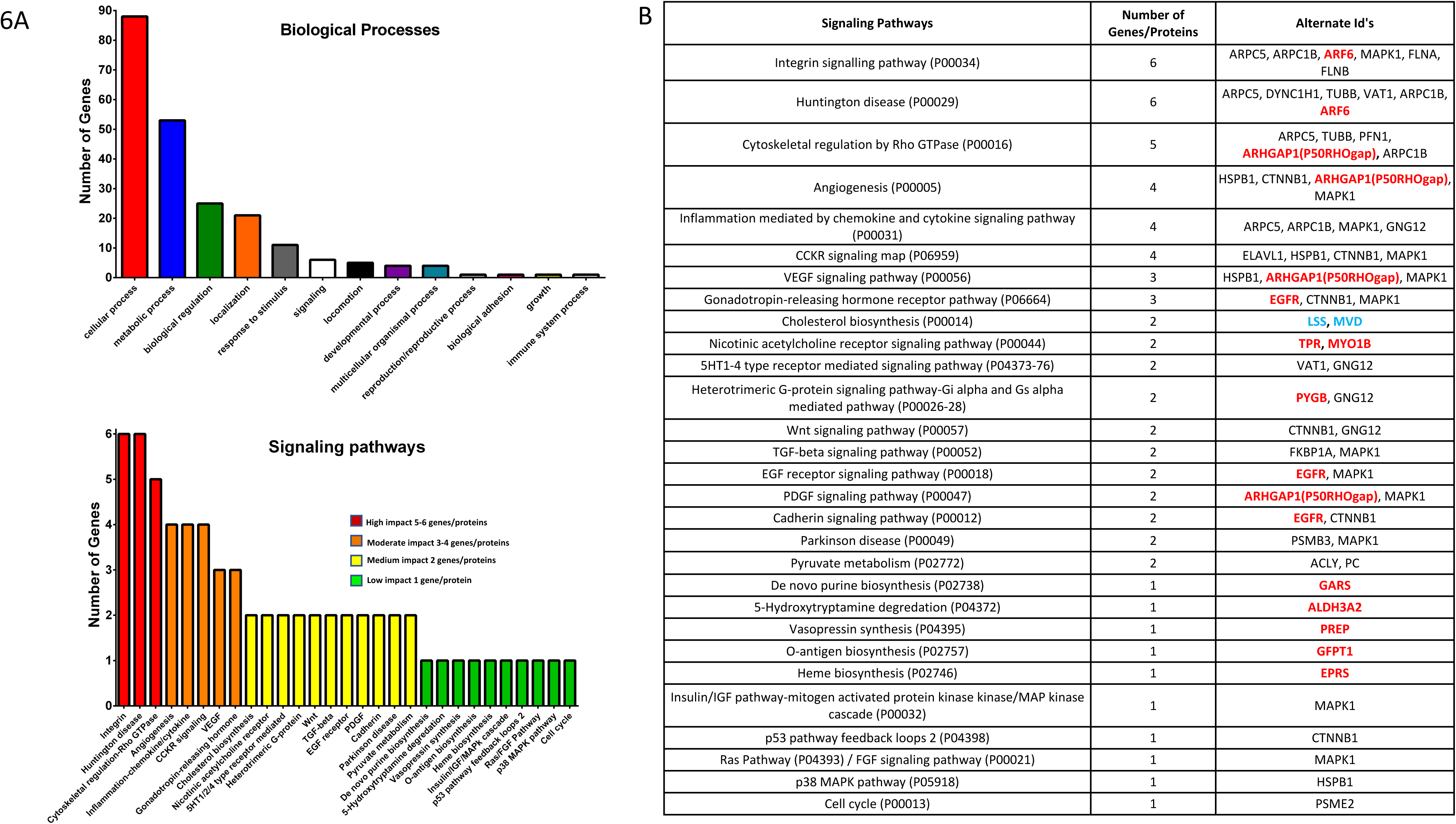
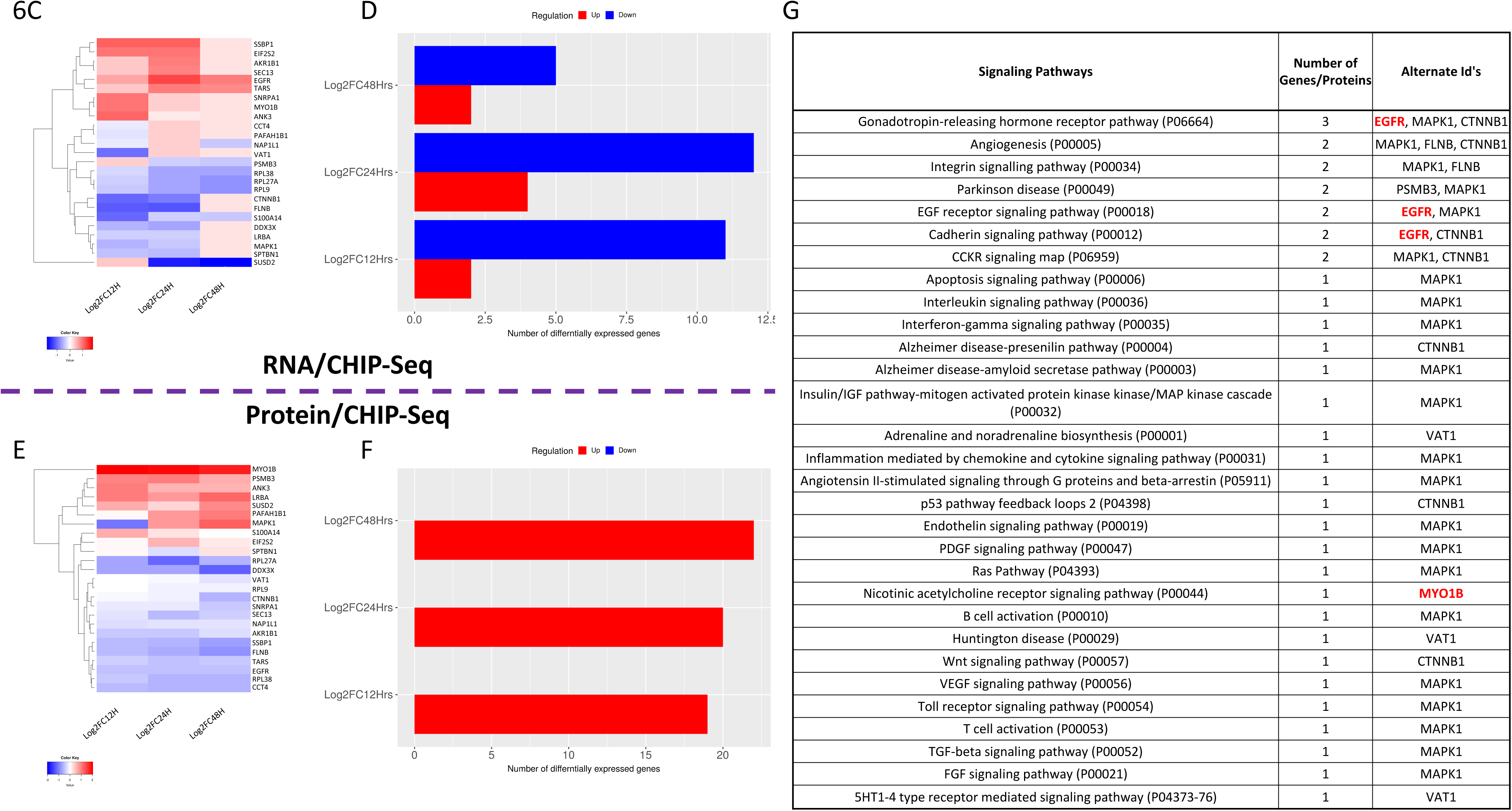
Systems biology analysis utilizing high-throughput omics data to identify novel biomarkers unique to tumorigenesis in AAW-TNBC cells under steroid actions: Extracted significant DEGs/DEPs were further filtered using corresponding MB231/Jurkat cells to obtain DEGs/DEPs specific to AAW-TNBC cells. **A.** Pathway functional enrichment results for corresponding DEGs/DEPs were compiled using Biological processes and signaling pathways data retrieved from the PANTHER classification system. For bottom panel, coloring indicates the potential impact on each signaling pathway; high (5-6 genes/proteins): red, moderate (3-4 genes/proteins): orange, medium (2 genes/proteins): yellow, low (1 gene/protein): green. **B.** A corresponding table is provided for each signaling pathway that details the genes involved, and the statistical analysis used with the proteomics data (A, ANOVA; T, *t-test*) for the identified proteins. Genes/Proteins with similar regulation at both the transcriptional and translational levels are color coded and bolded with red DEGs/DEPs (up-regulated) or blue DEGs/DEPs (down-regulated) while Genes/Proteins in black display opposite expression at the transcriptional and translational levels (black colored ID’s). **C-F** Cluster software and Euclidean distance matrixes were used for the hierarchical clustering analysis of DEGs/DEPs along with overlaps in CHIP-seq databases to generate the overall Heatmap of hierarchical clustering (panels C and E); x axis represents each comparing sample and Y axis represents DEGs/DEPs. Further Bioinformatics analysis was performed to stratify significant DEGs/DEPs, compared to vehicle controls that overlapped with CHIP-seq databases in hormone treated breast cancer cells (panels D and F). **G.** A corresponding table is provided with affected signaling pathways that details the overlapped genes/protein data from our omics studies and supporting CHIP-seq data. DEGs/DEPs with similar regulation at both the transcriptional and translational levels are color coded and bolded with red DEGs/DEPs (up-regulated), while Genes in black display opposite expression at the transcriptional and translational levels (black colored ID’s). Pathway functional enrichment results were performed using signaling pathway data retrieved from the PANTHER classification system. For all functional pathway’s analysis, X axis represents the terms of pathway while the Y axis represents the number of identified DEGs/DEPs involved in each pathway. For panels B-G, Coloring indicates the log2 transformed fold change (high: red, low: blue). For all panels, each timepoint was run with biological duplicates (RNA) or triplicates (Protein) and technical triplicates (both) for treated and vehicle samples.

We then performed a meta-analysis using a working database (Suppl. Table 6C) with CHIP-seq data from various breast cancer cells (containing MDA-MB-468 cells) treated with either PRG/MIF that encompassed binding domains to help elucidate potential mechanisms of action from known protein characteristics (Xi et al., 2018). This task was accomplished by using the NCBI Batch Web CD-Search tool (Marchler-Bauer and Bryant, 2004; Mellacheruvu et al., 2013). The list of overlapped genes (Suppl. Table 6C) was graphed through Euclidian heat maps for both RNA and protein as well as their regulation patterns (Figure 6, panel C-F). In general, we observed that the majority of the genes identified in our systems-wide analysis with combined RNA-seq/Proteomics/CHIP-seq data appear to be capable of being auto-regulated through feedback mechanisms in AAW-TNBC cells under steroid actions. However, we still identified two steroid-inducible AAW-TNBC specific candidate biomarkers that were coordinately up-regulated at both the transcriptional and translational levels namely EGFR and MYO1B (Figure 6, panel G). Interestingly, EGFR has been shown to enhance chemosensitivity of TNBCs by rewiring the apoptotic signaling network and high EGFR expression was an independent predictor of efficacy of chemotherapy and poor prognosis in inflammatory breast cancer and TNBCs (Babyshkina et al., 2020; Masuda et al., 2012). Additionally, identified MYO1B has been shown to have an important role in progression of several cancers, including prostate and head and neck squamous cell carcinoma as well as having a potential role in cervical carcinogenesis and tumor progression (Zhang et al., 2018).

### Differential expression of our candidate biomarkers in breast cancer tissues and their associated prognostic effects

We set out to filter our identified candidate biomarkers for AAW-TNBCs from our omics approaches to obtain a more clinically relevant set of prognostic biomarkers. To accomplish this, 11 publicly available breast cancer datasets from University of California-Santa Cruz (UCSC) Xena Browser (Chin et al., 2006; Desmedt et al., 2007; Goldman et al., 2020; Heiser et al., 2012; Hess et al., 2006; Miller et al., 2005; Naderi et al., 2007; Neve et al., 2006; van’t Veer et al., 2002; van de Vijver et al., 2002; Wang et al., 2005; Yau et al., 2010) as well as two TCGA-BRCA databases available in Xena Browser (TCGA-BRCA and GDC-TCGA-BRCA) were compiled to validate differential expression among Breast cancer subtypes for our Identified candidate biomarkers (Suppl. Table 6D). Additionally, we evaluated prognostic effects for our candidate biomarkers (Gyorffy et al., 2010) integrating clinical and gene expression data simultaneously to generate Kaplan-Meier (KM) survival curves. Breast cancer patient samples were filtered to only analyze patient samples classified as ER(-)/PR(-)/HER2(-)/TNBC subtype, yielding 176/392 patient samples to be utilized. For the first group of biomarkers, a worst prognostic effect was observed with higher expression of ACOT9, AIMP1, ALDH3A2, ARF6, P50rhoGAP, CNBP, EML2, G6PD, GARS, LANCL2, CNN3 and LSS (Figure 7A1-7A12). We also observed up-regulation of 10/12 biomarkers (CNN3 and LSS were down-regulated) with a disrupted CmP signaling network, under steroid actions, in TNBC cells (Suppl. Table 6D), demonstrating that combined steroid actions could lead to worst prognostic outcomes in AAW-TNBCs. Alternatively, our second group of identified biomarkers demonstrated a better prognostic effect with higher expression of AKR1B1, GAI7, DDX1, AKR1C2, HEBP2, EGFR, COPE, PYGB, and SSBP1 (Figure 7B1-7B9). Interestingly, we observed up-regulation of all 9 genes/proteins with a disrupted CmP signaling network, under steroid actions, in TNBC cells (Suppl. Table 6D). Together these results demonstrate the heterogeneity response of TNBCs under steroid treatments. Utilizing our KM survival curve data, we further assessed our biomarkers by only proceeding with biomarkers that had significant KM survival curves in TNBC patients to assess their basal expression patterns in clinical samples.

**Figure 7:**
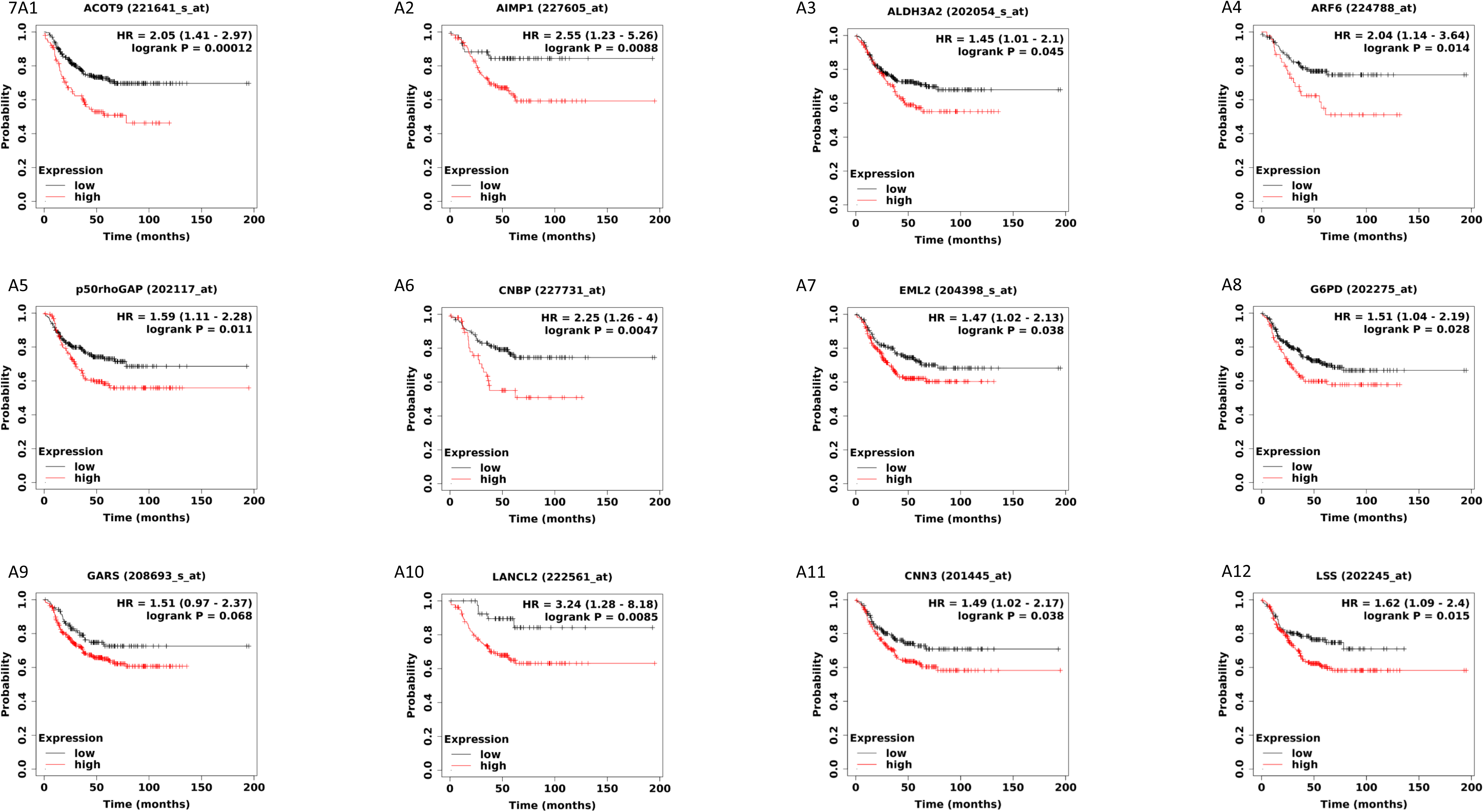
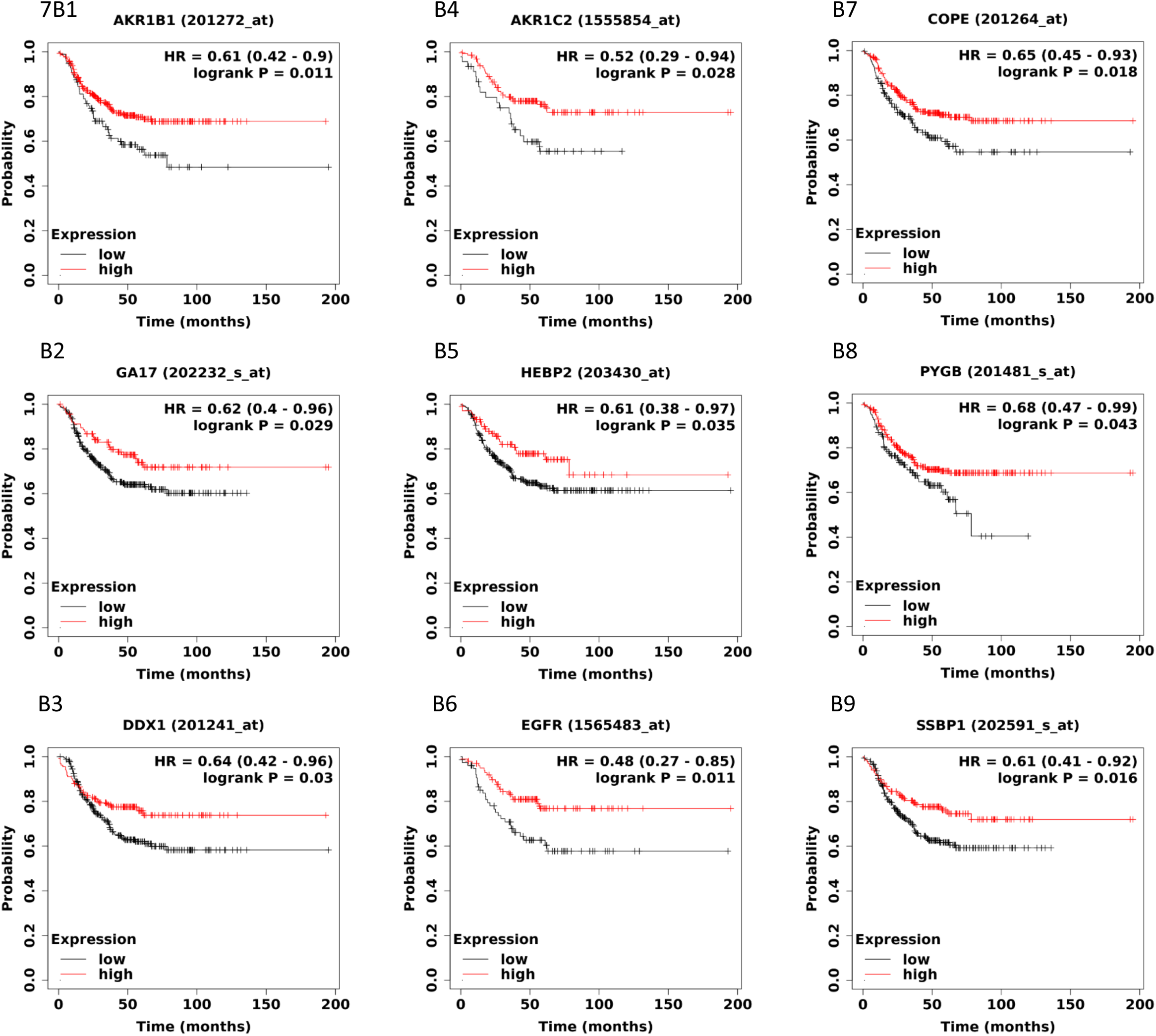
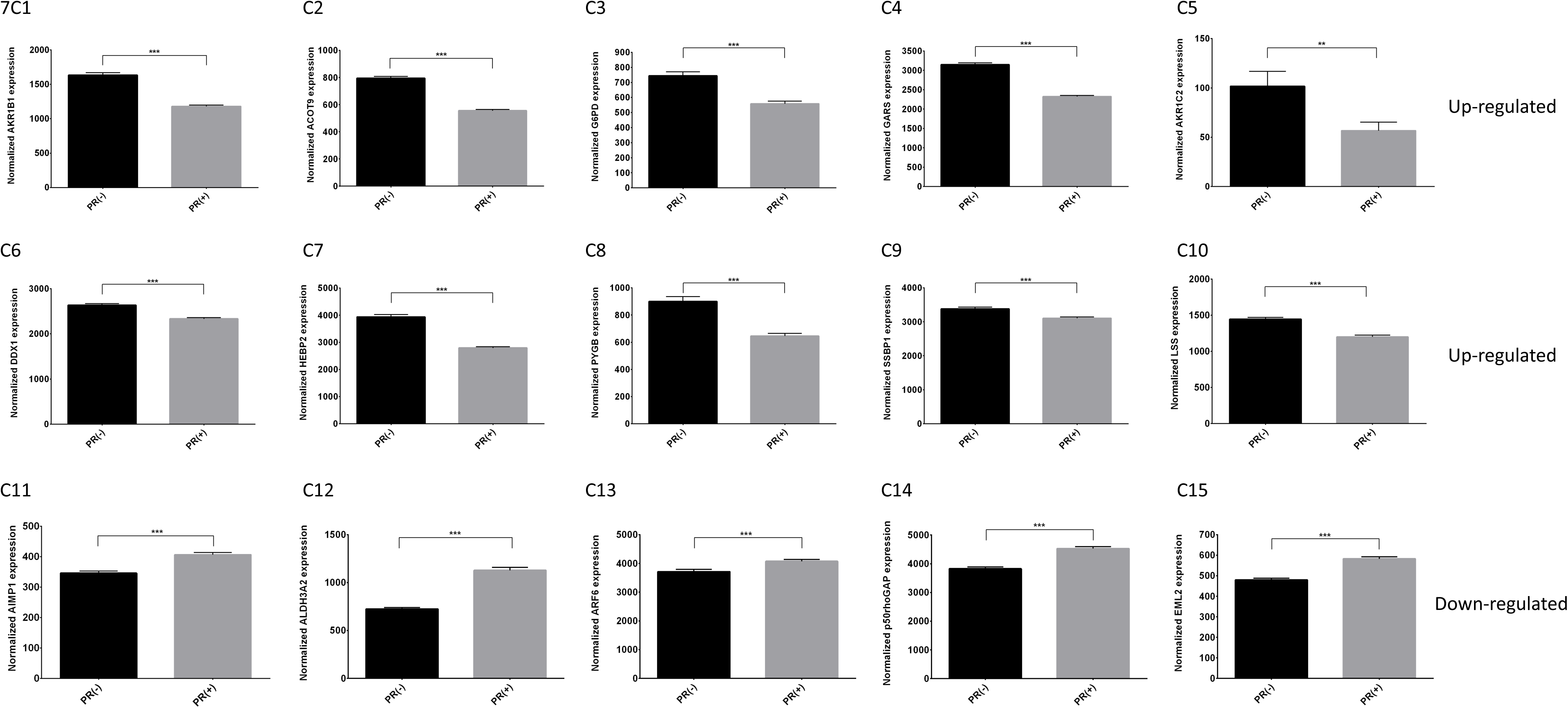
Expression and prognostic effects for identified candidate biomarkers utilizing microarray data of Breast cancer patient samples. Publicly available microarray data (22,277 probes) from 1,809 breast cancer patients was analyzed using kmplot software to integrate gene expression and clinical data simultaneously to generate the displayed Kaplan-Meier survival curves. Breast cancer patients were filtered to only analyze patient samples classified as ER(-)/PR(-)/HER2(-)/TNBC subtype, which reduced our 1,809 patients to 176/392 patient samples. Results demonstrated **A)** significantly worst prognostic effect with higher expression of **A1)** ACOT9, **A2)** AIMP1 **(A3)** ALDH3A2 **(A4)** ARF6 **(A5)** P50rhoGAP**, (A6)** CNBP, **A7)** EML2, **(A8)** G6PD, **(A9)** GARS, **(A10)** LANCL2, **(A11)** CNN3 and **(A12)** LSS. We observed higher expression of genes (A1-A10) but lower expression of genes (A11-A12) in our omics studies under a disrupted CmP network (combined steroid actions) in AAW-TNBCs. Alternatively, Results also demonstrated **B)** significantly better prognostic effects with higher expression of **B1)** AKR1B1, **B2)** GAI7 **(B3)** DDX1 **(B4)** AKR1C2 **(B5)** HEBP2**, (B6)** EGFR, **B7)** COPE **(B8)** PYGB **(B9)** and SSBP1. We observed higher expression of these genes in our omics studies under a disrupted CmP network (combined steroid actions) in AAW-TNBCs. Logrank P values are calculated and displayed as well as hazard ratio (and 95% confidence intervals). Red line demonstrates high gene expression, while black line demonstrates low gene expression. **C)** Microarray data containing breast cancer tumors, analyzed using kmplot software, were divided into two groups based on classic Progesterone Receptor (PR) status determined by Immunohistochemistry (IHC); after filtering, there were 925 PR(-) and 926 PR(+) breast cancer samples. Up-regulation of candidate biomarkers, in PR(-) breast cancer tissue samples, compared to PR(+) tissues, included **C1)** AKR1B1, **C2)** ACOT9 **(C3)** G6PD **(C4)** GARS **(C5)** AKR1C2**, (C6)** DDX1, **C7)** HEBP2, **(C8)** PYGB, **(C9)** SSBP1, and **(C10)** LSS, while down-regulation of candidate biomarkers in PR(-) breast cancer tissue samples, compared to PR(+) tissues, included **C11)** AIMP1, **C12)** ALDH3A2 **(C13)** ARF6 **(C14)** P50rhoGAP and **(C15)** EML2. All biomarkers in panels C displayed significant survival curves in panels A and B, for breast cancer cells. Statistical significance was performed with students *t*-test where **, *** above bar indicates P ≤ 0.01 or 0.001, respectively.

Utilizing publicly available breast cancer tumor tissue gene expression data (microarray) (Gyorffy et al., 2010) we explored expression patterns of our identified candidate biomarkers in nPR(+/-) breast cancer cells (ER and HER2 receptor status not available). Expression data was obtained for our candidate biomarkers from tissue data that was divided based on Progesterone Receptor status (nPR+/-) resulting in 925 nPR(-) and 926 nPR(+) breast cancer samples. 10 of our candidate biomarkers, AKR1B1, ACOT9, G6PD, GARS, AKR1C2, DDX1, HEBP2, PYGB, SSBP1 and LSS were found to be significantly up-regulated in nPR(-) tumor tissues, compared to nPR(+) tumor tissues (Fig. 7C1-C10). We observed the same up-regulated trends for 9/10 biomarkers, with the exception of LSS, with a disrupted CmP signaling network under steroid treatment conditions in AAW-TNBC cells (Suppl. Table 6D), suggesting little effect of hormone actions on their expression levels. LSS however, was down-regulated with a disrupted CmP signaling network suggesting a disrupted CmP signaling network, under hormone actions, can alter expression, at both the transcriptional and translational levels, in TNBC cells (Suppl. Table 6D). Additionally, 5 of our candidate biomarkers, AIMP1, ALDH3A2, ARF6, P50rhoGAP, and EML2 were found to be significantly down-regulated in nPR(-) tumor tissues, compared to nPR(+) tumor tissues (Fig. 7C11-C15). Interestingly, we observed opposing up-regulation of these 5 biomarkers with a disrupted CmP signaling network, suggesting a disrupted CmP signaling network, under hormone actions, alters expression, at both the transcriptional and translational levels, in AAW-TNBCs (Suppl. Table 6D) for these naturally under expressed genes in nPR(-) tumor tissues.

Furthermore, 6 of our biomarkers, EGFR, CNBP, LANCL2, COPE, GAI7, and CNN3 were found to have similar expression patterns in nPR(+/-) tumor tissues (Suppl. Fig. 7). However, disruption of the CmP signaling network, with combined steroid actions, resulted in significant up-regulation of 5/6 biomarkers, with the exception of CNN3 which was down-regulated (Suppl. Table 6D). These results suggest that combined steroid actions induces altered expression, at both the transcriptional and translational levels for these 6 biomarkers in AAW-TNBC cells.

We next sought to not only evaluate expression differences between nPR(+/-) breast cancer tissues, but wanted to further our analysis to investigate differential expression patterns between AAW-TNBCs and CAW-TNBCs. To accomplish this task, we utilized publicly available RNA-seq data comparing expression levels in 23 AAW-TNBC and 19 CAW-TNBC samples (Saleh et al., 2021) that were used to obtain expression data for the candidate biomarkers associated with significant KM curves identified in this study. AKR1B1, G6PD, GARS, LANCL2, DDX1, PYGB, SSBP1, COPE, and CNN3, all displayed similar trends with a disrupted CmP signaling network, as was naturally observed in the 23 AAW-TNBC patient samples, compared to CAW-TNBC samples (Suppl. Table 6D, red and blue colored biomarkers, column 1). This allowed us to define these genes/proteins as intrinsic biomarkers for AAW-TNBCs. Alternatively, EGFR, ACOT9, AIMP1, ALDH3A2, ARF6, P50rhoGAP, EML2, CNBP, AKR1C2, HEBP2, LSS and EIF3M displayed either opposite or inducible trends with a disrupted CmP signaling network in AAW-TNBC cells, as was naturally observed in the 23 AAW-TNBC patient samples, compared to CAW-TNBC samples, allowing us to define these genes/proteins as inducible biomarkers (CmP disruption) for AAW-TNBCs (Suppl. Table 6D, black colored biomarkers).

Utilizing all of the expression and prognostic data, we filtered out our initial 47 biomarker hits (Suppl. Table 6D) to a condensed, clinically relevant list of 21 intrinsic and inducible biomarkers for AAW-TNBCs (Table 1).

**Table 1:**
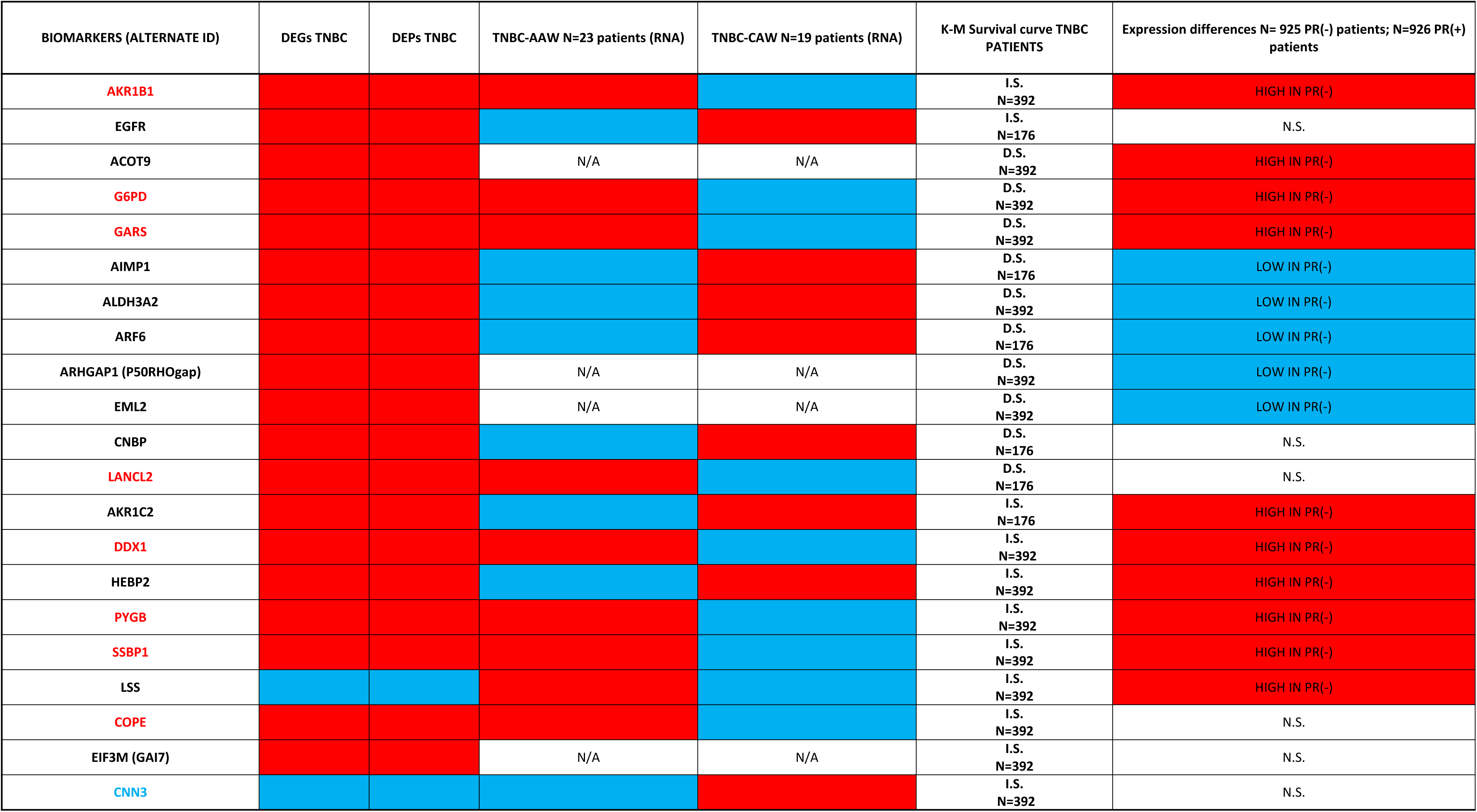
List of 21 identified prognostic candidate biomarkers associated with clinically significant data in AAW-TNBCs. Candidate biomarkers shared at both RNA/Protein levels under hormone treatments (columns 1, 2, 3), were further validated in AAW-TNBC (n=23) and CAW-TNBC (n=19) (columns 4, 5) tumor tissues. These biomarkers were further analyzed by a set of publicly available microarray data (22,277 probes) from 1,809 breast cancer samples using kmplot software to integrate gene expression and clinical data simultaneously to generate the displayed Kaplan-Meier survival curves (column 6). Candidate biomarkers were further assessed with two groups of breast cancer tissue samples, based on classic progesterone Receptor (PR) status determined by Immunohistochemistry (IHC), to evaluate clinical expression levels focusing on 925 PR(-) breast cancer samples (column 7). Red colored background indicates increased/higher expression levels, while blue colored background indicates decreased/lower expression levels. Colored biomarkers (red or blue in Column 1) indicate intrinsic biomarkers which are selected as candidate biomarkers for AAW-TNBCs, while black colored biomarkers are inducible biomarkers, like CCM1. Abbreviations: I.S., Increased Survival; D.S., Decreased Survival; N.S., No statistical significance; HIGH IN PR(-), Higher expression levels in nPR(-) breast cancer tissues; LOW IN PR(-), Lower expression levels in nPR(-) breast cancer tissues; N/A, Not Available.

## Discussion

There has been sufficient evidence for the existence of PRG-mPRs signaling cascade in either nPR(+/-) cells (Dosiou et al., 2008; Karteris et al., 2006; Pang and Thomas, 2011; Sleiter et al., 2009; Zuo et al., 2010), suggesting that PRG signaling in nPR(-) cell lines can be mediated solely through mPRs (PAQRs) (Zuo et al., 2010). In this current study, we further defined the novel CSC-mPRs-PRG (CmP) signaling network in nPR(-) breast cancer cells (Fig. 8) which overlaps with our previously defined CmPn (CSC-mPRs-PRG-nPRs) network in nPR(+) breast cancer cells (Abou-Fadel et al., 2020b) using *in-vitro* models. In the CmP signaling network, the CSC is able to stabilize mPRs under steroid actions in a forward fashion (CSC→mPRs), indicating a more essential role of the CSC on the stability of mPRs (PAQRs) in nPR(-) breast cancer cells under steroid actions. Overall, the CSC complex can stabilize the expression of mPR proteins in TNBC cells (Fig. 1B), in concordance with our observations in nPR(+) breast cancer cells, indicating the consistent function of the CSC on the stability of mPRs under steroid actions (Abou-Fadel et al., 2020b). In contrast to previously observed nPR(+) breast cancer and nPR(-) endothelial cell data (Abou-Fadel et al., 2020a; Abou-Fadel et al., 2020b), combined steroid actions have a positive effect on protein expression on the CSC in both TNBC sub-types with the inducible pattern of expression more apparent in Type 2B AAW-TNBC cells (Figs. 2E, 3A-C). Interestingly, RNA expression of CCM genes is also enhanced with combined steroid treatment, while variable effects are observed in Type 2A CAW-TNBC cells (Fig. 1C, 2D). In general, expression levels of mPRs are enhanced at both the transcriptional and translational levels in Type 2A cells under combined steroid treatment, while mPRs expression is variable at both the transcriptional and translational level in Type-2B TNBC cells, making these genes great candidate biomarkers for TNBC sub-type classification (Fig. 8). Another major finding is that mPRs are localized within the nucleus in both TNBC cells (Figs. 3A-D), and exhibit capabilities of shuttling between the nucleus and cytosol (Fig. 3D), identical to the cellular compartmentations of other steroid receptors, making mPRs a new type of PRG receptors functionally similar to classic nPRs, which can perform both genomic and non-genomic PRG actions. Extensive multi-omics analysis was performed confirming alterations in key tumorigenesis pathways including Integrin, cytokine-mediated responses, GnRH, WNT, and angiogenesis signaling pathways. Additionally, our multi-omics approach identified 21 new AAW-TNBC candidate biomarkers (Table 1) that will be investigated further to evaluate their potential clinical applications for AAW-TNBCs.

**Fig. 8:**
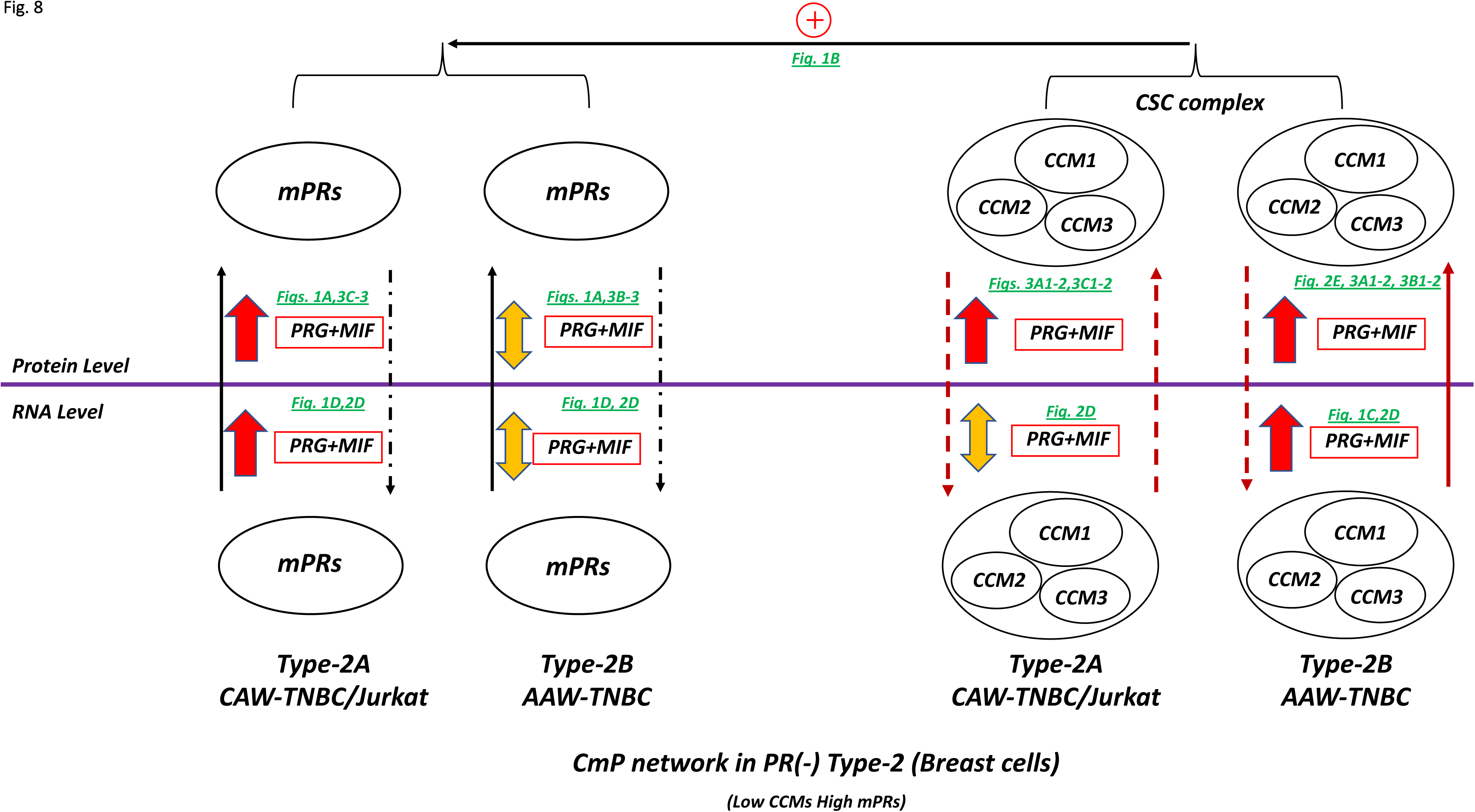
Schematic model of the CSC-mPRs-PRG (CmP) signaling network under combined steroid treatment for TNBC cells. Utilizing our molecular works, we summarized the CmP signaling network in TNBC cells under combined steroid actions. In general, combined steroid actions (PRG+MIF) have a positive effect on protein expression of the CSC in both TNBC sub-types but the inducible pattern of expression is more apparent in type 2B (AAW-TNBC) cells. Interestingly, RNA expression of CCM genes is also enhanced with PRG+MIF treatment, while variable effects are observed in Type 2A (CAW-TNBC) cells. In general, expression levels of mPRs are enhanced at both the transcriptional and translational levels in Type 2A cells under combined steroid treatment, while mPRs expression is variable at both the transcriptional and translational levels in Type 2B cells. Overall, the CSC complex can stabilize expression of mPR proteins in TNBC cells, in concordance with our observations in nPR(+) breast cancer cells, indicating the consistent function of the CSC on the stability of mPRs. The purple line separates transcriptional and translational levels. The + symbols represent enhancement, for the expression of targeted genes/proteins. Red colored symbols/lines represent positive effects of combined steroid treatment, blue colored symbols/lines represent negative effects of treatment, while yellow colored symbols represent variable effects, temporally sub-cellular compartmentation and/or fluctuating expression along the time-course under steroid actions. Dark green colored letters indicate the direct supporting data generated from this report. Arrow indicates effect direction, solid line is the direct impact, dotted line for indirect effects.

This project provides new insights into the CmP signaling network in TNBC tumorigenesis, which may revolutionize the current concepts and molecular mechanisms of tumorigenesis in AAW-TNBC, and reproductive cancers in general, leading to new therapeutic strategies.

### Novel discoveries in the CmP signaling network with potential revolutionary impacts

#### The CmP signaling network and mPRs subcellular compartmentations

mPR expression patterns in many tissues and cell lines, and the possible involvement of mPRs in various biological processes have been extensively studied, however, the precise functions of mPRs remain controversial (Tokumoto et al., 2016), and even more controversial data on their subcellular compartmentation (Salhi et al., 2010). Up to date, mPRs have been agreed upon as a new type of PRG receptor, with only confirmed non-genomic PRG-actions, and while their stimulation varies drastically in a tissue- and cell-dependent fashion, debates on their subcellular compartmentation continue (Salhi et al., 2010). As mentioned earlier, mPRs were initially identified as a new family of membrane-bound PRG receptors (Smith et al., 2008; Zhu et al., 2003a; Zhu et al., 2003b), and PRG-mPRs signaling cascades, as well as their mediated cellular functions, have been investigated in various organisms and cell types (Josefsberg Ben-Yehoshua et al., 2007; Kasubuchi et al., 2017; Pang et al., 2015; Tischkau and Ramirez, 1993; Xie et al., 2015). However, recently, evidence of cytoplasmic localization or cytoplasmic compartmentation of mPRs in various cell types, including MB231 cells, have emerged (Krietsch et al., 2006; Zhu et al., 2003a). Initial pieces of evidence of cytoplasmic vesicles of mPRs (Krietsch et al., 2006; Zhu et al., 2003a) were speculated as the results of either technical issues/cytotoxic events for recombinant mPR proteins from transiently transfected constructs (Zhu et al., 2008) or found in the membrane of the endoplasmic reticulum (ER) (Krietsch et al., 2006). Recently, contradicting evidence has emerged of localized recombinant mPRα protein into the ER of CHO cells, while others reported mPRα in the nuclear membrane of small and large luteal cells (Ashley et al., 2006). Additionally, reports showed cytoplasmic (vesicles) and nuclear membrane localization of different recombinant mPRα proteins in COS1 cells (Fernandes et al., 2005), while cytoplasmic (vesicles, ER) localization of various forms of recombinant mPRs in MB231 (Krietsch et al., 2006), HEK293, LLC-PK and MDCK cells (Lemale et al., 2008) were also reported. The question of whether the cytoplasmic localization of mPRs is due to artifacts of transient transfection (Zhu et al., 2008), clathrin-mediated endocytosis to cytoplasmic endosomes (Foster et al., 2010; Nader et al., 2021), lack and/or deficiency of PGRMC1 (Thomas et al., 2014), or misclassification of mPRs as membrane proteins (Fernandes et al., 2008; Salhi et al., 2010) are still under debate. In this paper, we have discovered that mPRs are predominantly nuclear proteins and furthermore are capable of shuttling in and out of the nucleus as necessary under steroid actions.

Our novel finding of the cytoplasmic-nuclear trafficking of mPRs will certainly revolutionize the current view of the functions and subcellular compartmentation of mPRs. Since only endogenous mPRβ was examined for this revolutionary discovery, questions about the quality and specificity of primary antibodies will certainly be raised despite their excellent and consistent outcomes in multiple experiments. After extensive examination, we found previous data from recombinant mPRs in transient transfection experiments, in various cells, that support our observed results. The nuclear compartmentation with strongly condensed localization within the nucleolus of recombinant NFlag-mPRα protein in HEK293 cells (Lemale et al., 2008), and in MDCK cells (Salhi et al., 2010) match our data of endogenous mPRβ in MB231 cells (Fig. 3A3); furthermore, the nuclear localization of endogenous mPRα in M11 cells (Foster et al., 2010) and COS-1 cells (Krietsch et al., 2006) provide more supportive evidence.

The most common and significant characteristic shared by all steroid hormone receptors is the ability for cytoplasmic-nuclear trafficking (Guiochonmantel and Milgrom, 1993). The molecular mechanisms of nucleocytoplasmic shuttling have been well defined for AR (Kesler et al., 2004; Tan et al., 1988), GR (Galigniana et al., 1999; Mazaira et al., 2020), ERs (King and Greene, 1984; Moriyama et al., 2020), and classic PRG receptors (nPRs) (Guiochonmantel et al., 1992; Tyagi et al., 1998). In this study, our finding that nucleocytoplasmic shuttling of PAQR8 is a dynamic event under steroid actions in nPR(-) cells is almost identical to the behaviors of other steroid hormone receptors. These findings indicate that the currently known non-classic mPRs are a novel group of progesterone receptors with the same or similar functionalities of classic nPRs that can perform both genomic and non-genomic PRG-actions, with possibly more tissue- and cell-specific intracellular performance. Lastly, another convincing piece of evidence is that PGRMC1, a close relative to mPRs, has been localized to the nucleus and participates in nucleocytoplasmic shuttling after treatment with human chorionic gonadotropin (hCG) in rat granulosa cells (Peluso et al., 2006), behaving the same as mPRs. Since then, PGRMC1 has been widely accepted as localizing to both the nucleus and cytosol. So far, PGRMC1 cytoplasmic-nuclear trafficking has been observed in spontaneously immortalized granulosa cells (SIGCs) (Peluso et al., 2010; Peluso et al., 2008a; Peluso et al., 2008b), human granulosa cells (Engmann et al., 2006; Peluso et al., 2009), and human ovarian cancer cells (Peluso et al., 2008a; Peluso et al., 2008b). The nucleocytoplasmic shuttling was also observed (Peluso, 2007; Peluso et al., 2008a; Peluso et al., 2008b) in various cells, further validating the ability of membrane PRG receptors for nucleocytoplasmic shuttling under steroid actions.

### CmP signaling network and breast cancers

#### CmP signaling network and breast cancers

Potential involvement of mPRs in breast cancer has been suggested (Dressing and Thomas, 2007), however, due to the existence of both nPR(+/-) breast tumors and breast cancer cell lines, it is extremely challenging to define the role of PRG in breast cancer (Lange, 2008). Non-classic mPRs are widely expressed in breast cancer cell lines and biopsies, regardless of nPR (+/-) subtype (Dressing et al., 2012; Dressing and Thomas, 2007; Pang and Thomas, 2011; Xie et al., 2012; Zuo et al., 2010). Overexpression of major mPRs (PAQR7, PAQR8, and PAQR5) has been associated with breast cancer development and progression (Dressing et al., 2012; Dressing and Thomas, 2007), as well as other reproductive tumors, such as cervical and ovarian cancer (Charles et al., 2010; Romero-Sanchez et al., 2008; Thomas, 2008) and has also been detected in diverse epithelial human ovarian cancer biopsies (Romero-Sanchez et al., 2008) and the HeLa cervical cancer cell line (Thomas, 2008), suggesting a potential role for mPRs in reproductive tumor biology (Abou-Fadel et al., 2020c; Dressing and Thomas, 2007). Our data certainly provide more evidence in supporting these views (Fig. 8). Epithelial-mesenchymal transition (EMT), a key step in tumorigenesis, is often associated with BPBC cells; it was demonstrated that PRG decreases protein levels of EMT markers, such as snail and fibronectin, and increases cell-cell junction proteins, such as E-cadherin and occludin in a dose-dependent manner via mPR*α* (PAQR7) (Zuo et al., 2010). Further, mPR*α* was found to function as an essential mediator for PRG-induced inhibitory effects on cell migration and invasion in BPBC cells (Xie et al., 2015). Interestingly, mPR*α*, caveolin-1 (Cav-1), and epidermal growth factor receptor (EGFR) have been co-localized at caveolae, and PRG induces repression of EMT via a caveola bound signaling complex (Dressing and Thomas, 2007; Thomas, 2008); mPRα was also found to mediate EMT through activation of the PI3K/Akt pathway in MB468 cells, suggesting that mPRs transactivate membrane-localized receptor tyrosine kinases (RTKs) (Zuo et al., 2010).

Previous reports suggested that mPRα is not produced in response to PRG induction (Zuo et al., 2010) and there are undetectable levels of mPRα in MB231 cells (Xie et al., 2015), however, our current data demonstrated that the expression of mPRα can be induced by combined steroids in MB231 cells, at both the transcriptional and translational levels (Figs. 1A, 1D, 2D), suggesting that very subtle changes in either the CmP network or cell types could lead to very different outcomes. Similarly, the expression of CCM1 can also be induced by combined steroids in MB468 cells, at both the transcriptional and translational levels (Figs. 2D, 2E). These data suggest mPRα might contribute to PRG induced EMT suppression in both BPBC cells.

#### Different responses to sex steroid actions in TNBCs

In the era of personalized medicine or precision medicine, the correct classification of breast cancer into molecular subtypes with distinctive gene expression signatures or profiles to provide a needed prognosis for treatment is an unmet task for this remarkably heterogeneous and deadly human condition. Although nPR(-) cells lack the appropriate nuclear receptors to enact genomic responses under PRG actions, PRG can activate downstream signaling in both nPR(+/-) cells by binding to mPRs (PAQRs) (Abou-Fadel et al., 2020a; Dosiou et al., 2008; Pang and Thomas, 2011; Sleiter et al., 2009). In light of a growing number of studies that have implicated mPRs in breast cancer (Adam et al., 2003; Cancer Genome Atlas, 2012; Castelnovo et al., 2019; Charles et al., 2010; Dressing et al., 2012; Dressing and Lange, 2009; Dressing and Thomas, 2007; Jiang et al., 2016; Zhao et al., 2017), we sought to characterize the CmP signaling network in patient-derived TNBC lines. In contrast to nPR(+) T47D cells, TNBC cells are nPR(-)/mPR(+) (Dressing et al., 2012; Dressing and Thomas, 2007; Pang and Thomas, 2011). The distinct RNA/Protein expression profiles of key factors within the CmP network under steroid actions have been utilized to discover potential biomarkers to classify two TNBC cells into two subgroups (Fig. 8) based on their different responses to sex hormone actions in this study. Additional classification of more new subtypes among this specific response to sex steroid actions has added another layer of complexity in the role of steroid hormones in breast tumorigenesis, progression, and metastasis. Finally, our data support the previous findings that multiple mPRs can be co-expressed in various mammalian cell types (Dosiou et al., 2008; Dressing et al., 2012; Karteris et al., 2006; Nutu et al., 2009; Thomas and Pang, 2012) to perform multifaceted non-classic PRG signaling cascades among different nPR(+/-) mammalian cells (Dressing et al., 2012).

#### Potential implication for cancer therapy

PRG plays a critical role in the progression of breast, ovarian and brain cancer (Cabrera-Munoz et al., 2011; Hennessy et al., 2009; Lange and Yee, 2008). PRG regulates specific functions in nPR(-) cancer cells that implicates mPRs in cancer initiation and progression (Xie et al., 2012; Zuo et al., 2010). Earlier reports showed that PRG promotes cell proliferation, migration, and invasion of glioblastoma cells both *in-vitro* (German-Castelan et al., 2014; Gonzalez-Aguero et al., 2007; Pina-Medina et al., 2016) and *in-vivo* (German-Castelan et al., 2016; Gonzalez-Aguero et al., 2007; Pina-Medina et al., 2016). Data also showed that PRG promotes cell proliferation through induced activation of MAPK signaling in both nPR(+/-) breast cancer cell lines, independent from nPRs (Wiebe et al., 2016), suggesting mPRs may play some role in this signaling pathway. However, reports also showed that PRG also displays a dual effect on apoptosis induction in ovarian cancer cells via mPRs and nPRs mechanisms (Diep et al., 2015; Peluso et al., 2002). Activation of mPRs increases proliferation, migration and invasion of glioblastoma cells (Gonzalez-Orozco et al., 2018), and PRG-induced mPR activity can also enhance β1/β2-adrenergic receptor (βARs) -mediated increases in cAMP levels (Charles et al., 2010), limiting proliferation in response to other signals in nPR(-) ovarian cancer cell lines (Charles et al., 2010).

As a well-known antagonist of PRG, MIF has been regarded as a great candidate for the treatment of breast, prostate, ovarian, endometrial cancers, and endometriosis (Bakker et al., 1990; Lanari et al., 2012) and is currently on many active interventional clinical trials (Goyeneche and Telleria, 2015; Kamaraju et al., 2021; Klijn et al., 1994). Some data demonstrated that a growth inhibitory agent, elevated levels of MIF, can enhance the growth inhibition and induction of apoptosis triggered by high doses of progesterone in either nPR(+/-) cancer cells (Fjelldal et al., 2010; Moe et al., 2009). Other reports showed that MIF significantly improves treatment efficacy of chemotherapy regimens for human ovarian carcinoma cell lines (Gamarra-Luques et al., 2012) and was able to inhibit the growth of nPR(-) MB231 breast cancer cells (Liang et al., 2003), suggesting MIF has anti-tumor effects on gynecological-related tumor cells (Sang et al., 2018).

Although anti-proliferative activity of MIF in cancer cells is independent of nPRs (Bardon et al., 1987; Tieszen et al., 2011), the cellular effects of MIF on proliferation is controversial, reported as both a growth inhibitor (Bardon et al., 1985) as well as a stimulator (Bowden et al., 1989; Skildum et al., 2005), depending on cell types (Bardon et al., 1985; Bowden et al., 1989; Goyeneche et al., 2012), dosage, and perhaps even existence or ratio of mPRs/nPRs (Hissom and Moore, 1987; Louie and Sevigny, 2017). Interestingly, MIF works synergistically with high dose PRG to inhibit endometrial cancer cell growth (Moe et al., 2009) and anticancer properties of MIF are known as an endocrine-related phenomenon (Fjelldal et al., 2010). In summary, the previous contradicting data with PRG, MIF, or a combination of both might all reflect one thing in common, the final cellular effects and outcomes are dependent on cell types, dosage, and existence or ratio of mPRs/nPRs in breast cancer cells.

Our data of PRG/MIF actions on two nPR(-) TNBC cells are unique and interesting. PRG displayed no major change in cell migration in both AAW-TNBC and CAW-TNBC cells, while MIF appeared to enhance cell migration in both TNBC cells at early stages (Fig. 2A). The combined steroid effect was in-between the individual responses only in AAW-TNBC cells also at early stages (Fig. 2A, bottom panel). Similarly, combined steroids appeared to enhance wound closure ability only in AAW-TNBC cells as well (Fig. 2B, bottom panel), suggesting that PRG/MIF might not be good anti-cancer candidate drugs to be used in AAW-TNBC patients. With newly emerging mPR specific agonists with higher binding affinities (Kelder et al., 2010), mechanistic studies to uncover the key roles and cellular relationship of key players in the CmP network among various breast cancer cell types will be timely and exciting (Tieszen et al., 2011).

#### Potential prognostic biomarkers for AAW-TNBCs

Biomarkers have been widely used for breast cancer therapies and offer precise treatment based on tumor and patient characteristics by utilizing almost all available current cutting-edge biomedical technologies (genetics, epigenetics, genomics, proteomics, etc.) (Fasching et al., 2015). Biomarkers can be used as a prognostic tool to predict the future course of the disease and patient response to the designated therapy (Schmidt et al., 2012). The treatment of metastatic breast cancer has become more complicated due to the increasing number of new therapies necessary to treat the elevated number of various histological, molecular, and cellular subgroups. More biomarkers are urgently needed to assist these new therapies, such as the treatment of patients with/without BRCA mutations (breast and ovarian cancers) (Schneeweiss et al., 2018; Taran et al., 2018). The difference among breast cancer subtypes is quite large in terms of clinical relevance, patterns of gene expression, selection of therapeutic strategies, responses to treatment, and prognosis (Andre et al., 2019; Colomer et al., 2018; Krop et al., 2017a; Krop et al., 2017b; Liang et al., 2019; Liu et al., 2016; Vieira and Schmitt, 2018).Therefore, identification of new biomarkers for specific breast cancer subtypes is important in guiding treatment decisions and predicting prognosis (Wu and Chu, 2021).

With our experience in multi-omics and systems biology analysis, we utilized AAW-derived TNBC cells, to discover a total of 47 potential biomarkers synchronously regulated at both the transcriptional and translational levels, associated with tumorigenesis in AAW-derived TNBCs (Suppl. Table 6D). Utilizing publicly available clinical data of expression and prognosis, we filtered these 47 biomarkers down to 21 clinically-relevant AAW-TNBC specific biomarkers with differential expression and significant survival curves (Table 1) resulting in two groups of biomarkers; intrinsic and inducible biomarkers for AAW-TNBCs (Table 1). Despite previous reports of differentially expressed cellular markers between TNBC basal subtypes (BaA-specific vs BaB-specific) (Dai et al., 2017; Lehmann et al., 2011; Neve et al., 2006; Toft and Cryns, 2011), our data show that sex steroid hormone-induced differential expression patterns among AAW-TNBCs is not overlapped with any previous data and is unrelated to the reported differentially expressed cellular markers in TNBC subtypes, making our newly identified biomarkers great candidates for AAW-TNBC clinical applications. The identified potential AAW-associated TNBC biomarkers (Table 1) will be the first important tool for breast cancer therapy/prevention in AAW-TNBC patients, a pioneering first step to tackle the currently alarming Black-White disparity in breast cancer mortality.

## Conclusion

With very limited knowledge, there has been much speculation regarding the relationship between classic PRG actions via nPRs and non-classic PRG actions via mPRs (PAQRs). It has been reported that increased expression of mPRs often leads to activation of nuclear response elements and transcriptional changes, suggesting an integrated model of sex steroid receptor signaling where steroid-dependent early (non-genomic) effects can contribute to late (genomic) nuclear actions (Boonyaratanakornkit et al., 2018). However, this integrated model is only viable in nPR(+) cells regarding mPRs-PRG-nPRs (mPn) signaling. In our previous study, we provided strong evidence that the CSC couples both classic PRG receptors (nPRs) and non-classic PRG receptors (mPRs/PAQRs) to form the CmPn signaling network in nPR(+) breast cancer cells (Abou-Fadel et al., 2020b). In this report, we extended our findings that the CSC coordinately works with non-classic PRG receptors (mPRs/PAQRs) to form the CmP signaling network in nPR(-) breast cancer cells (Fig. 8). Our discovery that mPRß is predominantly a nuclear protein and capable of shuttling between the nucleus and cytosol under steroid actions suggests that mPRs are a new type of PRG receptors, behaving the same as classic nPRs that can perform both genomic and non-genomic PRG-mediated actions. Based on these novel findings, we provide proof of principle that the CmP signaling network is a core component of our previously defined CmPn signaling network (Abou-Fadel et al., 2020b). To emphasize one of the most exciting findings in this study, utilizing comprehensive omic approaches, we identified a total of 21 AAW-TNBC specific biomarkers (Table 1) associated with tumorigenesis of AAW-derived TNBC cells. Further validation of these biomarkers among AAW-TNBC patients would have a tremendous impact on future cancer therapy and health disparities in this highly aggressive breast cancer group in African American women.

## Materials and Methods

### Cell culture, treatments and performance assays

#### Cell culture and treatments

Multiple nPR(-) cell lines (MDA-MB-231, MDA-MB-468, MCF10A, and Jurkat) were cultured in RPMI1640 medium following manufacturer’s instructions (ATCC); when cells reached 80% confluency, cells were starved with FBS-free RPMI1640 medium for 3 hrs, then treated with either vehicle control (ethanol/DMSO, VEH), mifepristone (MIF, 20 µM), progesterone (PRG, 20 µM), or combined steroids (MIF+PRG, 20 µM each). For RNA knockdown experiments, 80% confluent selected cells were transfected with a set of siRNAs, targeting specific genes (Suppl. Table 1), by RNAiMAX (Life Technologies) as described before (Abou-Fadel et al., 2020c; Jiang et al., 2019). After cells were treated in various conditions, they were harvested for RNA and protein extraction to measure the expression levels of *CCMs* and *mPRs* genes as described before (Abou-Fadel et al., 2020c; Jiang et al., 2019).

#### Cell migration Assay

Cell migration was assessed using scratch assays. Briefly, a cell scratch was pressed through the confluent TNBC (triple negative breast cancer cells, MB231, MB468) cell monolayer in the plate to mark the starting line. The cells were swept away on one side of that line. Vehicle control (EtOH, DMSO), Mifepristone (MIF, 20 µM), progesterone (PRG, 20 µM), or combined steroids (MIF+PRG, 20 μM each) were added to start the experiments. Migration was monitored temporally. The migration area was visualized using a Nikon Biostation and recorded with a high-resolution digital camera. The migration area was measured from four different fields under 20× magnifications for each condition, time-point and cell type. Each experiment was repeated three times. We calculated the rate of cell migration (R_M_) according to the equation: RM=(Wi-Wf)/t, where W_i_ is the average of the initial scratch width, W_f_ is the average of the final scratch width and t is the time span of the assay.

#### Wound healing assay

Two TNBC cells (MB231, MB468) were cultured in 24-well plates (5×10^4^ cells/well) to reach confluency. Then, cells were starved in FBS-free medium for 3 hrs, then a 100 µl pipette tip was used to scratch the monolayer of confluent cells to create a middle wound groove. Cells were cultured in triplicates in FBS-free medium in the vehicle control (EtOH, DMSO), mifepristone (MIF, 20 µM), progesterone (PRG, 20 µM), or combined steroids (MIF+PRG, 20 μM each). The wound areas were visualized using a Nikon Biostation and recorded with a high-resolution digital camera. The wound area was measured from five different fields under 20× magnifications for each condition, time-point and cell type. Each experiment was repeated three times. The wound area, wound coverage of total area, as well as average and standard deviation of the scratch width was performed with the aid of an imageJ plugin (Suarez-Arnedo et al., 2020). We calculated the percentage of wound closure with the following equation: Wound Closure % = (A_t=0_-A_t=Δt_/At=0)*100, where *A_t_* _= 0_ is the initial wound area, *A_t = Δt_* is the wound area after n hours of the initial scratch, both in μm^2^.

#### Cell Invasion assay

The migration of two TNBC cells (MB231, MB468) were performed using a 24-well transwell chambers (8 µm pore size; Millipore, Billerica, MA, USA). The cells were plated in triplicates into the upper chamber (1 × 10^5^ cells/well) with 1640 (no FBS) in the presence of combined steroids (MIF+PRG, 20 μM each) or vehicle alone for 48 hrs at 37°C, 5% CO_2_ conditions. The bottom chambers were supplemented with 1640 containing 10% FBS. Non-invading cells remaining on the upper surface of the membrane were removed using a cotton swab, and the invading cells on the bottom surface were stained with Diff-Quick stain kit (Fisher). Invading cells were visualized using a 40× magnification Nikon microscope and photographed with a high-resolution digital camera. The invasion value was quantified from the mean number of five different fields per condition. Each experiment was repeated three times.

### Immunofluorescence (IF) of CAW-TNBC/AAW-TNBC cells under steroid actions

#### Growth of TNBC cells using chamber slides for IF applications

IF staining methods were performed as previously described (Abou-Fadel et al., 2020b). Briefly, cells were grown to 80% confluency and then treated with MIF+PRG (20 µM each) over time in glass chamber slides (Nunc-Lab-Tek II) and fixed using 4% (*w/v*) Paraformaldehyde (PFA). Slides were washed 3X with PBT (0.2% Triton X-100) before proceeding with antigen retrieval step.

#### Antigen Retrieval

Slides were submerged in 10mM sodium citrate buffer (Na_3_C_6_H_5_O_7_, pH 6.0) containing 0.01% Triton X-100 at 95-98° degrees Celsius (°C). Slides were kept in citrate buffer at 95-98°C for 30 minutes (mins) and then allowed to cool down to room temperature (RT).

#### Blocking and Antibody incubation

After antigen retrieval, slides were briefly incubated in PBS containing % Triton X-100 for 10 mins at RT to permeabilize cells after antigen retrieval. Following permeabilization, cells were blocked with Pierce fast blocking buffer (Fisher) for 2 hours at RT. CCM1-Alexafluor® 488 (1:50), CCM3-Alexafluor® 546 (1:50), and PAQR8 (1:50) antibodies were diluted into PBS buffer containing 2.0% BSA and 0.2% Triton X-100. 200 µl was placed directly in staining chamber to incubate for 4 hrs at RT in the dark with gentle agitation. Secondary antibody was added for PAQR8 stained slides only using mouse anti-rabbit IgG-CFL 555 (1:100). All antibodies used are detailed in Suppl. table 2.

#### Mounting/Sealing

Immunofluorescence slides are DAPI stained during the mounting/sealing process (Suppl. Table 2). To allow efficient staining of DAPI, slides should be allowed to rest O/N at 4 °C in the dark before sealing with nail polish to cure O/N.

#### Imaging and Quantification

Imaging was performed using a Nikon Eclipse Ti confocal microscope using a 60X objective lens. Quantification was done automatically using Elements Analysis software provided with the Nikon microscope. Thresholding (used for quantification) was defined and maintained throughout all images for each application to ensure no bias was applied to data and to exclude low and high outliers. Using Nikon elements analysis tools, localization of CCM1/CCM3/PAQR8 was performed using binary operations to identify the relative ratio of expressed proteins that localized inside the nucleus (overlaps of DAPI/GFP or mCherry) compared to expressed proteins that localized in the cytosol (unique GFP or mCherry without overlapping DAPI signal). Fluorescent images are quantified for CCM1 using wavelength channel 488nm while CCM3 and PAQR8 were quantified using wavelength channel 555nm.

### RNA extraction, RT-qPCR, and RNA-seq for TNBC cells

Total RNAs were extracted with TRIZOL reagent (Invitrogen) following the manufacturer’s protocol. For cultured cells, monolayer was rinsed with ice cold PBS. Cells were lysed directly in a culture dish by adding 1 ml of TRIZOL reagent per flask and scraping with a cell scraper. The cell lysate was passed several times through a pipette and vortexed thoroughly. The quality (purity and integrity) of each RNA sample was assessed using a Bioanalyzer (Agilent) before downstream applications. All RNA-seq data were produced using Illumina HiSeq 2000; clean reads for all samples were over 99.5%; 60-80% of reads were mapped to respective reference genomes.

#### Real Time Quantitative PCR analysis (RT-qPCR)

Real-time quantitative PCR (RT-qPCR) assays were designed using primer sets (Suppl. Table 3) and applied to quantify the RNA levels of the endogenously expressed CCMs (1, 2, 3) and mPRs (PAQR5, 6, 7, 8, 9) using Power SYBR Green Master Mix with ViiA 7 Real-Time PCR System (Applied Biosystems). RT-qPCR plates with various cell-lines were prepared using an epMotion 5075 automated liquid handling system (Eppendorf). The RT-qPCR data were analyzed with DataAssist (ABI) and Rest 2009 software (Qiagen). The relative expression level (2^-ΔCT^) was calculated from all samples and normalized to a reference gene (β-actin); fold change (2^-ΔΔCT^) comparison was performed by further normalizing to control groups (Livak and Schmittgen, 2001). All experiments were performed with triplicates.

#### RNA-seq processing of files to assemble interactomes for TNBC cells

The RNA-seq files were obtained through paired-end (PE) sequencing with 100 bp reads (2X100) in Illumina HiSeq2000. The data consisted of 8 FASTQ files, 2 PE FASTQ files for each of the four groups: 468_Veh, 468_MP treated 12hrs, 468_MP treated 24hrs and 468_MP treated 48hrs. All cohorts consisted of two samples. RNA data were cleaned, and passed quality control before starting the analysis. After extracting the FASTQ files, the RNA-seq FASTQ files were aligned to the Human Genome Build 38 (GRCh38) using HISAT2 to generate two SAM files per cohort that were converted to SAM using SAMtools 1.9 and SAMtools quick check was then used to ensure the files were complete. The SAM files were then converted to BAM files, sorted and Cufflinks was used to assemble transcripts for each cohort and cuffmerge to merge all the transcript files. These files were analyzed for differential gene expression with CuffDiff. A Python script was created to identify shared differentially expressed genes between cohorts. These were annotated in Excel, and simple set operations were used to find exclusions between the sets. The overlaps were inputted into the PANTHER (Mi et al., 2021) classification system (GeneONTOLOGY) as well as iDEP (Ge et al., 2018) (Integrated Differential Expression and Pathway Analysis) program to build signaling networks with altered KEGG pathways, GO cellular components, GO molecular functions and GO biological processes.

### Protein extraction, Western blots and proteomics analysis for TNBC cells

#### Protein Extraction and quality assessment

TNBC cell lines were harvested and lysed using a digital sonifier cell disruptor (Branson model 450 with model 102C converter and double step microtip) in ice cold lysis buffer containing 50 mM Tris-HCl (pH 7.5), 150 mM NaCl, 0.5% NP-40 (Sigma), 50 mM sodium fluoride (Sigma), 1 mM PMSF (Sigma), 1 mM dithiothreitol (Invitrogen) and 1 EDTA-free complete protease inhibitor tablet (Roche). The concentration of protein lysates was measure by Qubit assay (Invitrogen) before analysis. For proteomics analysis, proteins were prepped using the filter-aided sample preparation (FASP) method (Expedeon, San Diego, CA). Samples were reduced with 10 mM DTT for 30 min at RT, and centrifuged on a spin filter at 14,000× g for 15 min at RT, and washed twice with 8 M urea in 50 mM Tris-HCl buffer. Samples were then washed 3× with 50 mM ammonium bicarbonate. Finally, samples were digested with trypsin (Sigma-Aldrich, St. Louis, MO) and peptides were eluted using 0.1% formic acid.

#### Western Blots

The relative expression levels of candidate proteins were measured with Western blots (WB). Equal amount of protein lysates from different treatments were loaded into Criterion precast gels for SDS-PAGE gel electrophoresis and transferred onto PVDF membranes at 4 ^0^C, then probed with confirmed antibodies (Suppl. Table 2) as described before (Abou-Fadel et al., 2020a; Abou-Fadel et al., 2020b; Abou-Fadel et al., 2020c; Jiang et al., 2019; Peluso et al., 2008b).

#### Liquid Chromatography-Tandem Mass Spectrometry (LC-MS/MS)

The cell lysates were generated from the four cohorts, 468_Veh, 468_MP treated 12hrs, 468_MP treated 24hrs and 468_MP treated 72hrs. All cohorts consisted of three samples. The cell lysates (20-25 µg of protein) were subjected to trypsin digestion using the FASP method, following the manufacturer’s protocol (Expedion Inc, San Diego, Ca). Four microliters of each digested sample (100 ng/µl) was loaded onto a 25-cm custom-packed porous silica Aqua C18, 125Å, 5 µm (Phenomenex) column. The porous silica was packed into a New Objective PicoTip Emitter, PF360-100-15-N-5, 15 ± 1 µm and pre-equilibrated with 95% solvent A (100% water, 0.1% formic acid) and 5% solvent B (90% acetonitrile, 10% water, 0.1% formic acid) before injection of the digested peptides, respectively. Liquid Chromatography (LC) separation of the peptides was performed on an Ultimate 3000 Dionex RSLC-nano UHPLC (Thermo Fisher Scientific), equilibrated with 95% solvent A and 5% solvent B (equilibration solution). Samples were loaded onto the column for 10 min, at a constant flow rate of 0.5µL/min, before beginning the elution gradient. Solvent B was then increased from 5% to 35% over 85 min, followed by increase to 95% solvent B over 5 min. The plateau was maintained at 95% solvent B for 9 min, followed by a sharp decrease to 5% solvent B over 1 min. The column was then re-equilibrated with 5% solvent B for 10 min. The total runtime consisted of 120 min. Peptides were analyzed using a Q Exactive Plus Hybrid Quadrupole-Orbitrap Mass Spectrometer (Thermo Fisher Scientific), equipped with a Nanospray Flex Ion Source (Thermo Fischer Scientific). Parameters for the mass spectrometer were as follows: full MS; resolution was set at 70,000 and 17,500, for MS1 and MS2, respectively; AGC target was set at 3e6 and 1e5 for MS1 and MS2, respectively; Max IT at 50 ms; scan range from 350 to 1600 m/z dd-MS2; Max IT at 100 ms; isolation window at 3.0.

#### Proteomics processing of files for TNBC cells

Proteomic data analysis was performed with Proteome Discoverer (PD) 2.1.1.21 (Thermo Fisher Scientific), using an estimated false-discovery rate (FDR) of 1%. The Human Database was downloaded in FASTA format on July, 1, 2020, from UniProtKB; http://www.uniprot.org/;177,661 entries. Common contaminants such as trypsin autolysis fragments, human keratins, and protein lab standards were included in the contaminants database (Mellacheruvu et al., 2013). The following parameters were used in the PD: HCD MS/MS; fully tryptic peptides only; up to 2 missed cleavages; parent-ion mass tolerance of 10 ppm (monoisotopic); and fragment mass tolerance of 0.6 Da (in Sequest) and 0.02 Da (in PD 2.1.1.21) (monoisotopic). A filter of two-high confidence peptides per protein were applied for identifications. PD dataset was further processed through Scaffold Q+ 4.8.2 (Proteome Software, Portland, OR). A protein threshold of 99%, peptide threshold of 95%, and a minimum number of 2 peptides were used for protein validation. Proteomic samples were analyzed via two statistical methods, Students *t-test* and ANOVA. This was done to determine changes between two specific time points (*t-test*) and changes between all-time points (ANOVA). A cutoff of p≤0.05 was executed to determine significance in all comparisons.

#### Processing of proteomic files to assemble interactomes for TNBC cells

A Python script was created to identify shared differentially expressed proteins between cohorts. These were annotated in Excel, and simple set operations were used to find exclusions between the sets. A scoring system of proteins identified in at least 3 groups were used to improve data validation. Samples that have 0 identified proteins or 0 Log2FC were proteins that were not identified in the analysis. Sample regulation over 12hrs, 24hrs and 72hrs were processed to determine trends in proteomic expression. Expression patterns were grouped to their respective fold change categories. The overlaps were inputted into the PANTHER (Mi et al., 2021) classification system (GeneONTOLOGY) as well as iDEP (Ge et al., 2018) program to build signaling networks with altered KEGG pathways, GO cellular components, GO molecular functions and GO biological processes.

#### Omics analysis to assemble interactomes for TNBC cells at both the transcriptional and translational levels

A Python script was created to identify shared differentially expressed genes/proteins between cohorts. These were annotated in Excel, and simple set operations were used to find exclusions between the sets. The overlaps were inputted into the PANTHER (Mi et al., 2021) classification system (GeneONTOLOGY) as well as iDEP (Ge et al., 2018) program to build signaling networks with altered KEGG pathways, GO cellular components, GO molecular functions and GO biological processes. Identified overlaps were compared between proteomics and RNA-seq data. The goal was to identify similarities between cohorts at both the transcriptional and translational levels. The agreements in differential expression were noted through the use of the Python comparison script.

#### Meta-analysis of CHIP-seq data overlaps with AAW-TNBC cells under steroid actions

Four databases were used (Xi et al., 2018), from various breast cancer cells (containing MDA-MB-468 cells) treated with either PRG/MIF to determine potential domain and binding domains (Xi et al., 2018). NCBI Batch Web CD-Search tool was used to determine additional details of the domain and binding domains from the database (Marchler-Bauer and Bryant, 2004; Marchler-Bauer et al., 2011). Additionally, 11 publicly available breast cancer datasets from University of California-Santa Cruz (UCSC) Xena Browser (Chin et al., 2006; Desmedt et al., 2007; Goldman et al., 2020; Heiser et al., 2012; Hess et al., 2006; Miller et al., 2005; Naderi et al., 2007; Neve et al., 2006; van’t Veer et al., 2002; van de Vijver et al., 2002; Wang et al., 2005; Yau et al., 2010) as well as two TCGA-BRCA databases available in Xena Browser (TCGA-BRCA and GDC-TCGA-BRCA) were compiled to validate differential expression among Breast cancer subtypes for our Identified candidate biomarkers. Proteomic and RNA matching to the curated domain/binding domain databases as well as the UCSC datasets was carried out using general python scripts. Gene names were used as identifiers when processing overlaps between RNA and Proteomic Analysis. Expression patterns of up or down regulation were integrated to determine transcriptional and proteomic relationships.

### Prognostic effects for identified candidate biomarkers associated with a perturbed CmP signaling network

#### Differential expression of candidate biomarkers using Microarray expression data

Expression analysis was performed as previously described (Abou-Fadel et al., 2020b). Briefly, microarray data containing breast cancer tumors, with Progesterone Receptor (PR) status determined by IHC, were analyzed using kmplot software (Gyorffy et al., 2010). This resulted in 925 PR(-)and 926 PR(+) breast cancer samples that were used to obtain expression data for the candidate biomarkers identified in this study.

#### Differential expression of candidate biomarkers using RNA-seq data for AAW-TNBCs and CAW-TNBCs

To evaluate basal expression of our identified candidate biomarkers in AAW and CAW TNBCs, we utilized publicly available RNA-seq data comparing expression levels in AAW-TNBCs and CAW-TNBCs (Saleh et al., 2021). This cohort contained 23 TNBC-AAW samples, and 19 TNB-CAW samples that were used to obtain expression data for the candidate biomarkers identified in this study.

#### Construction of Kaplan-Meier (KM) survival curves for identified biomarkers

KM survival curves were generated as previously described (Gyorffy et al., 2010). Briefly, publicly available microarray data (22,277 probes) from 1,809 breast cancer patients was analyzed using kmplot software (Gyorffy et al., 2010) integrating gene expression and clinical data simultaneously. Breast cancer patients were filtered to only analyze patient samples classified as ER(-)/PR(-)/HER2(-)/TNBC subtype, which reduced our initial 1,809 patients to either 176/392 patient samples (depending on biomarker analyzed). Logrank P values were calculated by the software (Gyorffy et al., 2010) as well as hazard ratio (and 95% confidence intervals).

### Statistical Analysis

<u>For RT-qPCR analysis</u>, the relative RNA expression changes of CmP network genes were measured by qPCR (Fold changes). All pairwise multiple comparison procedures were analyzed using Tukey and Student’s *t-test*. <u>For Western blots</u>, the relative expression levels of CmP network proteins were measured through quantification of band intensities and normalized against α-actinin (ACTN1) and vehicle controls (VEH). All pairwise multiple comparison procedures were analyzed using Tukey and Student’s *t-test*. <u>For cell migration/invasion and wound healing assays</u>, two-way analysis of variance (ANOVA) was used to detect the differences in the mean temporal values among the treatment groups in migration and wound healing assays. For invasion assays, all pairwise multiple comparison procedures were analyzed using Tukey and Student’s *t-test*. <u>For immunofluorescence analysis</u>, both one-way (analysis of a single time-point compared to control) and two-way (analysis of multiple time-points compared to controls) analysis of variance (ANOVA) was used to detect the differences in the mean values among the treatment groups. <u>For transcriptomics analysis</u>, all pairwise multiple comparison procedures were analyzed using Tukey and Student’s *t-test*. <u>For proteomics analysis</u>, one-way analysis of variance (ANOVA) was used to detect the differences in the mean values among the treatment groups. All pairwise multiple comparison procedures were analyzed using Tukey and Student’s un-paired *t-test*. <u>For microarray/RNA-seq analysis,</u> Statistical significance was performed with students *t*-test. All graphs, plots and charts were constructed and produced by SigmaPlot 12.0 (Systat Software, Inc.) and GraphPad Prism 8.

## Supporting information

Suppl Mat

## Abbreviations

(TNBC): Triple-Negative Breast Cancer
(BPBC): Basal Phenotype Breast Cancers
(BaA): Basal A
(BaB): Basal B
(CSC): CCM signaling complex
(PRG): progesterone
(MIF): Mifepristone
(nPRs): nuclear Progesterone Receptors
(mPRs/PAQRs): membrane Progesterone Receptors or Progestin and AdipoQ Receptors
(PAQR7): mPRα
(PAQR8): mPRβ
(PAQR5): mPRγ
(PAQR9): mPRε
(PGRMC1): Progesterone Receptor Membrane Component 1
(AAW): African American Women
(CAW): Caucasian American Women
(CmPn): CSC-mPRs-PRG-nPRs
(CmP): CSC-mPRs-PRG
(GR): Glucocorticoid Receptor
(AR): Androgen Receptor
(GPCRs): G-protein Coupled Receptors
(DEGs): Differentially Expressed Genes
(DEPs): Differentially Expressed Proteins
(GnRH): Gonadotropin-Releasing Hormone

## Acknowledgments

We wish to thank Kamran Falahati, Mark Smith, Khalid Shoukat, Deepak Muthyala, Mike Yao, Yanchun Qu, Shen Sheng, Ahmed Badr, Junli Zhang, Amna Siddiqui, Pallavi Dubey, Saafan Malik, and Edna Lopez at Texas Tech University Health Science Center El Paso (TTUHSCEP) for their technical help during the experiments. The results shown here are in part based upon data generated by the TCGA Research Network: https://www.cancer.gov/tcga

## Author contributions

JZ: Conceptualization, Methodology, Investigation, Writing-Original draft preparation, Writing-Reviewing and Editing; JAF: Investigation, Visualization, Software, Data curation, Validation, Writing-Reviewing and Editing, XTJ: Investigation, Visualization; BG: Software, Data curation, Validation; AP: Investigation, Writing-Original draft preparation; CCE: Data curation, Validation; EF and AMDCDLO: Investigation, Visualization.

## References

Abou-Fadel, J., Jiang, X., Padarti, A., Goswami, D., Marchuk, D., Walker, W., Grajeda, B., and Zhang, J. (2020a). CCM signaling complex (CSC) is a master regulator governing homeostasis of progesterone and its mediated signaling cascades. bioRxiv.

Abou-Fadel, J., Jiang, X., Padarti, A., Grajeda, B., and Zhang, J. (2020b). CCM signaling complex (CSC) coupling both classic and non-classic progesterone receptor signaling bioRxiv.

Abou-Fadel, J., Qu, Y., Gonzalez, E., Smith, M., and Zhang, J. (2020c). Emerging roles of CCM genes during tumorigenesis with potential application as novel biomarkers across major types of cancers. ONCOLOGY REPORTS.

Adam, P. J., Boyd, R., Tyson, K. L., Fletcher, G. C., Stamps, A., Hudson, L., Poyser, H. R., Redpath, N., Griffiths, M., Steers, G., et al. (2003). Comprehensive proteomic analysis of breast cancer cell membranes reveals unique proteins with potential roles in clinical cancer. J Biol Chem 278, 6482–6489.

Allicock, M., Graves, N., Gray, K., and Troester, M. A. (2013). African American women’s perspectives on breast cancer: implications for communicating risk of basal-like breast cancer. J Health Care Poor Underserved 24, 753–767.

Anastas, J. N., and Moon, R. T. (2013). WNT signalling pathways as therapeutic targets in cancer. Nature Reviews Cancer 13, 11–26.

Andre, F., Ismaila, N., Henry, N. L., Somerfield, M. R., Bast, R. C., Barlow, W., Collyar, D. E., Hammond, M. E., Kuderer, N. M., Liu, M. C., et al. (2019). Use of Biomarkers to Guide Decisions on Adjuvant Systemic Therapy for Women With Early-Stage Invasive Breast Cancer: ASCO Clinical Practice Guideline Update-Integration of Results From TAILORx. J Clin Oncol 37, 1956–1964.

Areia, A., Vale-Pereira, S., Alves, V., Rodrigues-Santos, P., Moura, P., and Mota-Pinto, A. (2015). Membrane progesterone receptors in human regulatory T cells: a reality in pregnancy. BJOG 122, 1544–1550.

Ashley, R. L., Clay, C. M., Farmerie, T. A., Niswender, G. D., and Nett, T. M. (2006). Cloning and characterization of an ovine intracellular seven transmembrane receptor for progesterone that mediates calcium mobilization. Endocrinology 147, 4151–4159.

Babyshkina, N. N., Dronova, T. A., Zambalova, E. A., Zavyalova, M. V., Slonimskaya, E. M., and Cherdyntseva, N. V. (2020). The role of epidermal growth factor receptor (EGFR) in the efficacy of neoadjuvant chemotherapy in triple-negative breast cancer patients. Byulleten Sib Med 19, 13–20.

Bakker, G. H., Setyono-Han, B., Portengen, H., De Jong, F. H., Foekens, J. A., and Klijn, J. G. (1990). Treatment of breast cancer with different antiprogestins: preclinical and clinical studies. J Steroid Biochem Mol Biol 37, 789–794.

Balkwill, F. (2004). Cancer and the chemokine network. Nature Reviews Cancer 4, 540–550.

Baquet, C. R., Mishra, S. I., Commiskey, P., Ellison, G. L., and DeShields, M. (2008). Breast cancer epidemiology in blacks and whites: disparities in incidence, mortality, survival rates and histology. J Natl Med Assoc 100, 480–488.

Bardon, S., Vignon, F., Chalbos, D., and Rochefort, H. (1985). RU486, a progestin and glucocorticoid antagonist, inhibits the growth of breast cancer cells via the progesterone receptor. J Clin Endocrinol Metab 60, 692–697.

Bardon, S., Vignon, F., Montcourrier, P., and Rochefort, H. (1987). Steroid receptor-mediated cytotoxicity of an antiestrogen and an antiprogestin in breast cancer cells. Cancer Res 47, 1441–1448.

Bauer, K. R., Brown, M., Cress, R. D., Parise, C. A., and Caggiano, V. (2007). Descriptive analysis of estrogen receptor (ER)-negative, progesterone receptor (PR)-negative, and HER2-negative invasive breast cancer, the so-called triple-negative phenotype: a population-based study from the California cancer Registry. Cancer-Am Cancer Soc 109, 1721–1728.

Bergh, J. J., Lin, H. Y., Lansing, L., Mohamed, S. N., Davis, F. B., Mousa, S., and Davis, P. J. (2005). Integrin alpha(v)beta(3) contains a cell surface receptor site for thyroid hormone that is linked to activation of mitogen-activated protein kinase and induction of angiogenesis. Endocrinology 146, 2864–2871.

Beverly, L. N., Flanders, W. D., Go, R. C., and Soong, S. J. (1987). A comparison of estrogen and progesterone receptors in black and white breast cancer patients. Am J Public Health 77, 351–353.

Boonyaratanakornkit, V., Hamilton, N., Marquez-Garban, D. C., Pateetin, P., McGowan, E. M., and Pietras, R. J. (2018). Extranuclear signaling by sex steroid receptors and clinical implications in breast cancer. Mol Cell Endocrinol 466, 51–72.

Bowden, R. T., Hissom, J. R., and Moore, M. R. (1989). Growth stimulation of T47D human breast cancer cells by the anti-progestin RU486. Endocrinology 124, 2642–2644.

Boyer-Chammard, A., Taylor, T. H., and Anton-Culver, H. (1999). Survival differences in breast cancer among racial/ethnic groups: a population-based study. Cancer Detect Prev 23, 463–473.

Cabrera-Munoz, E., Hernandez-Hernandez, O. T., and Camacho-Arroyo, I. (2011). Role of progesterone in human astrocytomas growth. Curr Top Med Chem 11, 1663–1667.

Cancer Genome Atlas, N. (2012). Comprehensive molecular portraits of human breast tumours. Nature 490, 61–70.

Carey, L. A., Perou, C. M., Livasy, C. A., Dressler, L. G., Cowan, D., Conway, K., Karaca, G., Troester, M. A., Tse, C. K., Edmiston, S., et al. (2006). Race, breast cancer subtypes, and survival in the Carolina Breast Cancer Study. JAMA 295, 2492–2502.

Castelnovo, L. F., Magnaghi, V., and Thomas, P. (2019). Expression of membrane progesterone receptors (mPRs) in rat peripheral glial cell membranes and their potential role in the modulation of cell migration and protein expression. Steroids 142, 6–13.

Chacon, R. D., and Costanzo, M. V. (2010). Triple-negative breast cancer. Breast Cancer Research 12.

Charles, N. J., Thomas, P., and Lange, C. A. (2010). Expression of membrane progesterone receptors (mPR/PAQR) in ovarian cancer cells: implications for progesterone-induced signaling events. Horm Cancer 1, 167-176.

Chen, L., and Li, C. I. (2015). Racial disparities in breast cancer diagnosis and treatment by hormone receptor and HER2 status. Cancer Epidemiol Biomarkers Prev 24, 1666–1672.

Chen, V. W., Correa, P., Kurman, R. J., Wu, X. C., Eley, J. W., Austin, D., Muss, H., Hunter, C. P., Redmond, C., Sobhan, M., and, et al. (1994). Histological characteristics of breast carcinoma in blacks and whites. Cancer Epidemiol Biomarkers Prev 3, 127–135.

Chin, K., DeVries, S., Fridlyand, J., Spellman, P. T., Roydasgupta, R., Kuo, W. L., Lapuk, A., Neve, R. M., Qian, Z. W., Ryder, T., et al. (2006). Genomic and transcriptional aberrations linked to breast cancer pathophysiologies. Cancer Cell 10, 529–541.

Chow, M. T., and Luster, A. D. (2014). Chemokines in Cancer. Cancer Immunol Res 2, 1125–1131.

Colomer, R., Aranda-Lopez, I., Albanell, J., Garcia-Caballero, T., Ciruelos, E., Lopez-Garcia, M. A., Cortes, J., Rojo, F., Martin, M., and Palacios-Calvo, J. (2018). Biomarkers in breast cancer: A consensus statement by the Spanish Society of Medical Oncology and the Spanish Society of Pathology. Clin Transl Oncol 20, 815–826.

Dai, X. F., Cheng, H. Y., Bai, Z. H., and Li, J. (2017). Breast Cancer Cell Line Classification and Its Relevance with Breast Tumor Subtyping. J Cancer 8, 3131–3141.

Danforth, D. N., Jr. (2013). Disparities in breast cancer outcomes between Caucasian and African American women: a model for describing the relationship of biological and nonbiological factors. Breast Cancer Res 15, 208.

Desmedt, C., Piette, F., Loi, S., Wang, Y., Lallemand, F., Haibe-Kains, B., Viale, G., Delorenzi, M., Zhang, Y., d’Assignies, M. S., et al. (2007). Strong time dependence of the 76-gene prognostic signature for node-negative breast cancer patients in the TRANSBIG multicenter independent validation series. Clin Cancer Res 13, 3207–3214.

Diep, C. H., Daniel, A. R., Mauro, L. J., Knutson, T. P., and Lange, C. A. (2015). Progesterone action in breast, uterine, and ovarian cancers. J Mol Endocrinol 54, R31–53.

Dignam, J. J. (2000). Differences in breast cancer prognosis among African-American and Caucasian women. CA Cancer J Clin 50, 50–64.

Dosiou, C., Hamilton, A. E., Pang, Y., Overgaard, M. T., Tulac, S., Dong, J., Thomas, P., and Giudice, L. C. (2008). Expression of membrane progesterone receptors on human T lymphocytes and Jurkat cells and activation of G-proteins by progesterone. J Endocrinol 196, 67–77.

Dressing, G. E., Alyea, R., Pang, Y., and Thomas, P. (2012). Membrane progesterone receptors (mPRs) mediate progestin induced antimorbidity in breast cancer cells and are expressed in human breast tumors. Horm Cancer 3, 101–112.

Dressing, G. E., and Lange, C. A. (2009). Integrated actions of progesterone receptor and cell cycle machinery regulate breast cancer cell proliferation. Steroids 74, 573–576.

Dressing, G. E., and Thomas, P. (2007). Identification of membrane progestin receptors in human breast cancer cell lines and biopsies and their potential involvement in breast cancer. Steroids 72, 111–116.

Engmann, L., Losel, R., Wehling, M., and Peluso, J. J. (2006). Progesterone regulation of human granulosa/luteal cell viability by an RU486-independent mechanism. J Clin Endocr Metab 91, 4962–4968.

Esquivel-Velazquez, M., Ostoa-Saloma, P., Palacios-Arreola, M. I., Nava-Castro, K. E., Castro, J. I., and Morales-Montor, J. (2015). The Role of Cytokines in Breast Cancer Development and Progression. J Interf Cytok Res 35, 1–16.

Fasching, P. A., Brucker, S. Y., Fehm, T. N., Overkamp, F., Janni, W., Wallwiener, M., Hadji, P., Belleville, E., Haberle, L., Taran, F. A., et al. (2015). Biomarkers in Patients with Metastatic Breast Cancer and the PRAEGNANT Study Network. Geburtshilfe Frauenheilkd 75, 41–50.

Fernandes, M. S., Brosens, J. J., and Gellersen, B. (2008). Honey, we need to talk about the membrane progestin receptors. Steroids 73, 942–952.

Fernandes, M. S., Pierron, V., Michalovich, D., Astle, S., Thornton, S., Peltoketo, H., Lam, E. W. F., Gellersen, B., Huhtaniemi, I., Allen, J., and Brosens, J. J. (2005). Regulated expression of putative membrane progestin receptor homologues in human endometrium and gestational tissues. Journal of Endocrinology 187, 89–101.

Filardo, E. J. (2002). Epidermal growth factor receptor (EGFR) transactivation by estrogen via the G-protein-coupled receptor, GPR30: a novel signaling pathway with potential significance for breast cancer. J Steroid Biochem Mol Biol 80, 231-238.

Fjelldal, R., Moe, B. T., Orbo, A., and Sager, G. (2010). MCF-7 cell apoptosis and cell cycle arrest: non-genomic effects of progesterone and mifepristone (RU-486). Anticancer Res 30, 4835–4840.

Foster, H., Reynolds, A., Stenbeck, G., Dong, J., Thomas, P., and Karteris, E. (2010). Internalisation of membrane progesterone receptor-alpha after treatment with progesterone: Potential involvement of a clathrin-dependent pathway. Mol Med Rep 3, 27–35.

Freese, J. L., Pino, D., and Pleasure, S. J. (2010). Wnt signaling in development and disease. Neurobiology of Disease 38, 148–153.

Fu, X., Liang, C., Li, F., Wang, L., Wu, X., Lu, A., Xiao, G., and Zhang, G. (2018). The Rules and Functions of Nucleocytoplasmic Shuttling Proteins. Int J Mol Sci 19.

Galigniana, M. D., Housley, P. R., DeFranco, D. B., and Pratt, W. B. (1999). Inhibition of glucocorticoid receptor nucleocytoplasmic shuttling by okadaic acid requires intact cytoskeleton. Journal of Biological Chemistry 274, 16222–16227.

Gamarra-Luques, C. D., Goyeneche, A. A., Hapon, M. B., and Telleria, C. M. (2012). Mifepristone prevents repopulation of ovarian cancer cells escaping cisplatin-paclitaxel therapy. BMC Cancer 12, 200.

Gao, J. J., Aksoy, B. A., Dogrusoz, U., Dresdner, G., Gross, B., Sumer, S. O., Sun, Y. C., Jacobsen, A., Sinha, R., Larsson, E., et al. (2013). Integrative Analysis of Complex Cancer Genomics and Clinical Profiles Using the cBioPortal. Sci Signal 6.

Ge, S. X., Son, E. W., and Yao, R. (2018). iDEP: an integrated web application for differential expression and pathway analysis of RNA-Seq data. Bmc Bioinformatics 19, 534.

Gerend, M. A., and Pai, M. (2008). Social determinants of Black-White disparities in breast cancer mortality: a review. Cancer Epidemiol Biomarkers Prev 17, 2913–2923.

German-Castelan, L., Manjarrez-Marmolejo, J., Gonzalez-Arenas, A., and Camacho-Arroyo, I. (2016). Intracellular Progesterone Receptor Mediates the Increase in Glioblastoma Growth Induced by Progesterone in the Rat Brain. Arch Med Res 47, 419–426.

German-Castelan, L., Manjarrez-Marmolejo, J., Gonzalez-Arenas, A., Gonzalez-Moran, M. G., and Camacho-Arroyo, I. (2014). Progesterone induces the growth and infiltration of human astrocytoma cells implanted in the cerebral cortex of the rat. Biomed Res Int 2014, 393174.

Goel, H. L., Breen, M., Zhang, J. Z., Das, I., Aznavoorian-Cheshire, S., Greenberg, N. M., Elgavish, A., and Languino, L. R. (2005). beta(1A) integrin expression is required for type 1 insulin-like growth factor receptor mitogenic and transforming activities and localization to focal contacts. Cancer Research 65, 6692–6700.

Goldhirsch, A., Wood, W. C., Coates, A. S., Gelber, R. D., Thurlimann, B., Senn, H. J., and Panel, m. (2011). Strategies for subtypes--dealing with the diversity of breast cancer: highlights of the St. Gallen International Expert Consensus on the Primary Therapy of Early Breast Cancer 2011. Ann Oncol 22, 1736-1747.

Goldman, M. J., Craft, B., Hastie, M., Repecka, K., McDade, F., Kamath, A., Banerjee, A., Luo, Y., Rogers, D., Brooks, A. N., et al. (2020). Visualizing and interpreting cancer genomics data via the Xena platform. Nat Biotechnol 38, 675–678.

Gonzalez-Aguero, G., Gutierrez, A. A., Gonzalez-Espinosa, D., Solano, J. D., Morales, R., Gonzalez-Arenas, A., Cabrera-Munoz, E., and Camacho-Arroyo, I. (2007). Progesterone effects on cell growth of U373 and D54 human astrocytoma cell lines. Endocrine 32, 129–135.

Gonzalez-Orozco, J. C., Hansberg-Pastor, V., Valadez-Cosmes, P., Nicolas-Ortega, W., Bastida-Beristain, Y., Fuente-Granada, M., Gonzalez-Arenas, A., and Camacho-Arroyo, I. (2018). Activation of membrane progesterone receptor-alpha increases proliferation, migration, and invasion of human glioblastoma cells. Mol Cell Endocrinol 477, 81–89.

Goyeneche, A. A., Seidel, E. E., and Telleria, C. M. (2012). Growth inhibition induced by antiprogestins RU-38486, ORG-31710, and CDB-2914 in ovarian cancer cells involves inhibition of cyclin dependent kinase 2. Invest New Drugs 30, 967-980.

Goyeneche, A. A., and Telleria, C. M. (2015). Antiprogestins in gynecological diseases. Reproduction 149, R15–33.

Guiochonmantel, A., Lescop, P., Christinmaitre, S., Perrotapplanat, M., and Milgrom, E. (1992). Nucleocytoplasmic Shuttling of the Progesterone-Receptor. Ann Biol Clin-Paris 50, 387–392.

Guiochonmantel, A., and Milgrom, E. (1993). Cytoplasmic Nuclear Trafficking of Steroid-Hormone Receptors. Trends Endocrin Met 4, 322–328.

Gyorffy, B., Lanczky, A., Eklund, A. C., Denkert, C., Budczies, J., Li, Q. Y., and Szallasi, Z. (2010). An online survival analysis tool to rapidly assess the effect of 22,277 genes on breast cancer prognosis using microarray data of 1,809 patients. Breast Cancer Research and Treatment 123, 725–731.

Hall, I. J., Moorman, P. G., Millikan, R. C., and Newman, B. (2005). Comparative analysis of breast cancer risk factors among African-American women and White women. Am J Epidemiol 161, 40–51.

Heikkinen, K., Rapakko, K., Karppinen, S. M., Erkko, H., Knuutila, S., Lundan, T., Mannermaa, A., Borresen-Dale, A. L., Borg, A., Barkardottir, R. B., et al. (2006). RAD50 and NBS1 are breast cancer susceptibility genes associated with genomic instability. Carcinogenesis 27, 1593–1599.

Heiser, L. M., Sadanandam, A., Kuo, W. L., Benz, S. C., Goldstein, T. C., Ng, S., Gibb, W. J., Wang, N. J., Ziyad, S., Tong, F., et al. (2012). Subtype and pathway specific responses to anticancer compounds in breast cancer. P Natl Acad Sci USA 109, 2724–2729.

Hennessy, B. T., Coleman, R. L., and Markman, M. (2009). Ovarian cancer. Lancet 374, 1371–1382.

Hess, K. R., Anderson, K., Symmans, W. F., Valero, V., Ibrahim, N., Mejia, J. A., Booser, D., Theriault, R. L., Buzdar, A. U., Dempsey, P. J., et al. (2006). Pharmacogenomic predictor of sensitivity to preoperative chemotherapy with paclitaxel and fluorouracil, doxorubicin, and cyclophosphamide in breast cancer. J Clin Oncol 24, 4236–4244.

Hissom, J. R., and Moore, M. R. (1987). Progestin effects on growth in the human breast cancer cell line T-47D--possible therapeutic implications. Biochem Biophys Res Commun 145, 706–711.

Ide, H., Inoue, S., and Miyamoto, H. (2018). The Role of Glucocorticoid Receptor Signaling in Bladder Cancer Progression. Cancers (Basel) 10.

Jiang, G., Zhang, S., Yazdanparast, A., Li, M., Pawar, A. V., Liu, Y., Inavolu, S. M., and Cheng, L. (2016). Comprehensive comparison of molecular portraits between cell lines and tumors in breast cancer. BMC Genomics 17 *Suppl 7*, 525.

Jiang, X., Padarti, A., Qu, Y., Sheng, S., Abou-Fadel, J., Badr, A., and Zhang, J. (2019). Alternatively spliced isoforms reveal a novel type of PTB domain in CCM2 protein. Sci Rep 9, 15808.

Josefsberg Ben-Yehoshua, L., Lewellyn, A. L., Thomas, P., and Maller, J. L. (2007). The role of Xenopus membrane progesterone receptor beta in mediating the effect of progesterone on oocyte maturation. Mol Endocrinol 21, 664–673.

Joslyn, S. A. (1995). Racial differences in survival from breast cancer. JAMA 273, 1000.

Joslyn, S. A. (2002). Hormone receptors in breast cancer: racial differences in distribution and survival. Breast Cancer Res Treat 73, 45–59.

Kamaraju, S., Fowler, A. M., Weil, E., Wisinski, K. B., Truong, T. H., Lehr, M., Chaudhary, L. N., Cheng, Y. C., Chitambar, C., Rui, H., et al. (2021). Leveraging Antiprogestins in the Treatment of Metastatic Breast Cancer. Endocrinology.

Kappler, C. S., Guest, S. T., Irish, J. C., Garrett-Mayer, E., Kratche, Z., Wilson, R. C., and Ethier, S. P. (2015). Oncogenic signaling in amphiregulin and EGFR-expressing PTEN-null human breast cancer. Mol Oncol 9, 527–543.

Karteris, E., Zervou, S., Pang, Y., Dong, J., Hillhouse, E. W., Randeva, H. S., and Thomas, P. (2006). Progesterone signaling in human myometrium through two novel membrane G protein-coupled receptors: potential role in functional progesterone withdrawal at term. Mol Endocrinol 20, 1519–1534.

Kasubuchi, M., Watanabe, K., Hirano, K., Inoue, D., Li, X., Terasawa, K., Konishi, M., Itoh, N., and Kimura, I. (2017). Membrane progesterone receptor beta (mPRbeta/Paqr8) promotes progesterone-dependent neurite outgrowth in PC12 neuronal cells via non-G protein-coupled receptor (GPCR) signaling. Sci Rep 7, 5168.

Kelder, J., Azevedo, R., Pang, Y., de Vlieg, J., Dong, J., and Thomas, P. (2010). Comparison between steroid binding to membrane progesterone receptor alpha (mPRalpha) and to nuclear progesterone receptor: correlation with physicochemical properties assessed by comparative molecular field analysis and identification of mPRalpha-specific agonists. Steroids 75, 314–322.

Kesler, C. T., Weber, M. J., and Paschal, B. M. (2004). Nucleocytoplasmic shuttling of the androgen receptor is critical for transactivation. Mol Biol Cell 15, 265a-266a.

King, W. J., and Greene, G. L. (1984). Monoclonal antibodies localize oestrogen receptor in the nuclei of target cells. Nature 307, 745–747.

Klijn, J. G., Setyono-Han, B., Sander, H. J., Lamberts, S. W., de Jong, F. H., Deckers, G. H., and Foekens, J. A. (1994). Pre-clinical and clinical treatment of breast cancer with antiprogestins. Hum Reprod 9 *Suppl 1*, 181–189.

Klopocki, E., Kristiansen, G., Wild, P. J., Klaman, I., Castanos-Velez, E., Singer, G., Stohr, R., Simon, R., Sauter, G., Leibiger, H., et al. (2004). Loss of SFRP1 is associated with breast cancer progression and poor prognosis in early stage tumors. International Journal of Oncology 25, 641–649.

Krietsch, T., Fernandes, M. S., Kero, J., Losel, R., Heyens, M., Lam, E. W. F., Huhtaniemi, I., Brosens, J. J., and Gellersen, B. (2006). Human homologs of the putative G protein-coupled membrane progestin receptors (mPR alpha, beta, and gamma) localize to the endoplasmic reticulum and are not activated by progesterone. Molecular Endocrinology 20, 3146–3164.

Krop, I., Ismaila, N., Andre, F., Bast, R. C., Barlow, W., Collyar, D. E., Hammond, M. E., Kuderer, N. M., Liu, M. C., Mennel, R. G., et al. (2017a). Use of Biomarkers to Guide Decisions on Adjuvant Systemic Therapy for Women With Early-Stage Invasive Breast Cancer: American Society of Clinical Oncology Clinical Practice Guideline Focused Update. J Clin Oncol 35, 2838–2847.

Krop, I., Ismaila, N., and Stearns, V. (2017b). Use of Biomarkers to Guide Decisions on Adjuvant Systemic Therapy for Women With Early-Stage Invasive Breast Cancer: American Society of Clinical Oncology Clinical Practice Focused Update Guideline Summary. J Oncol Pract 13, 763–766.

Lanari, C., Wargon, V., Rojas, P., and Molinolo, A. A. (2012). Antiprogestins in breast cancer treatment: are we ready? Endocr Relat Cancer 19, R35–50.

Lange, C. A. (2008). Challenges to defining a role for progesterone in breast cancer. Steroids 73, 914–921.

Lange, C. A., and Yee, D. (2008). Progesterone and breast cancer. Womens Health (Lond) 4, 151–162.

Lee, E., Lee, T. A., Yoo, H. J., Lee, S., and Park, B. (2019). CNBP controls tumor cell biology by regulating tumor-promoting gene expression. Mol Carcinog 58, 1492–1501.

Lehmann, B. D., Bauer, J. A., Chen, X., Sanders, M. E., Chakravarthy, A. B., Shyr, Y., and Pietenpol, J. A. (2011). Identification of human triple-negative breast cancer subtypes and preclinical models for selection of targeted therapies. Journal of Clinical Investigation 121, 2750–2767.

Lemale, J., Bloch-Faure, M., Grimont, A., El Abida, B., Imbert-Teboul, M., and Crambert, G. (2008). Membrane progestin receptors alpha and gamma in renal epithelium. Bba-Mol Cell Res 1783, 2234–2240.

Li, R., Peng, C., Zhang, X. Z., Wu, Y. W., Pan, S. D., and Xiao, Y. C. (2017). Roles of Arf6 in cancer cell invasion, metastasis and proliferation. Life Sciences 182, 80–84.

Liang, X., Li, H., Coussy, F., Callens, C., and Lerebours, F. (2019). An update on biomarkers of potential benefit with bevacizumab for breast cancer treatment: Do we make progress? Chin J Cancer Res 31, 586–600.

Liang, Y. Y., Hou, M., Kallab, A. M., Barrett, J. T., El Etreby, F., and Schoenlein, P. V. (2003). Induction of antiproliferation and apoptosis in estrogen receptor negative MDA-231 human breast cancer cells by mifepristone and 4-hydroxytamoxifen combination therapy: A role for TGF beta I. International Journal of Oncology 23, 369–380.

Liu, F., Gu, L. N., Shan, B. E., Geng, C. Z., and Sang, M. X. (2016). Biomarkers for EMT and MET in breast cancer: An update. Oncol Lett 12, 4869–4876.

Liu, H., Rigamonti, D., Badr, A., and Zhang, J. (2010). Ccm1 assures microvascular integrity during angiogenesis. Transl Stroke Res 1, 146–153.

Liu, H., Rigamonti, D., Badr, A., and Zhang, J. (2011). Ccm1 regulates microvascular morphogenesis during angiogenesis. J Vasc Res 48, 130–140.

Livak, K. J., and Schmittgen, T. D. (2001). Analysis of relative gene expression data using real-time quantitative PCR and the 2(T)(-Delta Delta C) method. Methods 25, 402–408.

Louie, M. C., and Sevigny, M. B. (2017). Steroid hormone receptors as prognostic markers in breast cancer. Am J Cancer Res 7, 1617–1636.

Lund, M. J., Trivers, K. F., Porter, P. L., Coates, R. J., Leyland-Jones, B., Brawley, O. W., Flagg, E. W., O’Regan, R. M., Gabram, S. G., and Eley, J. W. (2009). Race and triple negative threats to breast cancer survival: a population-based study in Atlanta, GA. Breast Cancer Res Treat 113, 357–370.

Ma, H., Ursin, G., Xu, X., Lee, E., Togawa, K., Duan, L., Lu, Y., Malone, K. E., Marchbanks, P. A., McDonald, J. A., et al. (2017). Reproductive factors and the risk of triple-negative breast cancer in white women and African-American women: a pooled analysis. Breast Cancer Res 19, 6.

Marchler-Bauer, A., and Bryant, S. H. (2004). CD-Search: protein domain annotations on the fly. Nucleic Acids Research 32, W327–W331.

Marchler-Bauer, A., Lu, S., Anderson, J. B., Chitsaz, F., Derbyshire, M. K., DeWeese-Scott, C., Fong, J. H., Geer, L. Y., Geer, R. C., Gonzales, N. R., et al. (2011). CDD: a Conserved Domain Database for the functional annotation of proteins. Nucleic Acids Res 39, D225–229.

Masuda, H., Zhang, D. W., Bartholomeusz, C., Doihara, H., Hortobagyi, G. N., and Ueno, N. T. (2012). Role of epidermal growth factor receptor in breast cancer. Breast Cancer Res Tr 136, 331–345.

Mazaira, G. I., Echeverria, P. C., and Galigniana, M. D. (2020). Nucleocytoplasmic shuttling of the glucocorticoid receptor is influenced by tetratricopeptide repeat-containing proteins. Journal of Cell Science 133.

McCullough, M. L., Feigelson, H. S., Diver, W. R., Patel, A. V., Thun, M. J., and Calle, E. E. (2005). Risk factors for fatal breast cancer in African-American women and White women in a large US prospective cohort. Am J Epidemiol 162, 734–742.

McGee, A. M., Douglas, D. L., Liang, Y. Y., Hyder, S. M., and Baines, C. P. (2011). The mitochondrial protein C1qbp promotes cell proliferation, migration and resistance to cell death. Cell Cycle 10, 4119–4127.

Mellacheruvu, D., Wright, Z., Couzens, A. L., Lambert, J. P., St-Denis, N. A., Li, T., Miteva, Y. V., Hauri, S., Sardiu, M. E., Low, T. Y., et al. (2013). The CRAPome: a contaminant repository for affinity purification-mass spectrometry data. Nature Methods 10, 730-+.

Mi, H. Y., Ebert, D., Muruganujan, A., Mills, C., Albou, L. P., Mushayamaha, T., and Thomas, P. D. (2021). PANTHER version 16: a revised family classification, tree-based classification tool, enhancer regions and extensive API. Nucleic Acids Research 49, D394–D403.

Miller, L. D., Smeds, J., George, J., Vega, V. B., Vergara, L., Ploner, A., Pawitan, Y., Hall, P., Klaar, S., Liu, E. T., and Bergh, J. (2005). An expression signature for p53 status in human breast cancer predicts mutation status, transcriptional effects, and patient survival (vol 102, pg 13550, 2005). P Natl Acad Sci USA 102, 17882-17882.

Moe, B. G., Vereide, A. B., Orbo, A., and Sager, G. (2009). High concentrations of progesterone and mifepristone mutually reinforce cell cycle retardation and induction of apoptosis. Anticancer Res 29, 1053–1058.

Moriyama, T., Yoneda, Y., Oka, M., and Yamada, M. (2020). Transportin-2 plays a critical role in nucleocytoplasmic shuttling of oestrogen receptor-alpha. Sci Rep-Uk 10.

Nader, N., Dib, M., Hodeify, R., Courjaret, R., Elmi, A., Hammad, A. S., Dey, R., Huang, X. Y., and Machaca, K. (2021). Membrane progesterone receptor induces meiosis in Xenopus oocytes through endocytosis into signaling endosomes and interaction with APPL1 and Akt2 (vol 18, e3000901, 2020). Plos Biol 19.

Naderi, A., Teschendorff, A. E., Barbosa-Morais, N. I., Pinder, S. E., Green, A. R., Powe, D. G., Robertson, J. F. R., Aparicio, S., Ellis, I. O., Brenton, J. D., and Caldas, C. (2007). A gene-expression signature to predict survival in breast cancer across independent data sets. Oncogene 26, 1507–1516.

Neve, R. M., Chin, K., Fridlyand, J., Yeh, J., Baehner, F. L., Fevr, T., Clark, L., Bayani, N., Coppe, J. P., Tong, F., et al. (2006). A collection of breast cancer cell lines for the study of functionally distinct cancer subtypes. Cancer Cell 10, 515–527.

Newman, L. A. (2014). Breast cancer disparities: high-risk breast cancer and African ancestry. Surg Oncol Clin N Am 23, 579–592.

Newman, L. A. (2017). Breast Cancer Disparities: Socioeconomic Factors versus Biology. Ann Surg Oncol 24, 2869–2875.

Newman, L. A., and Kaljee, L. M. (2017). Health Disparities and Triple-Negative Breast Cancer in African American Women: A Review. JAMA Surg 152, 485–493.

Nutu, M., Weijdegard, B., Thomas, P., Thurin-Kjellberg, A., Billig, H., and Larsson, D. G. (2009). Distribution and hormonal regulation of membrane progesterone receptors beta and gamma in ciliated epithelial cells of mouse and human fallopian tubes. Reprod Biol Endocrinol 7, 89.

Pang, Y., Dong, J., and Thomas, P. (2015). Progesterone increases nitric oxide synthesis in human vascular endothelial cells through activation of membrane progesterone receptor-alpha. Am J Physiol Endocrinol Metab 308, E899–911.

Pang, Y., and Thomas, P. (2011). Progesterone signals through membrane progesterone receptors (mPRs) in MDA-MB-468 and mPR-transfected MDA-MB-231 breast cancer cells which lack full-length and N-terminally truncated isoforms of the nuclear progesterone receptor. Steroids 76, 921–928.

Peluso, J. J. (2007). Non-genomic actions of progesterone in the normal and neoplastic mammalian ovary. Semin Reprod Med 25, 198–207.

Peluso, J. J., Fernandez, G., Pappalardo, A., and White, B. A. (2002). Membrane-initiated events account for progesterone’s ability to regulate intracellular free calcium levels and inhibit rat granulosa cell mitosis. Biol Reprod 67, 379–385.

Peluso, J. J., Liu, X., Gawkowska, A., Lodde, V., and Wu, C. A. (2010). Progesterone inhibits apoptosis in part by PGRMC1-regulated gene expression. Molecular and Cellular Endocrinology 320, 153–161.

Peluso, J. J., Liu, X. F., Gawkowska, A., and Johnston-MacAnanny, E. (2009). Progesterone Activates a Progesterone Receptor Membrane Component 1-Dependent Mechanism That Promotes Human Granulosa/Luteal Cell Survival But Not Progesterone Secretion. J Clin Endocr Metab 94, 2644–2649.

Peluso, J. J., Liu, X. F., Saunders, M. M., Claffey, K. P., and Phoenix, K. (2008a). Regulation of ovarian cancer cell viability and sensitivity to cisplatin by progesterone receptor membrane component-1. J Clin Endocr Metab 93, 1592–1599.

Peluso, J. J., Pappalardo, A., Losel, R., and Wehling, M. (2006). Progesterone membrane receptor component 1 expression in the immature rat ovary and its role in mediating progesterone’s antiapoptotic action. Endocrinology 147, 3133–3140.

Peluso, J. J., Romak, J., and Liu, X. F. (2008b). Progesterone receptor membrane component-1 (PGRMC1) is the mediator of progesterone’s antiapoptotic action in spontaneously immortalized granulosa cells as revealed by PGRMC1 small interfering ribonucleic acid treatment and functional analysis of PGRMC1 mutations. Endocrinology 149, 534–543.

Perou, C. M., Sorlie, T., Eisen, M. B., van de Rijn, M., Jeffrey, S. S., Rees, C. A., Pollack, J. R., Ross, D. T., Johnsen, H., Akslen, L. A., et al. (2000). Molecular portraits of human breast tumours. Nature 406, 747–752.

Perrett, R. M., and McArdle, C. A. (2013). Molecular mechanisms of gonadotropin-releasing hormone signaling: integrating cyclic nucleotides into the network. Front Endocrinol (Lausanne) 4, 180.

Pina-Medina, A. G., Hansberg-Pastor, V., Gonzalez-Arenas, A., Cerbon, M., and Camacho-Arroyo, I. (2016). Progesterone promotes cell migration, invasion and cofilin activation in human astrocytoma cells. Steroids 105, 19–25.

Romero-Sanchez, M., Peiper, S. C., Evans, B., Wang, Z., Catasus, L., Ribe, A., Prat, J., and Giri, J. G. (2008). Expression profile of heptahelical putative membrane progesterone receptors in epithelial ovarian tumors. Hum Pathol 39, 1026–1033.

Saleh, M., Chandrashekar, D. S., Shahin, S., Agarwal, S., Kim, H. G., Behring, M., Shaikh, A. J., Moloo, Z., Eltoum, I. A., Yates, C., et al. (2021). Comparative analysis of triple-negative breast cancer transcriptomics of Kenyan, African American and Caucasian Women. Transl Oncol 14, 101086.

Salhi, A., Lemale, J., Paris, N., Bloch-Faure, M., and Crambert, G. (2010). Membrane progestin receptors: beyond the controversy, can we move forward? Biomol Concepts 1, 41–47.

Sang, L., Wang, X. Y., and Zhao, X. B. (2018). Mifepristone Inhibits the Migration of Cervical Cancer Cells by Inhibiting Exocrine Secretion. Pharmacology 101, 322–329.

Schally, A. V. (1999). Luteinizing hormone-releasing hormone analogs: their impact on the control of tumorigenesis. Peptides 20, 1247–1262.

Schiffenbauer, Y. S., Meir, G., Maoz, M., Even-Ram, S. C., Bar-Shavit, R., and Neeman, M. (2002). Gonadotropin stimulation of MLS human epithelial ovarian carcinoma cells augments cell adhesion mediated by CD44 and by alpha(v)-integrin. Gynecol Oncol 84, 296–302.

Schmidt, M., Fasching, P. A., Beckmann, M. W., and Kolbl, H. (2012). Biomarkers in Breast Cancer - An Update. Geburtshilfe Frauenheilkd 72, 819–832.

Schneeweiss, A., Lux, M. P., Janni, W., Hartkopf, A. D., Nabieva, N., Taran, F. A., Overkamp, F., Kolberg, H. C., Hadji, P., Tesch, H., et al. (2018). Update Breast Cancer 2018 (Part 2) - Advanced Breast Cancer, Quality of Life and Prevention. Geburtshilfe Frauenheilkd 78, 246–259.

Sharma, P. (2018). Update on the Treatment of Early-Stage Triple-Negative Breast Cancer. Curr Treat Option On 19.

Shiao, Y. H., Chen, V. W., Lehmann, H. P., Wu, X. C., and Correa, P. (1997). Patterns of DNA ploidy and S-phase fraction associated with breast cancer survival in blacks and whites. Clin Cancer Res 3, 587–592.

Skildum, A., Faivre, E., and Lange, C. A. (2005). Progesterone receptors induce cell cycle progression via activation of mitogen-activated protein kinases. Mol Endocrinol 19, 327–339.

Sleiter, N., Pang, Y., Park, C., Horton, T. H., Dong, J., Thomas, P., and Levine, J. E. (2009). Progesterone receptor A (PRA) and PRB-independent effects of progesterone on gonadotropin-releasing hormone release. Endocrinology 150, 3833–3844.

Smith, J. L., Kupchak, B. R., Garitaonandia, I., Hoang, L. K., Maina, A. S., Regalla, L. M., and Lyons, T. J. (2008). Heterologous expression of human mPR alpha, mPR beta and mPR gamma in yeast confirms their ability to function as membrane progesterone receptors. Steroids 73, 1160–1173.

Srivastava, M., Eidelman, O., Craig, J., Starr, J., Kvecher, L., Liu, J., Hueman, M., Pollard, H. B., Hu, H., and Shriver, C. D. (2019). Serum Biomarkers for Racial Disparities in Breast Cancer Progression. Mil Med 184, 652–657.

Sturtz, L. A., Melley, J., Mamula, K., Shriver, C. D., and Ellsworth, R. E. (2014). Outcome disparities in African American women with triple negative breast cancer: a comparison of epidemiological and molecular factors between African American and Caucasian women with triple negative breast cancer. BMC Cancer 14, 62.

Suarez-Arnedo, A., Torres Figueroa, F., Clavijo, C., Arbelaez, P., Cruz, J. C., and Munoz-Camargo, C. (2020). An image J plugin for the high throughput image analysis of in vitro scratch wound healing assays. PLoS One 15, e0232565.

Sun, W., Tian, B. X., Wang, S. H., Liu, P. J., and Wang, Y. C. (2020). The function of SEC22B and its role in human diseases. Cytoskeleton (Hoboken) 77, 303–312.

Tan, J. A., Joseph, D. R., Quarmby, V. E., Lubahn, D. B., Sar, M., French, F. S., and Wilson, E. M. (1988). The rat androgen receptor: primary structure, autoregulation of its messenger ribonucleic acid, and immunocytochemical localization of the receptor protein. Mol Endocrinol 2, 1276–1285.

Taran, F. A., Schneeweiss, A., Lux, M. P., Janni, W., Hartkopf, A. D., Nabieva, N., Overkamp, F., Kolberg, H. C., Hadji, P., Tesch, H., et al. (2018). Update Breast Cancer 2018 (Part 1) - Primary Breast Cancer and Biomarkers. Geburtshilfe Frauenheilkd 78, 237–245.

Thomas, P. (2008). Characteristics of membrane progestin receptor alpha (mPRalpha) and progesterone membrane receptor component 1 (PGMRC1) and their roles in mediating rapid progestin actions. Front Neuroendocrinol 29, 292–312.

Thomas, P., and Pang, Y. (2012). Membrane progesterone receptors: evidence for neuroprotective, neurosteroid signaling and neuroendocrine functions in neuronal cells. Neuroendocrinology 96, 162–171.

Thomas, P., Pang, Y. F., and Dong, J. (2014). Enhancement of Cell Surface Expression and Receptor Functions of Membrane Progestin Receptor alpha (mPR alpha) by Progesterone Receptor Membrane Component 1 (PGRMC1): Evidence for a Role of PGRMC1 as an Adaptor Protein for Steroid Receptors. Endocrinology 155, 1107-1119.

Tieszen, C. R., Goyeneche, A. A., Brandhagen, B. N., Ortbahn, C. T., and Telleria, C. M. (2011). Antiprogestin mifepristone inhibits the growth of cancer cells of reproductive and non-reproductive origin regardless of progesterone receptor expression. BMC Cancer 11, 207.

Tischkau, S. A., and Ramirez, V. D. (1993). A specific membrane binding protein for progesterone in rat brain: sex differences and induction by estrogen. Proc Natl Acad Sci U S A 90, 1285–1289.

Toft, D. J., and Cryns, V. L. (2011). Minireview: Basal-Like Breast Cancer: From Molecular Profiles to Targeted Therapies. Molecular Endocrinology 25, 199–211.

Tokumoto, T., Hossain, M. B., and Wang, J. (2016). Establishment of procedures for studying mPR-interacting agents and physiological roles of mPR. Steroids 111, 79–83.

Tyagi, R. K., Amazit, L., Lescop, P., Milgrom, E., and Guiochon-Mantel, A. (1998). Mechanisms of progesterone receptor export from nuclei: Role of nuclear localization signal, nuclear export signal, and ran guanosine triphosphate. Molecular Endocrinology 12, 1684–1695.

Valadez-Cosmes, P., Vazquez-Martinez, E. R., Cerbon, M., and Camacho-Arroyo, I. (2016). Membrane progesterone receptors in reproduction and cancer. Mol Cell Endocrinol 434, 166–175.

van’t Veer, L. J., Dai, H. Y., van de Vijver, M. J., He, Y. D. D., Hart, A. A. M., Mao, M., Peterse, H. L., van der Kooy, K., Marton, M. J., Witteveen, A. T., et al. (2002). Gene expression profiling predicts clinical outcome of breast cancer. Nature 415, 530–536.

van de Vijver, M. J., He, Y. D., van’t Veer, L. J., Dai, H., Hart, A. A. M., Voskuil, D. W., Schreiber, G. J., Peterse, J. L., Roberts, C., Marton, M. J., et al. (2002). A gene-expression signature as a predictor of survival in breast cancer. New Engl J Med 347, 1999–2009.

Veeraraghavan, J., Ma, J. C., Hu, Y. H., and Wang, X. S. (2016). Recurrent and pathological gene fusions in breast cancer: current advances in genomic discovery and clinical implications. Breast Cancer Res Tr 158, 219–232.

Vieira, A. F., and Schmitt, F. (2018). An Update on Breast Cancer Multigene Prognostic Tests-Emergent Clinical Biomarkers. Front Med (Lausanne) 5, 248.

Wang, C. H., Cheng, Y. Q., Zhang, X. P., Li, N., Zhang, L., Wang, S. D., Tong, X. M., Xu, Y., Chen, G. Q., Cheng, S. Q., et al. (2018). Vacuolar Protein Sorting 33B Is a Tumor Suppressor in Hepatocarcinogenesis. Hepatology 68, 2239–2253.

Wang, Y., Klijn, J. G., Zhang, Y., Sieuwerts, A. M., Look, M. P., Yang, F., Talantov, D., Timmermans, M., Meijer-van Gelder, M. E., Yu, J., et al. (2005). Gene-expression profiles to predict distant metastasis of lymph-node-negative primary breast cancer. Lancet 365, 671–679.

Weichert, W., Kristiansen, G., Winzer, K. J., Schmidt, M., Gekeler, V., Noske, A., Muller, B. M., Niesporek, S., Dietel, M., and Denkert, C. (2005). Polo-like kinase isoforms in breast cancer: expression patterns and prognostic implications. Virchows Arch 446, 442–450.

Wells, A. A., Gulbas, L., Sanders-Thompson, V., Shon, E. J., and Kreuter, M. W. (2014). African-American breast cancer survivors participating in a breast cancer support group: translating research into practice. J Cancer Educ 29, 619–625.

Wiebe, J. P., Pawlak, K. J., and Kwok, A. (2016). Mechanism of action of the breast cancer-promoter hormone, 5alpha-dihydroprogesterone (5alphaP), involves plasma membrane-associated receptors and MAPK activation. J Steroid Biochem Mol Biol 155, 166-176.

Wu, H. J., and Chu, P. Y. (2021). Recent Discoveries of Macromolecule- and Cell-Based Biomarkers and Therapeutic Implications in Breast Cancer. International Journal of Molecular Sciences 22.

Xi, Y. X., Shi, J. J., Li, W. Q., Tanaka, K., Allton, K. L., Richardson, D., Li, J., Franco, H. L., Nagari, A., Malladi, V. S., et al. (2018). Histone modification profiling in breast cancer cell lines highlights commonalities and differences among subtypes. Bmc Genomics 19.

Xie, M., Zhou, L., Chen, X., Gainey, L. O., Xiao, J., Nanes, M. S., Hou, A., You, S., and Chen, Q. (2015). Progesterone and Src family inhibitor PP1 synergistically inhibit cell migration and invasion of human basal phenotype breast cancer cells. Biomed Res Int 2015, 426429.

Xie, M., Zhu, X., Liu, Z., Shrubsole, M., Varma, V., Mayer, I. A., Dai, Q., Chen, Q., and You, S. (2012). Membrane progesterone receptor alpha as a potential prognostic biomarker for breast cancer survival: a retrospective study. PLoS One 7, e35198.

Yau, C., Esserman, L., Moore, D. H., Waldman, F., Sninsky, J., and Benz, C. C. (2010). A multigene predictor of metastatic outcome in early stage hormone receptor-negative and triple-negative breast cancer. Breast Cancer Research 12.

Zhang, H. R., Lai, S. Y., Huang, L. J., Zhang, Z. F., Liu, J., Zheng, S. R., Ding, K., Bai, X., and Zhou, J. Y. (2018). Myosin 1b promotes cell proliferation, migration, and invasion in cervical cancer. Gynecologic Oncology 149, 188–197.

Zhang, J., Basu, S., Clatterbuck, R. E., Rigamonti, D., and Dietz, H. C. (2004). Pathogenesis of cerebral cavernous malformation: Depletion of Krit1 leads to perturbation of 1 integrin-mediated endothelial cell mobility and survival. Am J Hum Genet suppl, S222.

Zhang, J., Basu, S., Rigamonti, D., Dietz, H. C., and Clatterbuck, R. E. (2005). Depletion of KRIT1 leads to perturbation of beta 1 integrin-mediated endothelial cell angiogenesis in the pathogenesis of cerebral cavernous malformation. Stroke 36, 425–425.

Zhang, J., Basu, S., Rigamonti, D., Dietz, H. C., and Clatterbuck, R. E. (2008). Krit1 modulates beta 1-integrin-mediated endothelial cell proliferation. Neurosurgery 63, 571–578; discussion 578.

Zhang, J., Carr, C., and Badr, A. (2011). The cardiovascular triad of dysfunctional angiogenesis. Transl Stroke Res 2, 339–345.

Zhao, Y., Ruan, X., Wang, H., Li, X., Gu, M., Wang, L., Li, Y., Seeger, H., and Mueck, A. O. (2017). The presence of a membrane-bound progesterone receptor induces growth of breast cancer with norethisterone but not with progesterone: A xenograft model. Maturitas 102, 26–33.

Zheng, J., Ali, A., and Ramirez, V. D. (1996). Steroids conjugated to bovine serum albumin as tools to demonstrate specific steroid neuronal membrane binding sites. J Psychiatry Neurosci 21, 187–197.

Zhu, Y., Bond, J., and Thomas, P. (2003a). Identification, classification, and partial characterization of genes in humans and other vertebrates homologous to a fish membrane progestin receptor. Proc Natl Acad Sci U S A 100, 2237–2242.

Zhu, Y., Hanna, R. N., Schaaf, M. J. M., Spaink, H. P., and Thomas, P. (2008). Candidates for membrane progestin receptors-Past approaches and future challenges. Comp Biochem Phys C 148, 381–389.

Zhu, Y., Rice, C. D., Pang, Y., Pace, M., and Thomas, P. (2003b). Cloning, expression, and characterization of a membrane progestin receptor and evidence it is an intermediary in meiotic maturation of fish oocytes. Proc Natl Acad Sci U S A 100, 2231–2236.

Zimmerli, D., Hausmann, G., Cantu, C., and Basler, K. (2017). Pharmacological interventions in the Wnt pathway: inhibition of Wnt secretion versus disrupting the protein-protein interfaces of nuclear factors. Br J Pharmacol 174, 4600–4610.

Zuo, L., Li, W., and You, S. (2010). Progesterone reverses the mesenchymal phenotypes of basal phenotype breast cancer cells via a membrane progesterone receptor mediated pathway. Breast Cancer Res 12, R34.

