## Supplementary material for "CmP signaling network unveils novel biomarkers for triple negative breast cancer in African American women": Suppl Mat

### CSC-mPRs-PRG (CmP) signaling network unveils novel biomarkers for triple negative breast cancer (TNBC) in African American women

Johnathan Abou-Fadel[1, *], Brian Grajeda[2,*], Xiaoting Jiang[1],
Alyssa-Marie D. Cailing-De La O[1], Esmeralda Flores[1], Akhil Padarti[1], Muaz Bhalli[1], Alexander Le[1], Jun Zhang[1**]

Department of Molecular and Translational Medicine (MTM) [1]
Texas Tech University Health Science Center El Paso, El Paso, TX 79905 USA

Department of Biological Sciences [2]
University of Texas at El Paso, El Paso, TX 79902 USA

*Running Title:* CCM signaling complex (CSC) modulates progesterone-mediated signaling in triple negative breast cancer (TNBC) cells

*Authors Contributed Equally

**All correspondence:
Jun Zhang, Sc.D., Ph.D.
Department of Molecular and Translational Medicine (MTM)
Texas Tech University Health Science Center El Paso
5001 El Paso Drive, El Paso, El Paso, TX 79905


**Table S1. Small double-stranded interfering RNAs (siRNA) used for *gene silencing* in RNA interference (RNAi) experiments.** Gene-specific knockdown was achieved by introducing small double-stranded siRNA oligos. Detailed information about application, siRNA oligo duplexes, manufacturers, and catalog numbers are listed.

**Table S2. Antibodies used in this study**. Detailed information about application, antigen, clone code, secondary antibody, manufacturer, and catalog number are listed. *IHC, Immunohistochemistry; IF, immunofluorescence; WB, Western Blots.

**Table S3. Primers for quantitative PCR (RT-qPCR) for the detection of differentially expressed targeted genes**. The primer sequence, RT-qPCR product size and optimized annealing temperature of each primer pairs are listed.

**Table S4A: RAW RNAseq FASTQ files from MB468-specific Differentially Expressed Genes (DEGs).** We generated our RNAseq data specifically for MB468 cells by removing shared similarly altered DEGs between type 2B (MB468) and 2A(MB231/Jurkat) cells. Comparison of DEGs (Column A) with technical *p*-values ≤ 0.05 from 12, 24, 48 hrs time points of TNBC cells under steroid actions were used. Data shows three time points: 12 hrs Log2F.C. from technical replicates in columns B and D (p-values in columns C and E), 24 hrs technical replicates in columns F and H (p-values in columns G and I), and 48 hrs technical replicates in columns J and L (p-values in columns K and M).

**Table S4B**: **Expression patterns in MB468-specific DEGs.** We filtered out our RNAseq data for MB468 cells by removing similarly altered DEGs shared between type 2B (MB468) and 2A (MB231/Jurkat) cells. Technical *p*-values ≤ 0.05 from 12, 24 and 48 hrs time points of TNBC cells under steroid actions were used. Comparisons were made using Log2F.C. of DEGs (Column A) from 12 hrs technical replicates in columns B and D (p-values in columns C and E), 24 hrs technical replicates in columns F and H (p-values in columns G and I), and 48 hrs technical replicates in columns J and L (p-values in columns K and M). Genes included in Fig. 4D-G analysis were found in at least of 3 of the 6 runs (column N). Each tab is separated based off differential expression of genes, using either up/down or down/up expression patterns. 0 values represent genes that were not identified in specific runs.

**Table S5A**: **RAW MS/MS data from AAW-TNBC specific Differentially Expressed Proteins (DEPs).** We filtered our MS/MS data for AAW-TNBC cells by removing similarly altered DEPs shared between type 2B (MB468) and 2A (MB231/Jurkat) cells. Comparisons were made of DEPs with a *p*-value ≤ 0.05 from 12, 24 and 72 hrs time points of AAW-TNBC cells under steroid actions. Two statistical analysis were run, ANOVA and Students *t*-test. In the *t*-test tab the Protein Name (column A), Accession Number (column B), Alternate ID (column C), Molecular Weight (column D), averaged log2fc from each time point (columns E,G,I), *p*-value for each of the corresponding comparisons (columns F,H,J), and score identifiers for how many times a protein was found in each time point (column K) are provided. In the ANOVA tab the Protein Name (column A), Accession Number (column B), Alternate ID (column C), Molecular Weight (column D), log2fc from each time point (columns E-G), ANOVA P-values (column H) and score identifiers for how many times a protein was found in each time point (column I) are provided. Proteins included in Figs. 5A-C analysis were found in at least 2 of the 3 runs (column I on the ANOVA tab, and column K on the *t*-test tab). 0 values represent genes that were not identified in specific runs.

**Table S5B**: **Expression Patterns of AAW-TNBC specific DEPs.** We generated our MS/MS data specifically for AAW-TNBC cells by removing shared similarly altered DEPs between type 2B (MB468) and 2A(MB231/Jurkat) cells. Comparisons were made of DEPs with *p*-value ≤ 0.05 from 12, 24 and 72 hrs time points of AAW-TNBC cells under steroid actions. Two statistical analysis were run, ANOVA and students *t*-test. In the *t*-test tab the Protein Name (column A), Accession Number (column B), Alternate ID (column C), Molecular Weight (column D), averaged log2fc from each time point (columns E, G, I), *p*-values for each of the corresponding comparisons (columns

F,H,J), score identifiers for how many times a protein was found in each time point (column K), and expression patterns (columns L-Q). In the ANOVA tab the Protein Name (column A), Accession Number (column B), Alternate ID (column C), Molecular Weight (column D), log2fc from each time point (columns E-G), ANOVA P-values (column H), score identifiers for how many times a protein was found in each time point (column I), and expression patterns (columns J-O). Proteins included in Figs. 5D-G analysis were found in at least 2 of the 3 averaged runs (Column I on the ANOVA tab, and Column K on the *t*-test tab). 0 values represent genes that were not identified in specific runs.

**Table S5C**: **Literature review of MB468-specific candidate biomarkers and their functions in cancer biology.** We evaluated our 47 identified candidate biomarkers and performed a comprehensive cancer biology literature search. Column A details the genes identified, column B details the supported functions of the corresponding gene in published manuscripts, and columns C-E provide the DOI of the supporting literature for the functions detailed in Column B.

**Table S6A: Overlaps in RNAseq and MS/MS from AAW-TNBC specific Differentially Expressed Genes and Proteins (DEGs/DEPs).** We filtered out our RNAseq and MS/MS data for AAW-TNBC cells by removing similarly altered DEGs/DEPs shared between type 2B (MB468) and 2A (MB231/Jurkat) cells. The RNA (transcriptomic) data was compared to both the *t*-test and ANOVA proteomic output respectively. Tab 1 provides the Log2FC as well as p-values for transcriptomic overlaps (RNASeq) with either *t*-test (Proteomics) and/or ANOVA (Proteomics). Columns A-I detail transcriptomic analysis with Column A providing the Alternate ID, Columns B, D, and F showing the averaged log2fc from each time point, Columns C, E, and G showing the p-values of each corresponding time point, Column H provides the RNA score illustrating the amount of times a transcript was found in the 6 replicates and Column I showing the overall expression trend of the 3 averaged time points (D, down; U, up). Columns K-V provide details from *t*-test proteomic analysis with Column K providing the protein name, Column L shows the accession number, Column M shows the alternate I.D., Column N shows the molecular weight, Columns O, Q, and S provide the averaged log2fc values from each time point, Columns P, R, and T provide the corresponding p-values, Column U shows score identifiers for how many times a protein was found in each replicate and Column V shows the overall expression of the corresponding protein in regards to all 3 averaged time points (D, down; U, up). Columns X-AG details the ANOVA proteomic analysis where Column X provides the protein name, Column Y and Z show the accession number and alternate I.D., respectively, Column AA provides the molecular weight, column AE provides the corresponding p-values for Log2F.C. values in Columns AB, AC and AD, while column AF shows how many times a protein was detected in the triplicate runs, and Column AG demonstrates the overall expression pattern of the corresponding protein in regards to all 3 averaged time points (D, down; U, up). Tab 2 shows only the proteomic *t*-test comparisons with the transcriptomic data. Columns A-L display *t*-test proteomic analysis where Column A shows the protein name, Column B shows the accession number, Column C shows the alternate I.D., Column D shows the molecular weight, Columns E,G,I show averaged log2fc values from each time point, Columns F,H,J show their corresponding p-values, Column K shows score identifiers for how many times a protein was found in each time point and Column L shows the overall expression of the corresponding protein in regards to all 3 averaged time points (D, down; U, up). Columns N-V details the transcriptomic analysis where Column N shows the Alternate ID, Columns O, Q, and S show the averaged log2fc values from each time point, Columns P,R,T show the p-values of each corresponding time point, Column U shows the RNA score illustrating the amount of times a transcript was found in the 6 replicate samples and Column V shows the overall expression trend of the three averaged time points (D, down; U, up). 0 values represent genes that were not identified in specific runs. Expression analysis with N/A or null are due to the inability to determine trends due to lack of patterns or low detection levels. Tab 3 shows the comparisons of the ANOVA proteomic analysis with the transcriptomic analysis. Columns A-J details the ANOVA proteomic analysis where Column A shows the protein name, Column B shows the accession number, Column C shows the alternate I.D., Column D shows the molecular weight, Column H shows corresponding p-values for Log2F.C. in Columns E-G, while column I shows how many times a protein was detected in the triplicate runs, and Column J demonstrating the overall expression patterns of the corresponding proteins in regards to all 3 averaged time points (D, down; U, up). Columns L-T detail the transcriptomic analysis where Column L shows the Alternate ID, Columns M, O, and Q provide the averaged log2fc values from each time point, Columns N, P, and R provide the p-values of each corresponding time point, Column S shows the RNA score illustrating the amount of times a transcript was found in the 6 samples and Column T shows the overall expression trend of the three averaged time points (D, down; U, up).

**Table S6B: Signaling pathways affected from MB468-specific Differentially Expressed Genes and Proteins (DEGs/DEPs).** We filtered out our RNAseq and MS/MS data for MB468 cells by removing similarly altered DEGs/DEPs shared between type 2B (MB468) and 2A (MB231/Jurkat) cells. The RNA (transcriptomic) data was compared to both the *t*-test and ANOVA proteomic output respectively. The Signaling pathways affected are detailed in Column A, along with the Number of DEGs/DEPs involved in Column B as well as the Alternate ID's for the corresponding DEGs/DEPs.

**Table S6C**: **Domain Binding Database matching in RNAseq and MS/MS from MB468-specific DEGs/DEPs.** We filtered out our RNAseq and MS/MS data for MB468 cells by removing similarly altered DEGs/DEPs shared between type 2B (MB468) and 2A (MB231/Jurkat) cells. 4 databases were used to search for promoter significance. Domain and Binding Domain searches were carried out to enrich database framework using NCBI Batch Web CD-Search tool. RNA and proteomic outputs from database matching were compared to see the overlaps. RNAseq data was compared to both the *t*-test and ANOVA proteomic output respectively along with overlaps in our working CHIP-seq database (tab 2) and displayed in Tab 1 (Summary). Column A shows the accession ID, Column B shows the alternate ID, Column C shows the protein/gene name, Column D shows the molecular weight, RNAseq DEG data (columns E-M) include technical log2fc from each time point (columns E-J) a scoring system (column K), and the total amount of positive and negative log2fcs values found in all time points (columns L-M) are provided. Additionally, columns N-S show the corresponding Proteomic data where columns N-P show the averaged log2fc values of each time point, a scoring system of the number of proteins identified in all runs (column Q) and the total amount of positive and negative log2fc values found in all time points (columns R-S); details of how the transcript data and protein data were compared, in regards to *t*-test, ANOVA analysis methods, binding domain or domain comparison from CHIP-seq database, and available regulation details in the CHIP-seq database (either a negative or positive expression-if applicable) from the literature (columns T-Z) are also provided. Tab 2 provides the compiled database searches that were used from the literature containing CHIP-seq data on breast cancer cell lines under steroid actions. Gene symbols can be found in Column A. Databases 1 and 2 illustrated subtype specific active enhancer/transcriptional states, database 3 showed signatures of active transcription states and active transcription flanking states and database 4 showed upstream regulators of subtype gene signatures of active transcription states and active transcription flanking states (column B). Database 3 and 4 were further subdivided into additional groups referring to their cell types (column C), and their regulation changes (column D). For simplicity, databases were given database keys (column E) and Expression Keys (column F), which are referenced in the comparisons with the transcript and protein data (Tab 1). Tab 3 shows the transcript matches with the CHIP-seq data (tab 2) where column A is the gene symbol, Columns B, D, F, H, J, and L provide the technical log2fc values between the three time points as well as their respective p-values (columns C, E, G, I, K, and M). Column N details how many times the gene was found altered in all replicate samples, and the database/potential subtypes it was found in the CHIP-seq data (Columns O-S, respectively). Tab 4 shows the matches of the

Proteomic ANOVA comparisons with the CHIP-seq data (tab 2), where Column A shows the protein name, Column B shows the accession number, Column C shows the alternate I.D., Column D shows the molecular weight, Columns E-G shows the averaged log2fc values from the three time points, column H shows the p-value for the ANOVA analysis, Column I refers to how many times the protein was found in all samples, and the databases/potential subtype it was found in the CHIP-seq data (columns J-N, respectively). Finally, tab 5 displays the matches of the proteomic *t*-test comparisons with the CHIP-seq data, where Column A shows the protein name, Column B shows the accession number, Column C shows the alternate I.D., Column D shows the molecular weight, Columns E, G, and I provide the averaged log2fc expression values for the three time points along with their respective p-values(Columns F, H, and J) Column K refers to how many times the protein was found in all samples, and the database and potential subtype it was found in the CHIP-seq data (columns L-P, respectively).

**Table S6D: list of 47 identified prognostic candidate biomarkers associated with clinical data in TNBC breast cancers.** Candidate biomarkers shared at both RNA/Protein levels under hormone treatments (columns 1, 2, 3), were further validated in AAW-TNBC (n=23) and CAW-TNBC (n=19) (columns 4, 5) tumor tissues. These biomarkers were further analyzed by a set of publicly available microarray data (22,277 probes) from 1,809 breast cancer samples using kmplot software to integrate gene expression and clinical data simultaneously to generate the displayed Kaplan-Meier survival curves (column 6). Candidate biomarkers were further filtered into two groups based on classic progesterone Receptor (PR) status determined by Immunohistochemistry (IHC), to evaluate clinical expression levels focusing on 925 PR(-) breast cancer samples (column 7). Our candidate biomarkers were further validated through available CHIPseq data (column 8). Red colored background indicates increased/higher expression levels, while blue colored background indicates decreased/lower expression levels. Colored biomarkers (red or blue in Column 1) indicate intrinsic biomarkers which are selected as candidate biomarkers for AAW-TNBCs, while black colored biomarkers are inducible biomarkers, like CCM1. Abbreviations: I.S., Increased Survival; D.S., Decreased Survival; N.S., No statistical significance; HIGH IN PR(-), Higher expression levels in nPR(-) breast cancer tissues; LOW IN PR(-), Lower expression levels in nPR(-) breast cancer tissues; N/A, Not Available; Y, Yes, N, No.

**Suppl. Fig. 1A. Expression levels of membrane progesterone receptors (*mPRs/PAQRs*) genes in various breast cancer cells.** RNA expression levels of mPRs (PAQR 5, 7, 8, 9 and PGRMC1) are low in most nPR(-) cells, with the exception of PAQR7 in Jurkat cells. This is in contrast to the high expression levels observed for all mPRs (PAQR 5, 7, 8, 9 and PGRMC1) that were observed in nPR(+) T47D cells. The relative RNA expression changes of major mPRs (PAQR 5, 7, 8, 9, PGRMC1) were measured by qPCR (Fold changes) (triplicates per experiment, n=3).

**Figure S1B**. **Protein expression levels of CCM3 are not influenced by combined steroid treatment**. After combined steroid treatment for 48 hrs (PRG+MIF,+), Western blots demonstrated no visual expression changes of low-abundant CCM3 protein observed in 2 different nPR(-) cell lines, MB468 and Jurkat cells, compared to vehicle control (VEH, -) (left panels), and there is no statistical difference between treatment and control (right panel). The relative expression levels of CCM3 protein was measured through quantification of band intensities and normalized against α-actinin (ACTN1) and vehicle control (VEH) (right panel red line) (n=3).

**Fig. S2: Wound healing quantification of TNBC cells under steroid actions**: Two TNBC cells (MB231, MB468) were cultured in 24-well plates ($5\times10^4$ cells/well) to reach confluency. Then, cells were starved in FBS-free medium for 3 hrs, then a 100 μl pipette tip was used to scratch the monolayer of confluent cells to create a middle wound groove. Cells were cultured in triplicates in FBS-free medium in the vehicle control (EtOH, DMSO), Mifepristone (MIF, 20 μM), progesterone (PRG, 20 μM), or combined steroids (MIF+PRG, 20 μM each). **A.** MIF appeared to enhance wound closure in MB231 cells compared to vehicle controls. **B.** MIF as well as the combined steroids (MIF+PRG), appeared to enhance wound closure in MB468 cells. The wound areas were visualized using a Nikon Biostation and recorded with a high-resolution digital camera. Wound closure was quantified by measuring the average width of the wounds using a Nikon Biostation. The wound area was measured from five different fields under 40× magnifications for each condition, time-point and cell type. Similar results were also obtained for cell migration assay (Data not shown).

**Suppl. Fig 3A: Subcellular compartmentation of membrane progesterone receptors (mPRs/PAQRs) under combined steroid actions in nPR(-) TNBC cells.** MB231/468 Cells were treated with vehicle control (ethanol/DMSO, VEH) or progesterone + mifepristone (PRG+MIF, 20 μM each) for 48 hrs and then immunohistochemically stained to localize mPRs expression. IHC approaches utilizing HRP/DAB staining revealed nuclear localization of both PAQR5 (mPRγ) and PAQR7 (mPRα) in both TNBC cells, compared to vehicle controls (n=9).

**Fig. S3B. Potential nuclear localization signal (NLS) and nuclear export signal (NES) of mPRβ (PAQR8).** One potential NLS site (A.A 10-46, highlighted with green color), and two potential NES sites (A.A. 84-91, 112-120, highlighted with red color) were predicted by cNLS Mapper and NetNES 1.1 respectively.

**Figure S4: Expression patterns of DEGs involved in enriched pathways among MDA-MB-468 cells utilizing high-throughput RNAseq.** MB468 cells were treated with MIF+PRG (20μM each) for 12, 24 and 48 hrs. Extracted DEGs were further filtered using corresponding Type 2A cells to obtain DEGs specific to MB468 cells. **A)** Expression patterns of DEGs involved in highly enriched pathways (≥40 genes). Pathway functional enrichment results are displayed for MIF+PRG treated cells. **B)** Expression patterns of DEGs involved in moderately enriched pathways (20-39 genes). Pathway functional enrichment results are displayed for MIF+PRG treated cells. **C)** Expression patterns of DEGs involved in medium enriched pathways (10-20 genes). Pathway functional enrichment results are displayed for MIF+PRG treated cells. **D)** Expression patterns of DEGs involved in medium enriched pathways (10-20 genes). Pathway functional enrichment results are displayed for MIF+PRG treated cells. **E)** Expression patterns of DEGs involved in low enriched pathways (5-10 genes). Pathway functional enrichment results are displayed for MIF+PRG treated cells. X axis represents the genes involved. Y axis represents the log2 transformed fold change of identified DEGs involved in each pathway compared to their respective vehicle controls. Coloring indicates the time points of extraction (12 hrs: blue, 24 hrs: red, 48 hrs: green). Each timepoint was run with biological duplicates and technical triplicates for treated and vehicle samples.

**Figure S5: Expression patterns of Differentially Expressed Proteins (DEPs) involved in enriched pathways among AAW-TNBC cells utilizing high-throughput proteomics.** AAW-TNBC cells were treated with combined steroids (MIF+PRG, 20µM each) for 12, 24 and 72 hrs. Extracted DEPs were further filtered using corresponding Type 2A (MB231/Jurkat) cells to obtain DEPs specific to MB468 cells. **A)** Expression patterns of Differentially Expressed Proteins (DEPs) involved in highly enriched pathways (4-6 proteins). **B)** Expression patterns of DEPs involved in moderately enriched pathways (3 proteins). **C)** Expression patterns of DEPs involved in medium enriched pathways (2 proteins). **D)** Expression patterns of DEPs involved in low enriched pathways (1 protein). All pathway functional enrichment results are displayed for MIF+PRG treated cells. X axis represents the genes involved. Y axis represents the log2 transformed fold change of identified DEPs involved in each pathway compared to their respective vehicle controls. Coloring indicates the time points of extraction (12 hrs: blue, 24 hrs: red, 72 hrs: green). Each timepoint was run with biological triplicates and technical triplicates for treated and vehicle samples.

**Figure S7. Equal basal expression of candidate biomarkers, displaying significant survival curves, for breast cancer patient samples.** Microarray data containing breast cancer tumors, analyzed using kmplot software, were divided into two groups based on Progesterone Receptor (PR) status determined by Immunohistochemistry (IHC); after filtering, there were 925 PR(-) and 926 PR(+) breast cancer samples. All panels illustrate genes that demonstrated equal expression in PR(-) breast cancer tissue samples, compared to PR(+) tissues. Additionally, all panels reflect genes that displayed significant differences in Kaplan-Meier survival curves. Statistical significance was performed with students *t*-test.

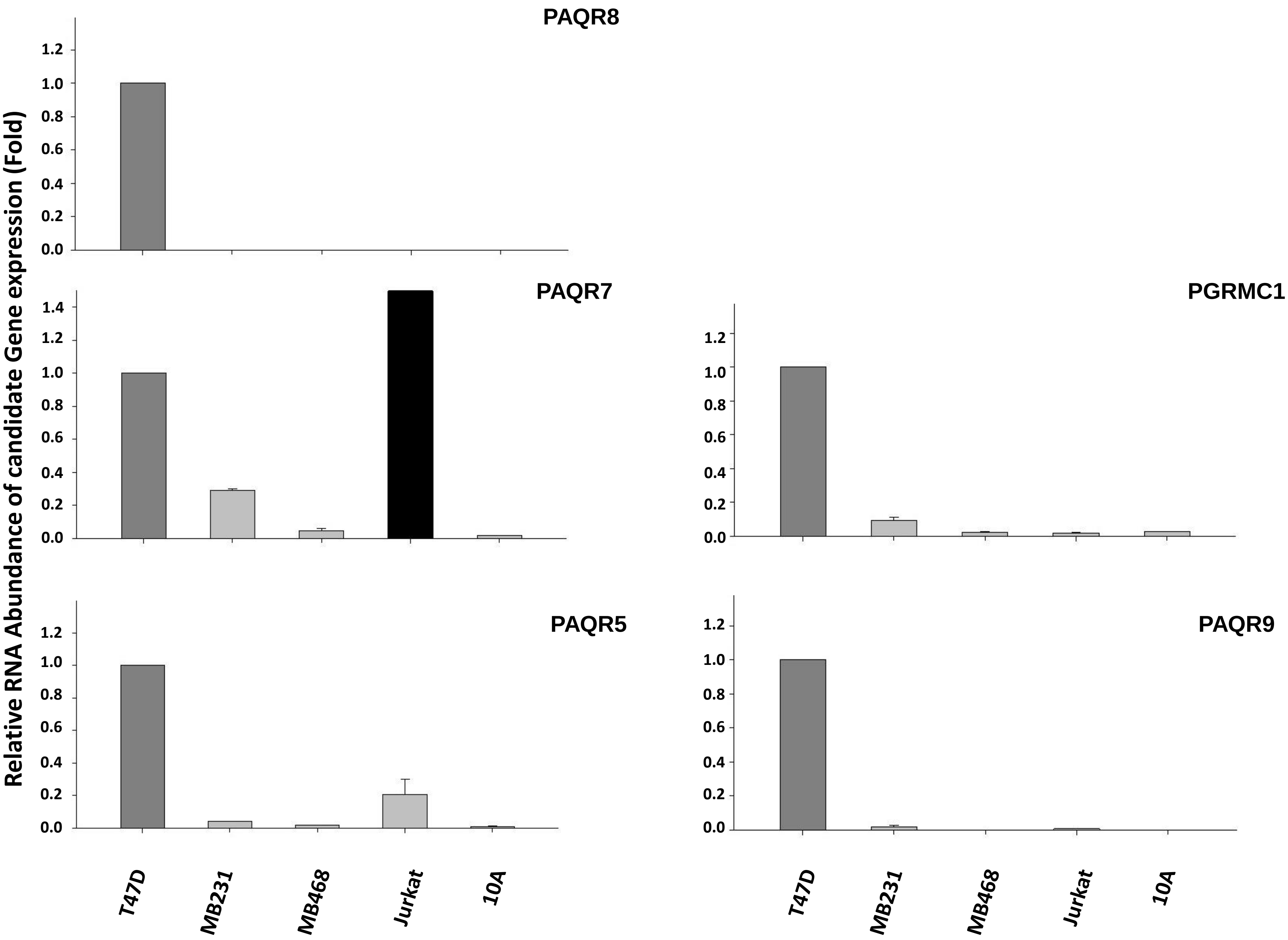
Relative RNA Abundance of candidate Gene expression (Fold)
PAQR8
PAQR7
PGRMC1
PAQR5
PAQR9
0.0
0.2
0.4
0.6
0.8
1.0
1.2
1.4
T47D
MB231
MB468
Jurkat
10A

1C

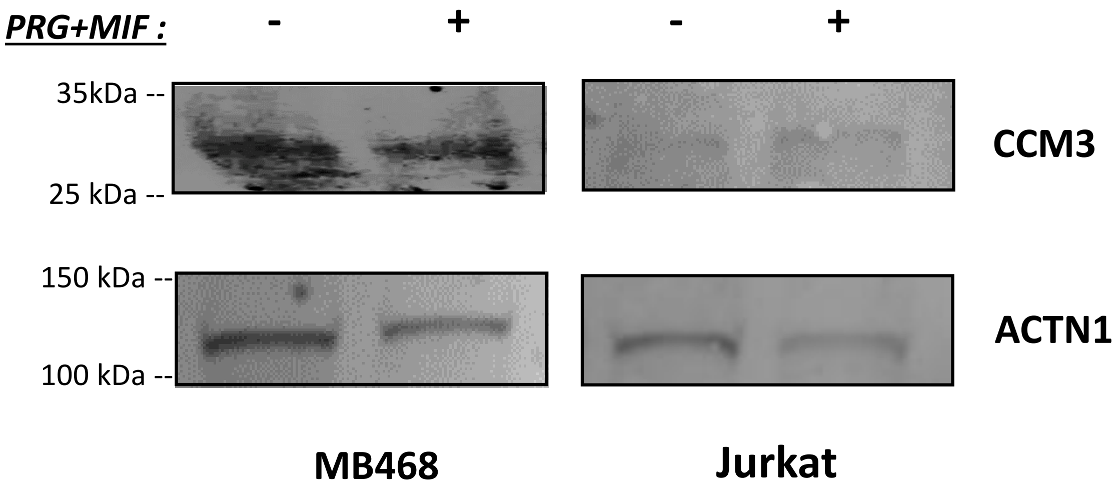
PRG+MIF :
-
+
-
+
35kDa --
25 kDa --
150 kDa --
100 kDa --
CCM3
ACTN1
MB468
Jurkat

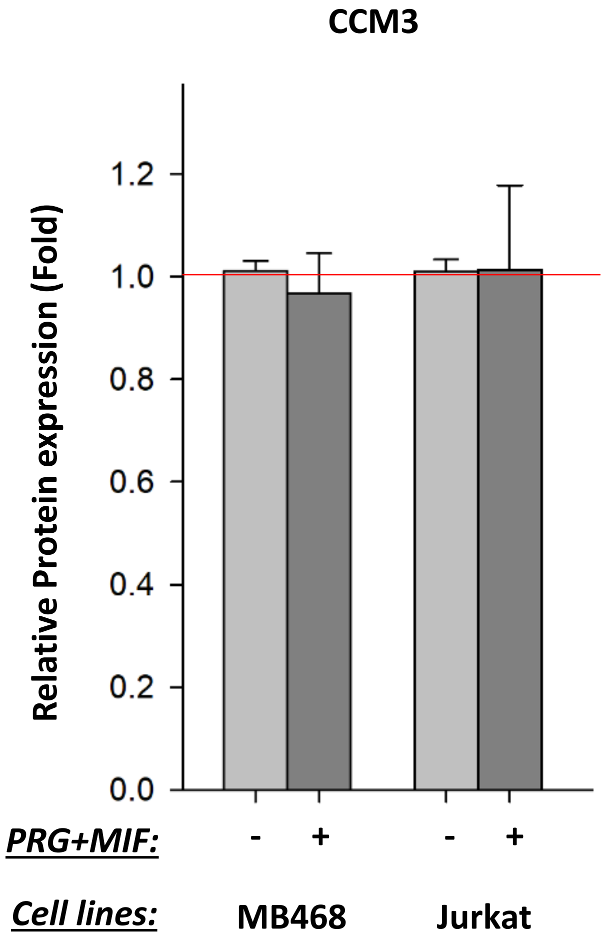
CCM3
Relative Protein expression (Fold)
1.2
1.0
0.8
0.6
0.4
0.2
0.0
PRG+MIF:
-
+
-
+
Cell lines:
MB468
Jurkat

A.

### MB231-MIF treated

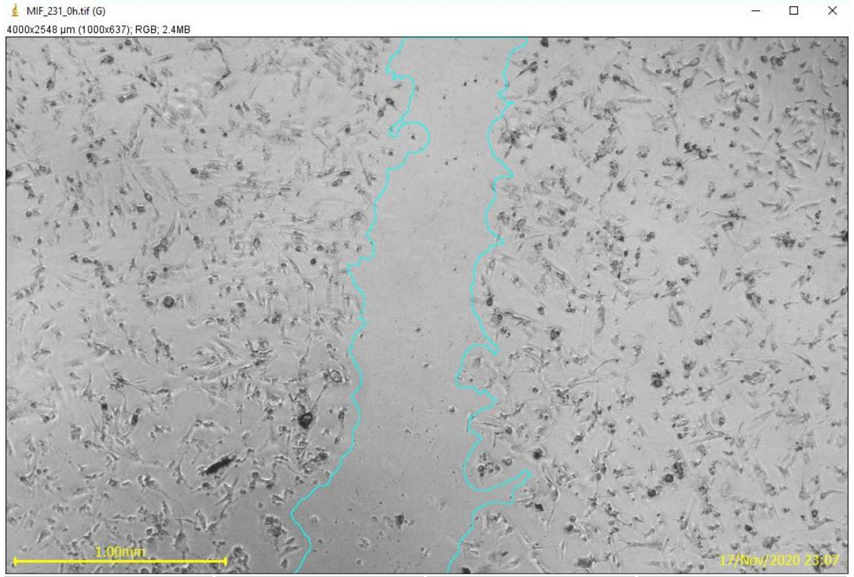


0 hrs

| Label | Area µm² | Width µm | S.D. (µm) |
|---|---|---|---|
| MIF-0 hrs | 1474192 | 581.521 | 142.408 |

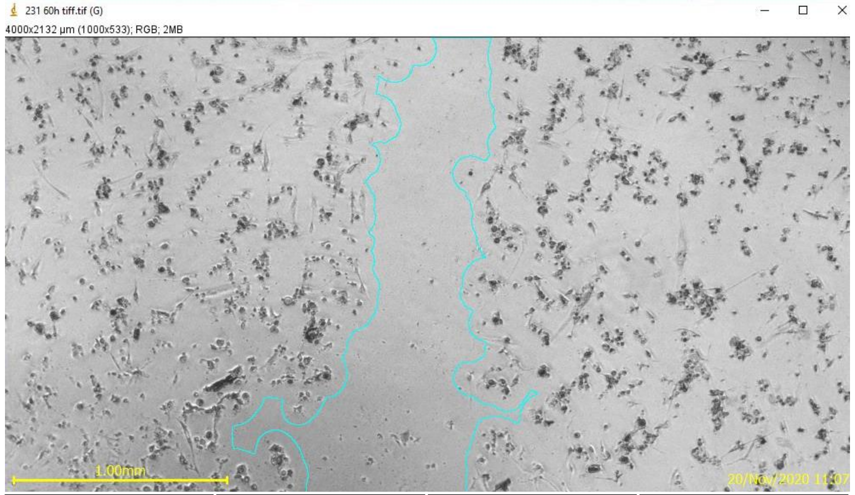


60 hrs

| Label | Area µm² | Width µm | S.D. (µm) |
|---|---|---|---|
| MIF-60 hrs | 1165856 | 563.853 | 259.838 |

B. **MB468-MIF** **MB468-MIF+PRG treated**

**0 hrs**

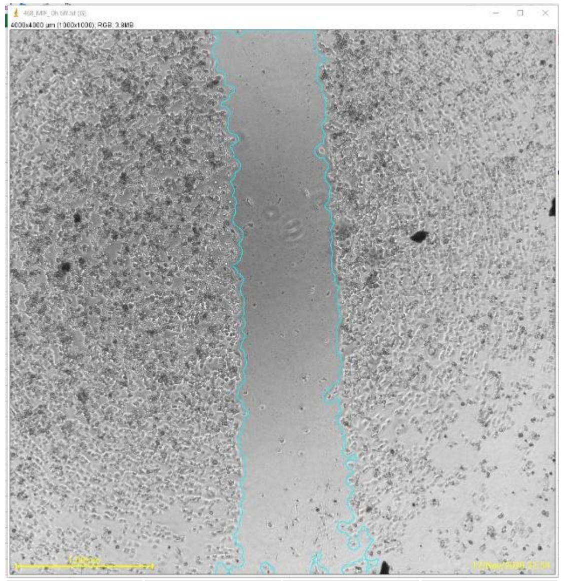

| Label | Area µm² | Width µm | S.D. (µm) |
|---|---|---|---|
| MIF-0 hrs | 2871712 | 727.547 | 95.473 |

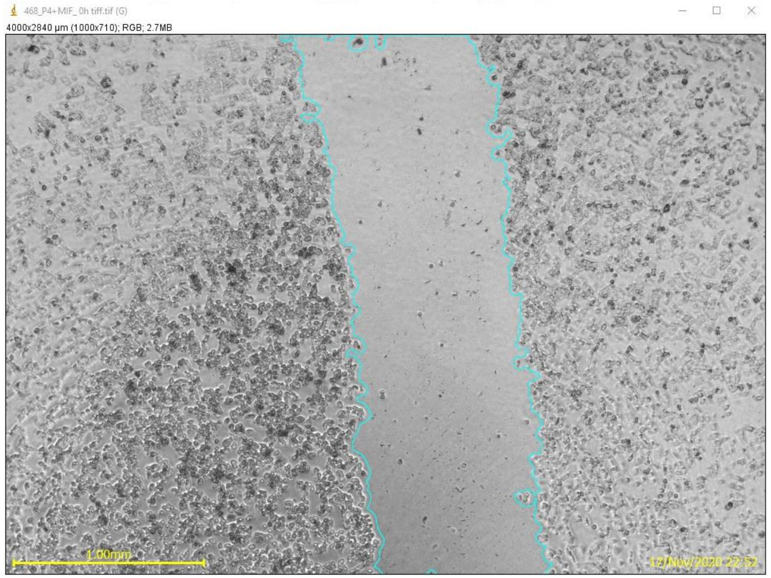


| Label | Area µm² | Width µm | S.D. (µm) |
|---|---|---|---|
| PRG+MIF-0 hrs | 2514992 | 880.706 | 58.854 |

**60 hrs**

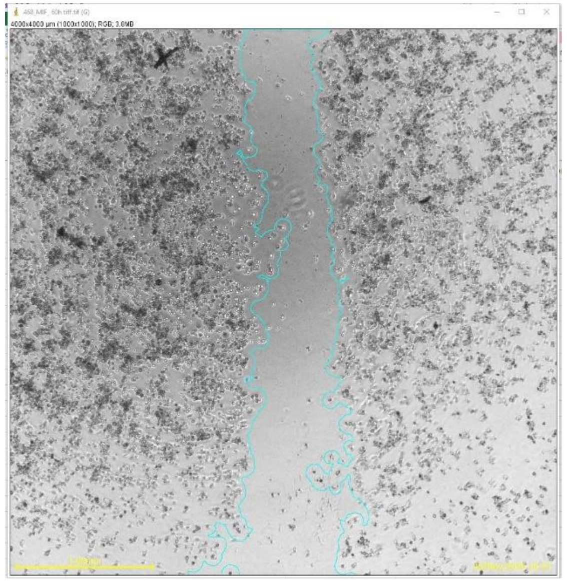

| Label | Area µm² | Width µm | S.D. (µm) |
|---|---|---|---|
| MIF-60 hrs | 2264544 | 592.330 | 167.652 |

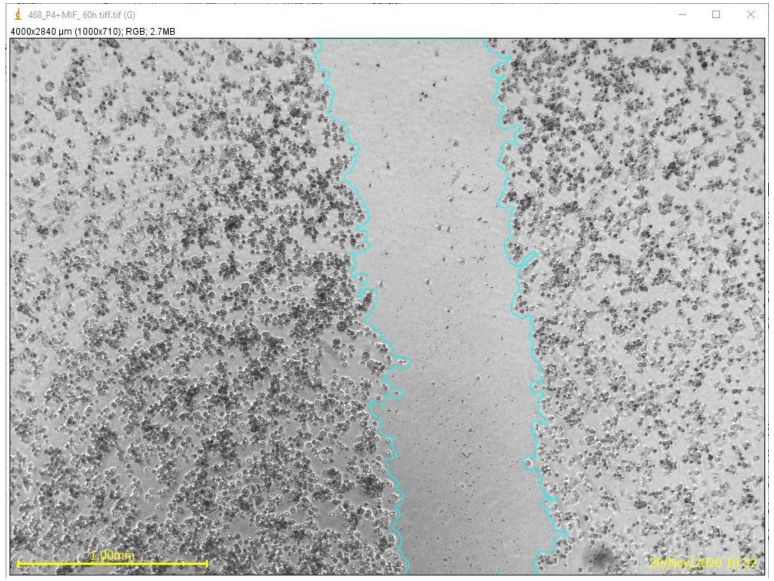


| Label | Area µm² | Width µm | S.D. (µm) |
|---|---|---|---|
| PRG+MIF-60 hrs | 2282208 | 799.382 | 73.915 |

3A

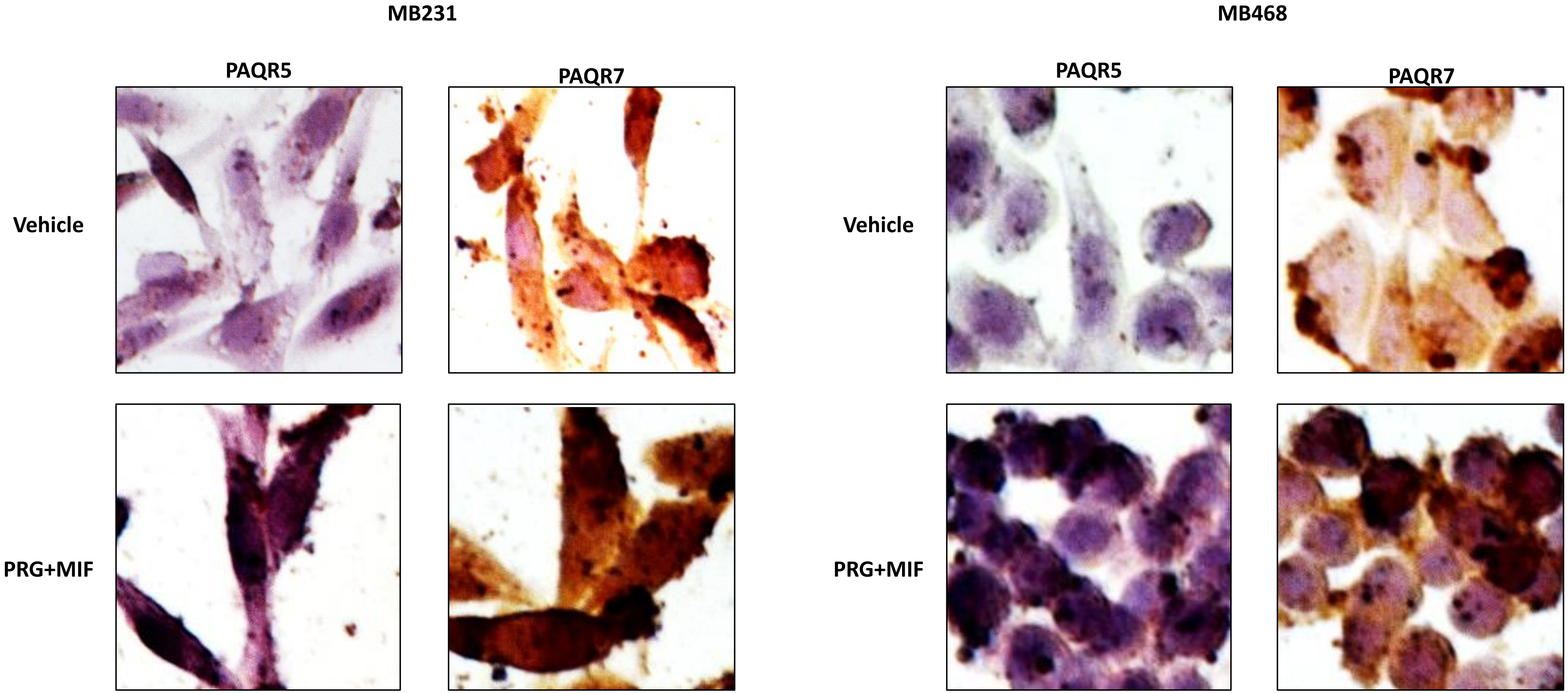
MB231
MB468
PAQR5
PAQR7
PAQR5
PAQR7
Vehicle
Vehicle
PRG+MIF
PRG+MIF

1 mttailerls tlsvsgqql**r rlpkiledgl pkmpctvpet dvpqlf**repy irtgyrptgh
61 ewryyffslf qkhnevvnvw thl**laalavl l**rfwafaeae alpwasthsl p**lllfilssi**
121 tyltcsllah llqskselsh ytfyfvdyvg vsvyqygsal ahffyssdqa wydrfwlffl
181 paaafcgwls cagccyakyr yrrpypvmrk icqvvpagla fildispvah rvalchlagc
241 qeqaawyhtl qilfflvsay ffscpvpeky fpgscdivgh ghqifhafls ictlsqleai
301 lldyqgrqei flqrhgplsv hmaclsfffl aacsaataal lrhkvkarlt kkds

S4A

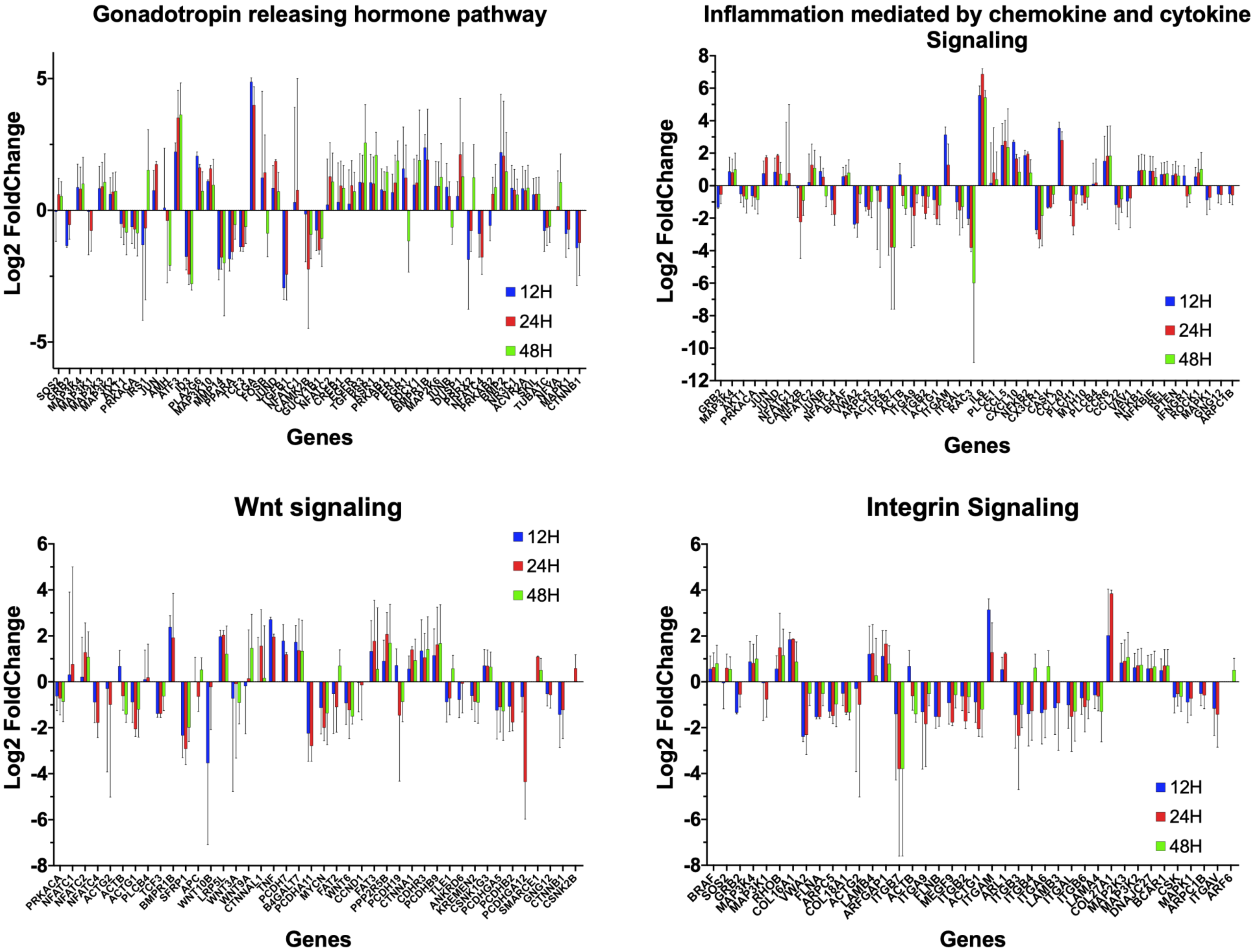

Gonadotropin releasing hormone pathway
Log2 FoldChange
5
0
-5
12H
24H
48H
SOS2 GRB2 MAP3K4 MAP3K1 MAP2K3 MAP3K2 AKT1 PRKACA IRS1 JUN AMH ATF3 ID3 PLA2G6 MAP3K10 MMP14 PPARA TCF3 CGA FOSB JUND TGFB1 NFATC1 CAMK2B GUCY1B1 NFATC2 CREB1 EGFR TGFBR3 KSR1 PRKAB1 PER1 EGR1 ADCY1 BMPR1B MAP2K6 JUNB DUSP1 PRRX2 NFATC4 PRKAB2 BMP2 NR3C1 ACVR2A SKIL TUBA1C NFYA MAPK1 CTNNB1
Genes
Inflammation mediated by chemokine and cytokine Signaling
Log2 FoldChange
8
6
4
2
0
-2
-4
-6
-8
-10
-12
12H
24H
48H
GRB2 MAP3K4 AKT1 PRKACA JUN JUND NFATC1 CAMK2B NFATC2 JUNB NFATC4 BRAF VWA2 ARPC5 ACTG2 ITGB7 ACTB ITGA9 ITGB2 ACTG1 ITGAM ITGAL RAC3 IL6 PLCE1 CCL5 CXCL10 NFKB2 CX3CR1 CASK CCL20 PLCH1 MYH10 PLCB4 CCR6 CCL22 VAV1 NFKB1 NFKBIE REL PTEN IFNGR1 ROCK1 MAPK1 GNG12 ARPC1B
Genes
Wnt signaling
Log2 FoldChange
6
4
2
0
-2
-4
-6
-8
12H
24H
48H
PRKACA NFATC1 NFATC2 NFATC4 ACTG2 ACTB ACTG1 PLCB4 TCF3 BMPR1B SFRP1 APC WNT10B LRP5L WNT3A WNT9A CTNNAL1 TNF PCDH7 B4GALT7 PCDHA11 MYCN FAT2 WNT6 CCND1 FAT3 PPP2R5B PCDH19 CTNNA1 CDHR3 PCDHB9 TLE1 ANKRD6 KREMEN2 CSNK1G3 PCDHGA5 PCDHB2 PCDHGA12 SMARCE1 GNG12 CTNNB1 CSNK2B
Genes
Integrin Signaling
Log2 FoldChange
6
4
2
0
-2
-4
-6
-8
12H
24H
48H
BRAF SOS2 GRB2 MAP3K4 MAP3K1 RHOB COL16A1 VWA2 FLNA ARPC5 COL18A1 ACTG2 LAMB4 ARFGAP1 ITGB7 ACTB ITGA9 FLNB MEGF9 ITGB2 ACTG1 ITGAM ARL1 ITGB3 ITGB4 ITGA6 LAMB3 ITGAL ITGB6 LAMA4 COL27A1 MAP2K3 MAP3K2 DNAJC27 BCAR1 CSK MAPK1 ARPC1B ITGAV ARF6
Genes

S4B

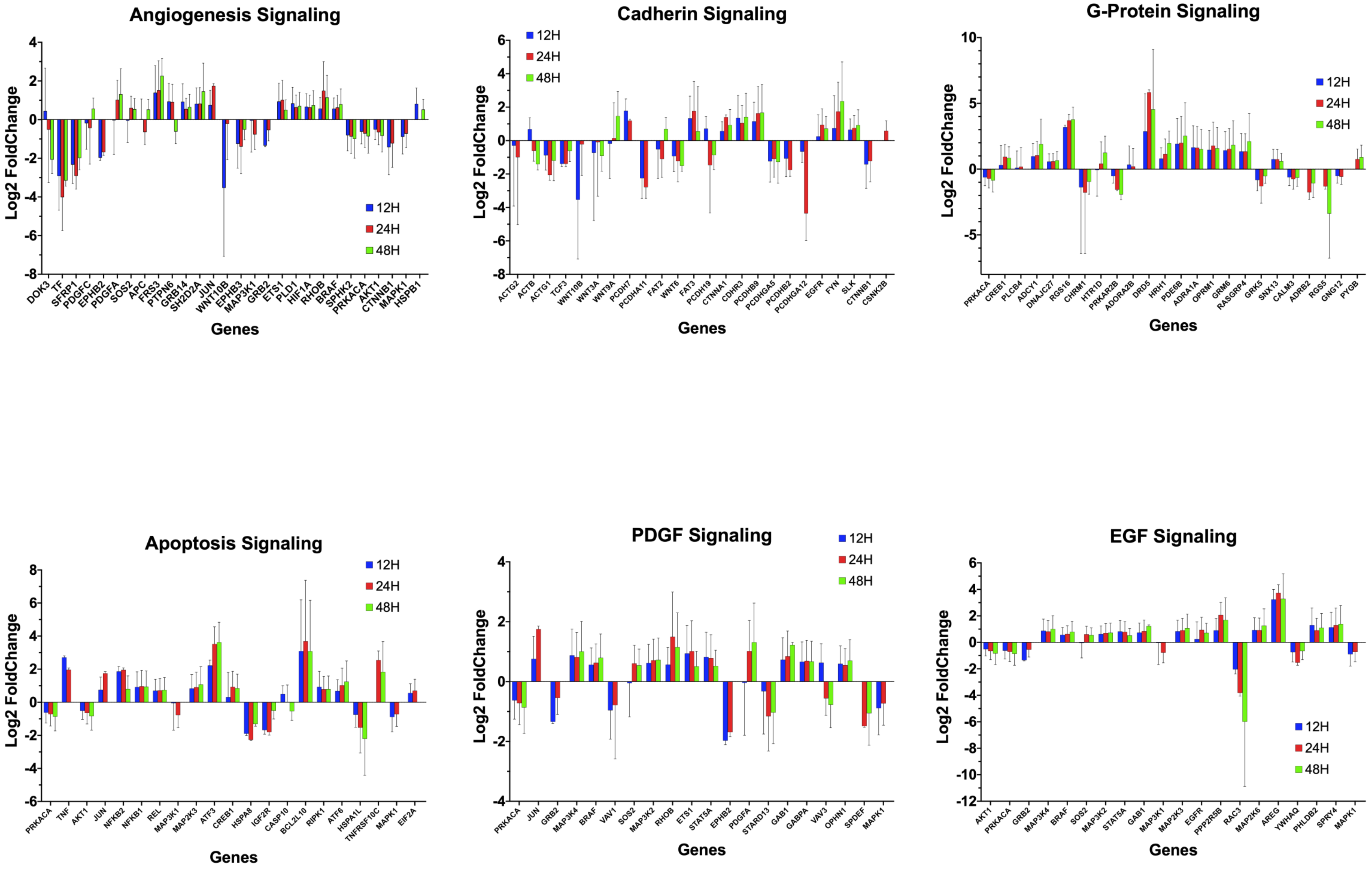

Angiogenesis Signaling
Log2 FoldChange
4 2 0 -2 -4 -6 -8
12H 24H 48H
DOK3 TF SFRP1 PDGFC EPHB2 PDGFA SOS2 APC FRS3 PTPN6 GRB14 SH2D2A JUN WNT10B EPHB3 MAP3K1 GRB2 ETS1 PLD1 HIF1A RHOB BRAF SPHK2 PRKACA AKT1 CTNNB1 MAPK1 HSPB1
Genes
Cadherin Signaling
Log2 FoldChange
6 4 2 0 -2 -4 -6 -8
12H 24H 48H
ACTG2 ACTB ACTG1 TCF3 WNT10B WNT3A WNT9A PCDH7 PCDHA11 FAT2 WNT6 FAT3 PCDH19 CTNNA1 CDHR3 PCDHB9 PCDHGA5 PCDHB2 PCDHGA12 EGFR FYN SLK CTNNB1 CSNK2B
Genes
G-Protein Signaling
Log2 FoldChange
10 5 0 -5
12H 24H 48H
PRKACA CREB1 PLCB4 ADCY1 DNAJC27 RGS16 CHRM1 HTR1D PRKAR2B ADORA2B DRD5 HRH1 PDE6B ADRA1A OPRM1 GRM6 RASGRP4 GRK5 SNX13 CALM3 ADRB2 RGS5 GNG12 PYGB
Genes
Apoptosis Signaling
Log2 FoldChange
8 6 4 2 0 -2 -4 -6
12H 24H 48H
PRKACA TNF AKT1 JUN NFKB2 NFKB1 REL MAP3K1 MAP2K3 ATF3 CREB1 HSPA8 IGF2R CASP10 BCL2L10 RIPK1 ATF6 HSPA1L TNFRSF10C MAPK1 EIF2A
Genes
PDGF Signaling
Log2 FoldChange
4 2 0 -2 -4
12H 24H 48H
PRKACA JUN GRB2 MAP3K4 BRAF VAV1 SOS2 MAP3K2 RHOB ETS1 STAT5A EPHB2 PDGFA STARD13 GAB1 GABPA VAV3 OPHN1 SPDEF MAPK1
Genes
EGF Signaling
Log2 FoldChange
6 4 2 0 -2 -4 -6 -8 -10 -12
12H 24H 48H
AKT1 PRKACA GRB2 MAP3K4 BRAF SOS2 MAP3K2 STAT5A GAB1 MAP3K1 MAP2K3 EGFR PPP2R5B RAC3 MAP2K6 AREG YWHAQ PHLDB2 SPRY4 MAPK1
Genes

S4C

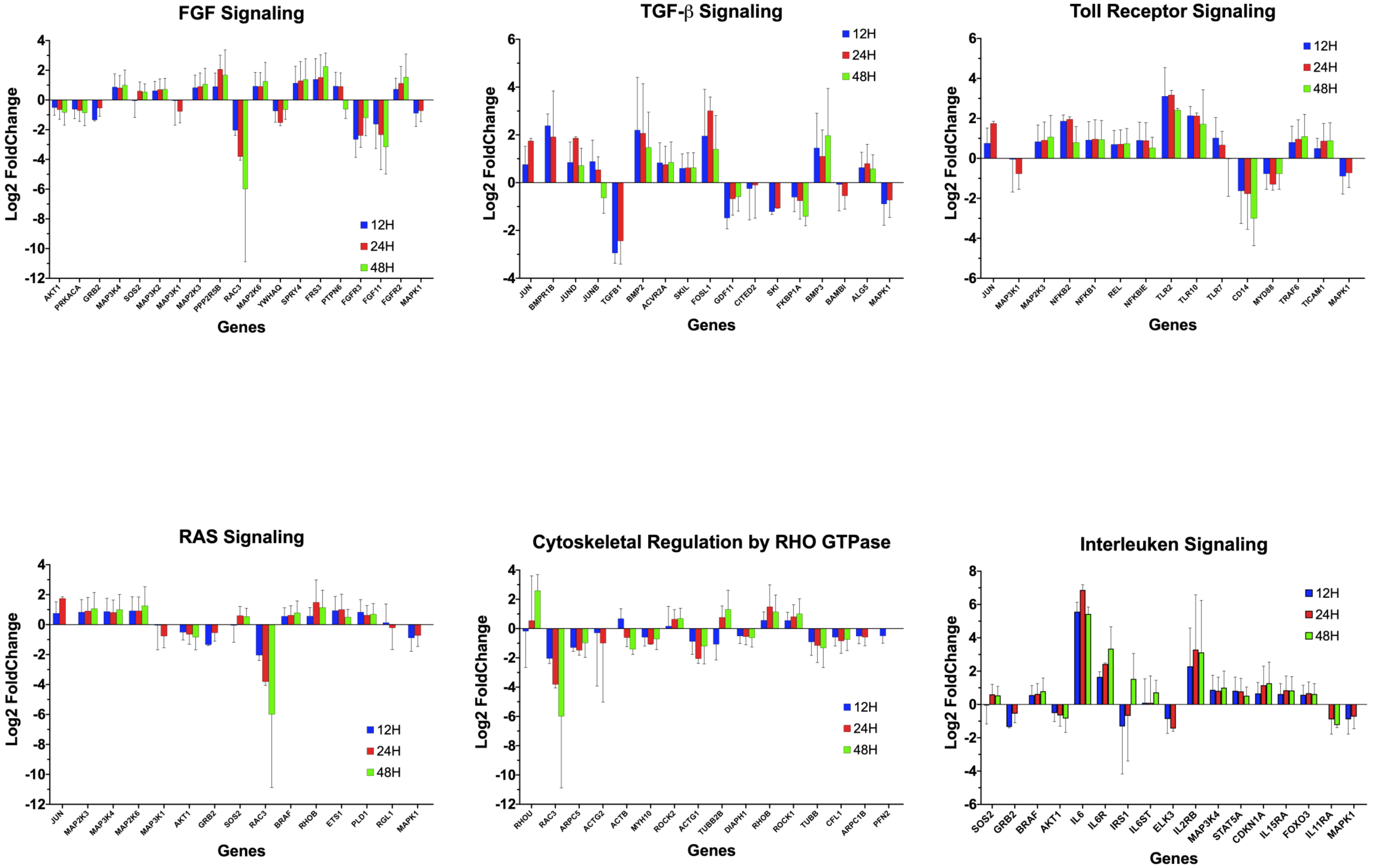

FGF Signaling
Log2 FoldChange
Genes
12H
24H
48H
AKT1
PRKACA
GRB2
MAP3K4
SOS2
MAP3K2
MAP3K1
MAP2K3
PPP2R5B
RAC3
MAP2K6
YWHAQ
SPRY4
FRS3
PTPN6
FGFR3
FGF11
FGFR2
MAPK1
TGF-β Signaling
Log2 FoldChange
Genes
12H
24H
48H
JUN
BMPR1B
JUND
JUNB
TGFB1
BMP2
ACVR2A
SKIL
FOSL1
GDF11
CITED2
SKI
FKBP1A
BMP3
BAMBI
ALG5
MAPK1
Toll Receptor Signaling
Log2 FoldChange
Genes
12H
24H
48H
JUN
MAP3K1
MAP2K3
NFKB2
NFKB1
REL
NFKBIE
TLR2
TLR10
TLR7
CD14
MYD88
TRAF6
TICAM1
MAPK1
RAS Signaling
Log2 FoldChange
Genes
12H
24H
48H
JUN
MAP2K3
MAP3K4
MAP2K6
MAP3K1
AKT1
GRB2
SOS2
RAC3
BRAF
RHOB
ETS1
PLD1
RGL1
MAPK1
Cytoskeletal Regulation by RHO GTPase
Log2 FoldChange
Genes
12H
24H
48H
RHOU
RAC3
ARPC5
ACTG2
ACTB
MYH10
ROCK2
ACTG1
TUBB2B
DIAPH1
RHOB
ROCK1
TUBB
CFL1
ARPC1B
PFN2
Interleuken Signaling
Log2 FoldChange
Genes
12H
24H
48H
SOS2
GRB2
BRAF
AKT1
IL6
IL6R
IRS1
IL6ST
ELK3
IL2RB
MAP3K4
STAT5A
CDKN1A
IL15RA
FOXO3
IL11RA
MAPK1

S4D

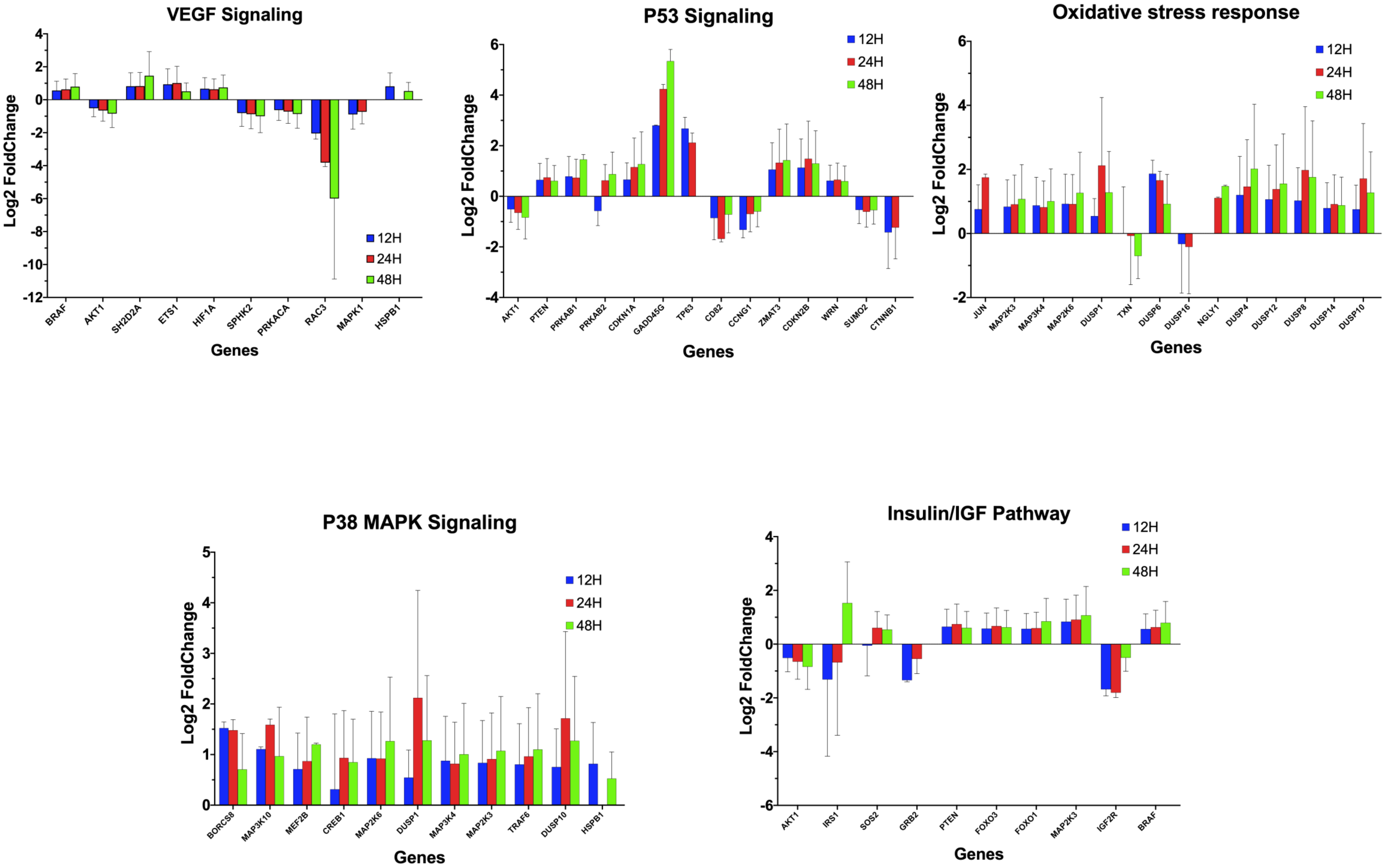
VEGF Signaling
Log2 FoldChange
Genes
BRAF
AKT1
SH2D2A
ETS1
HIF1A
SPHK2
PRKACA
RAC3
MAPK1
HSPB1
12H
24H
48H
P53 Signaling
Log2 FoldChange
Genes
AKT1
PTEN
PRKAB1
PRKAB2
CDKN1A
GADD45G
TP63
CD82
CCNG1
ZMAT3
CDKN2B
WRN
SUMO2
CTNNB1
12H
24H
48H
Oxidative stress response
Log2 FoldChange
Genes
JUN
MAP2K3
MAP3K4
MAP2K6
DUSP1
TXN
DUSP6
DUSP16
NGLY1
DUSP4
DUSP12
DUSP8
DUSP14
DUSP10
12H
24H
48H
P38 MAPK Signaling
Log2 FoldChange
Genes
BORCS8
MAP3K10
MEF2B
CREB1
MAP2K6
DUSP1
MAP3K4
MAP2K3
TRAF6
DUSP10
HSPB1
12H
24H
48H
Insulin/IGF Pathway
Log2 FoldChange
Genes
AKT1
IRS1
SOS2
GRB2
PTEN
FOXO3
FOXO1
MAP2K3
IGF2R
BRAF
12H
24H
48H

S4E

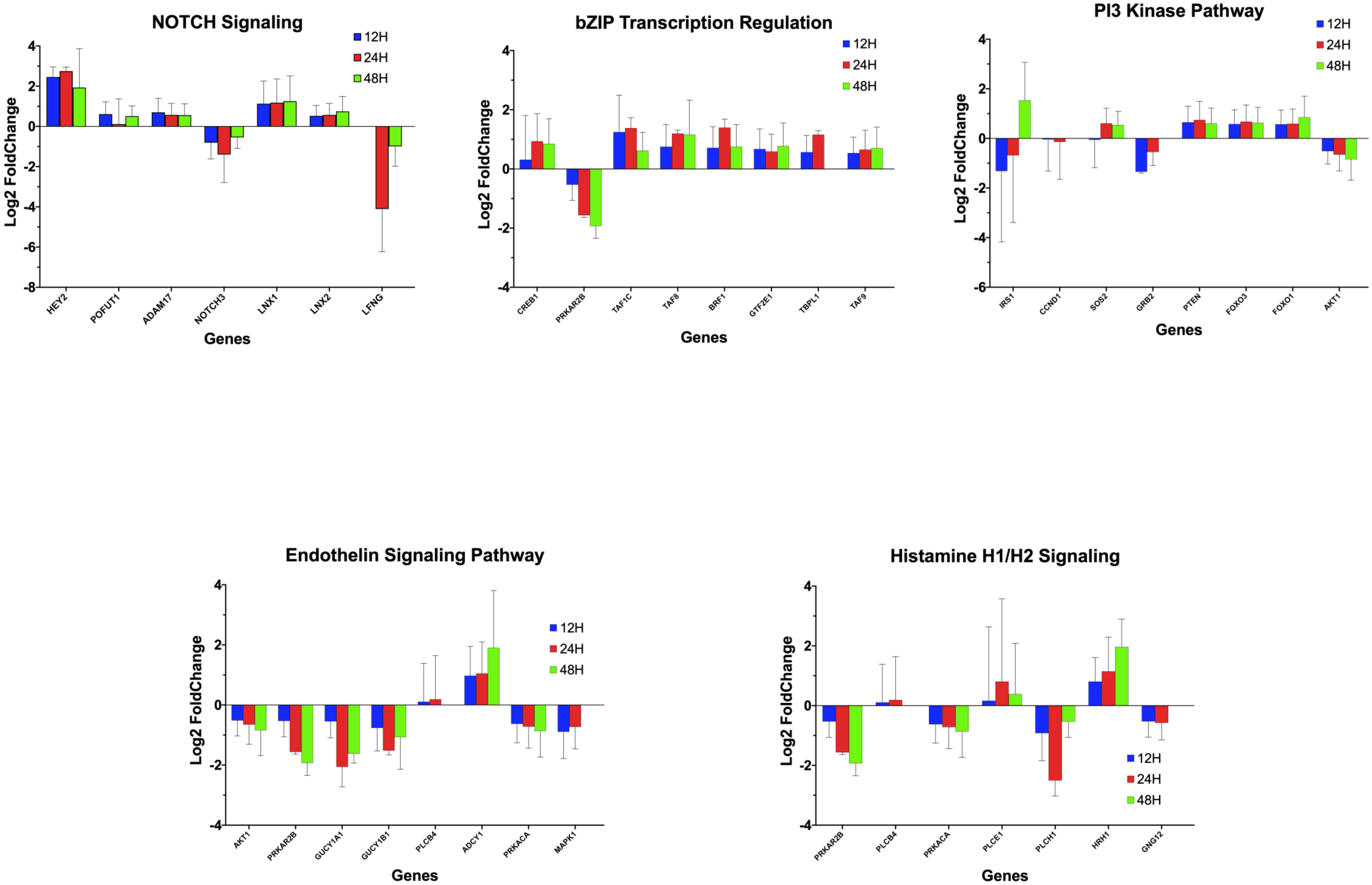
NOTCH Signaling
12H
24H
48H
Log2 FoldChange
4
2
0
-2
-4
-6
-8
HEY2
POFUT1
ADAM17
NOTCH3
LNX1
LNX2
LFNG
Genes
bZIP Transcription Regulation
12H
24H
48H
Log2 FoldChange
4
2
0
-2
-4
CREB1
PRKAR2B
TAF1C
TAF8
BRF1
GTF2E1
TBPL1
TAF9
Genes
PI3 Kinase Pathway
12H
24H
48H
Log2 FoldChange
4
2
0
-2
-4
-6
IRS1
CCND1
SOS2
GRB2
PTEN
FOXO3
FOXO1
AKT1
Genes
Endothelin Signaling Pathway
12H
24H
48H
Log2 FoldChange
4
2
0
-2
-4
AKT1
PRKAR2B
GUCY1A1
GUCY1B1
PLCB4
ADCY1
PRKACA
MAPK1
Genes
Histamine H1/H2 Signaling
12H
24H
48H
Log2 FoldChange
4
2
0
-2
-4
PRKAR2B
PLCB4
PRKACA
PLCE1
PLCH1
HRH1
GNG12
Genes

S5A

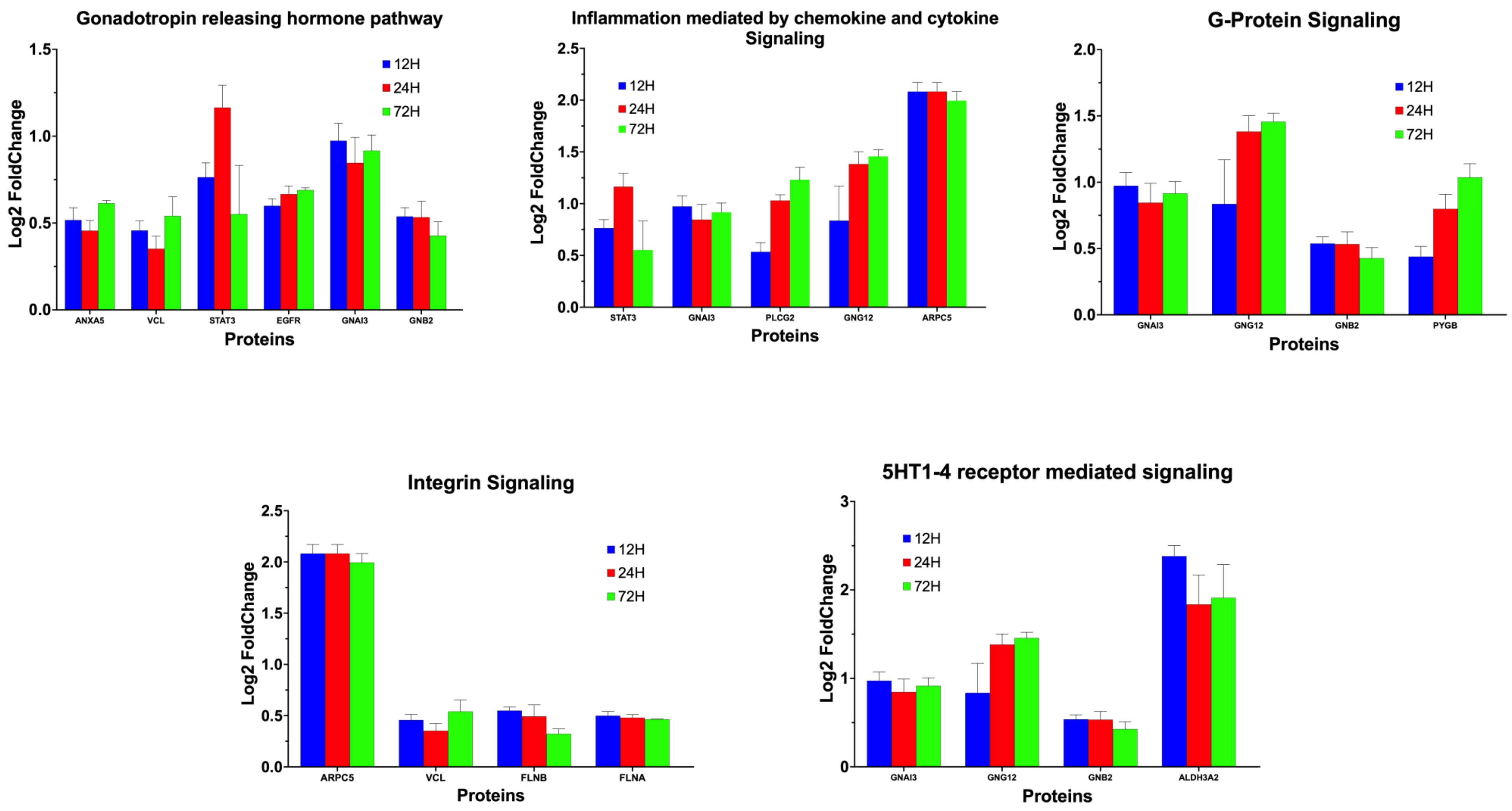
Gonadotropin releasing hormone pathway
Log2 FoldChange
Proteins
12H
24H
72H
ANXA5
VCL
STAT3
EGFR
GNAI3
GNB2
Inflammation mediated by chemokine and cytokine Signaling
Log2 FoldChange
Proteins
STAT3
GNAI3
PLCG2
GNG12
ARPC5
G-Protein Signaling
Log2 FoldChange
Proteins
GNAI3
GNG12
GNB2
PYGB
Integrin Signaling
Log2 FoldChange
Proteins
ARPC5
VCL
FLNB
FLNA
5HT1-4 receptor mediated signaling
Log2 FoldChange
Proteins
GNAI3
GNG12
GNB2
ALDH3A2

S5B

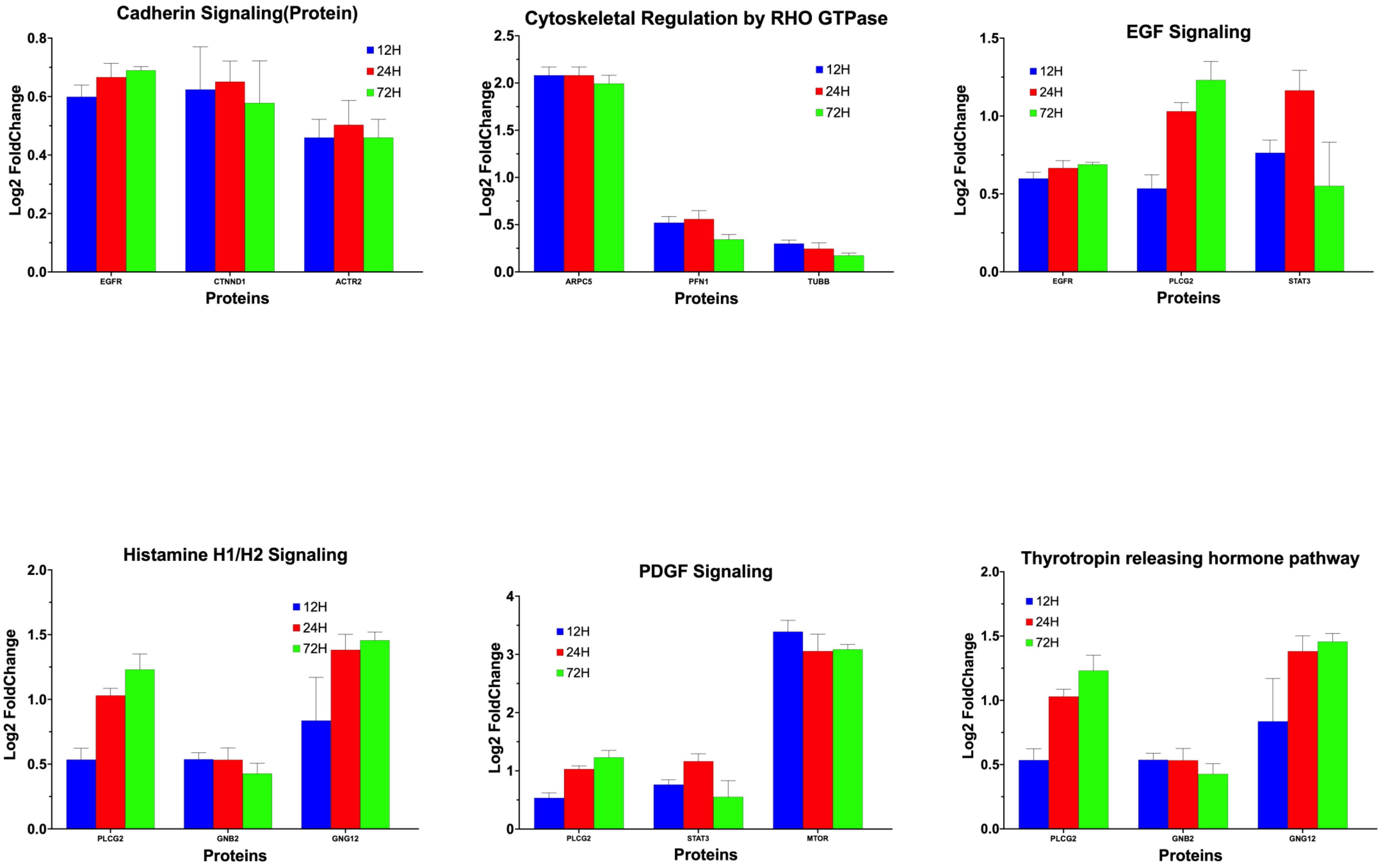
Cadherin Signaling(Protein)
Log2 FoldChange
0.0
0.2
0.4
0.6
0.8
12H
24H
72H
EGFR
CTNND1
ACTR2
Proteins
Cytoskeletal Regulation by RHO GTPase
Log2 FoldChange
0.0
0.5
1.0
1.5
2.0
2.5
12H
24H
72H
ARPC5
PFN1
TUBB
Proteins
EGF Signaling
Log2 FoldChange
0.0
0.5
1.0
1.5
12H
24H
72H
EGFR
PLCG2
STAT3
Proteins
Histamine H1/H2 Signaling
Log2 FoldChange
0.0
0.5
1.0
1.5
2.0
12H
24H
72H
PLCG2
GNB2
GNG12
Proteins
PDGF Signaling
Log2 FoldChange
0
1
2
3
4
12H
24H
72H
PLCG2
STAT3
MTOR
Proteins
Thyrotropin releasing hormone pathway
Log2 FoldChange
0.0
0.5
1.0
1.5
2.0
12H
24H
72H
PLCG2
GNB2
GNG12
Proteins

S5C

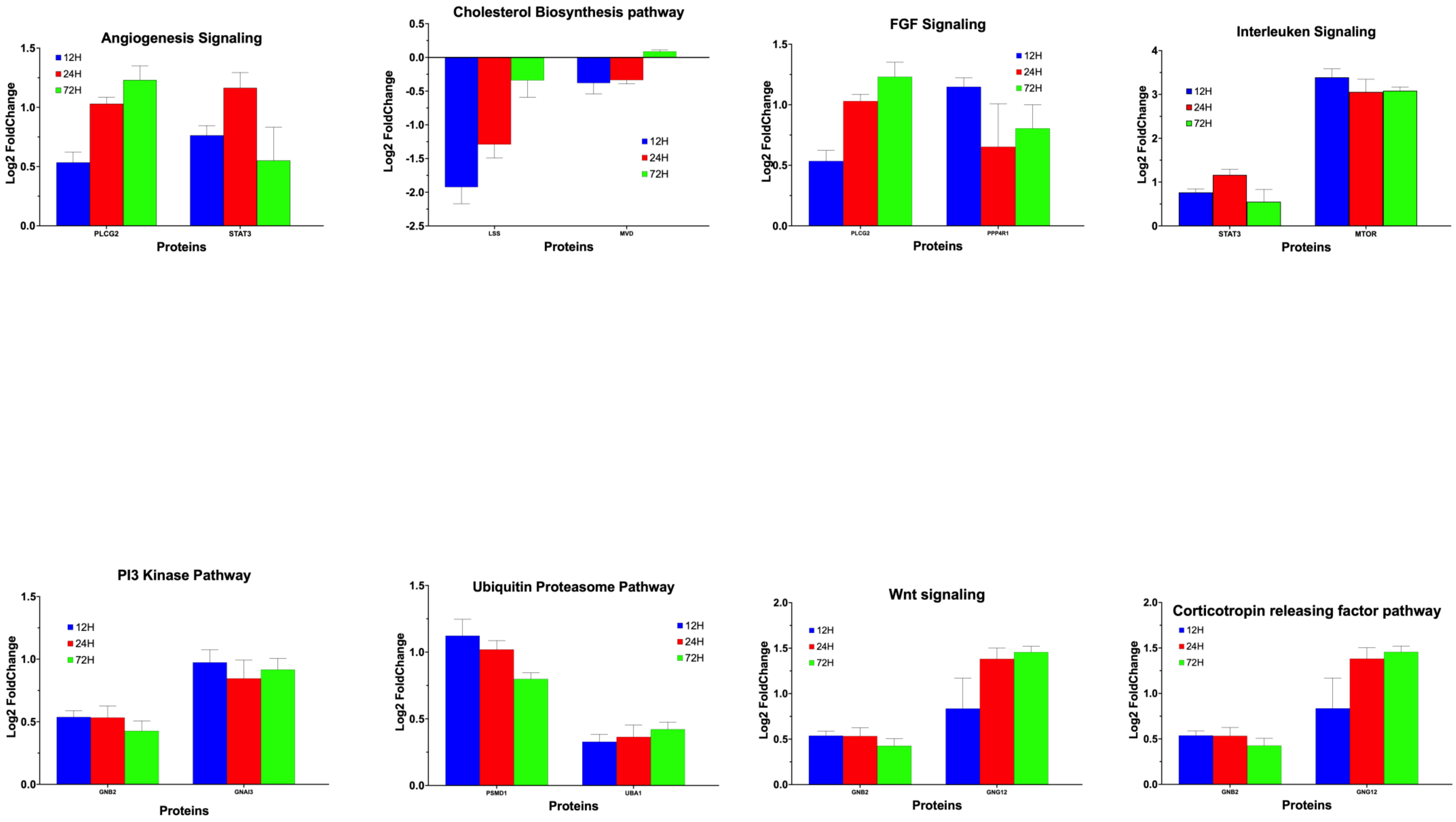
Angiogenesis Signaling
12H
24H
72H
Log2 FoldChange
PLCG2
STAT3
Proteins
Cholesterol Biosynthesis pathway
Log2 FoldChange
LSS
MVD
Proteins
FGF Signaling
Log2 FoldChange
PLCG2
PPP4R1
Proteins
Interleuken Signaling
Log2 FoldChange
STAT3
MTOR
Proteins
PI3 Kinase Pathway
Log2 FoldChange
GNB2
GNAI3
Proteins
Ubiquitin Proteasome Pathway
Log2 FoldChange
PSMD1
UBA1
Proteins
Wnt signaling
Log2 FoldChange
GNB2
GNG12
Proteins
Corticotropin releasing factor pathway
Log2 FoldChange
GNB2
GNG12
Proteins

S5D

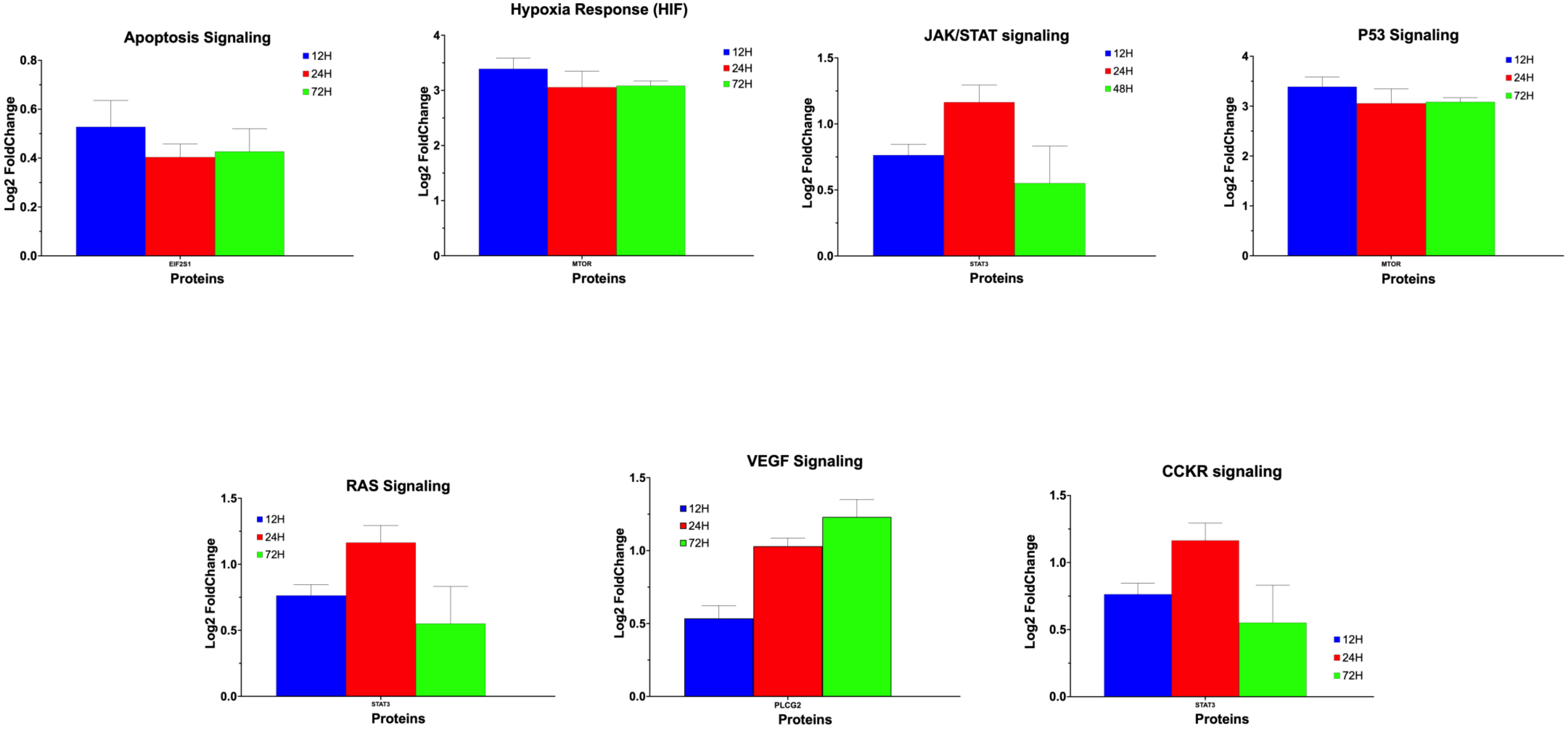
Apoptosis Signaling
Log2 FoldChange
0.0
0.2
0.4
0.6
0.8
12H
24H
72H
EIF2S1
Proteins
Hypoxia Response (HIF)
Log2 FoldChange
0
1
2
3
4
12H
24H
72H
MTOR
Proteins
JAK/STAT signaling
Log2 FoldChange
0.0
0.5
1.0
1.5
12H
24H
48H
STAT3
Proteins
P53 Signaling
Log2 FoldChange
0
1
2
3
4
12H
24H
72H
MTOR
Proteins
RAS Signaling
Log2 FoldChange
0.0
0.5
1.0
1.5
12H
24H
72H
STAT3
Proteins
VEGF Signaling
Log2 FoldChange
0.0
0.5
1.0
1.5
12H
24H
72H
PLCG2
Proteins
CCKR signaling
Log2 FoldChange
0.0
0.5
1.0
1.5
12H
24H
72H
STAT3
Proteins

Suppl. Table 1

| Application | siRNAs | Target | Vendor | Cat # |
|---|---|---|---|---|
| RNAi | ON-TARGETplus Human KRIT1 siRNA - SMARTpool | CCM1 | Dharmacon | L-003825-00-0010 |
| RNAi | KRIT1 (Human) - 3 unique 27mer siRNA duplexes | CCM1 | OriGene | SR313185 |
| RNAi | CCM2 (Human) - 3 unique 27mer siRNA duplexes | CCM2 | OriGene | SAB2500214 |
| RNAi | Ccm2 Rat siRNA Oligo Duplex (Locus ID 305505) | CCM2 | OriGene | SR509691 |
| RNAi | ON-TARGETplus Human CCM2 siRNA - SMARTpool | CCM2 | Dharmacon | L-014728-01-0010 |
| RNAi | ON-TARGETplus Human CCM2l siRNA - SMARTpool | CCM2a | Dharmacon | L-018652-02-0010 |
| RNAi | PDCD10 (Human) - 3 unique 27mer siRNA duplexes | CCM3 | OriGene | sc-365586 |
| RNAi | ON-TARGETplus Human PDCD10 siRNA - SMARTpool | CCM3 | Dharmacon | L-004436-00-0010 |

Suppl. Table 2

| Application | Antigen | Gene | Clone | Secondary | Vendor | Cat # |
|---|---|---|---|---|---|---|
| IF/WB | CCM1 | Krit1 | E-8 | (anti-mouse) | Santa Cruz | sc-514371 |
| IF/WB | CCM1-AF488 | Krit1 | E-8 | (anti-mouse) | Santa Cruz | sc-514371 |
| IF/WB | CCM1 | Krit1 | 8-RY2 | (anti-mouse) | Santa Cruz | sc-134376 |
| IF/WB | CCM1 | Krit1 | | (anti-rabbit) | OriGene | AP26021PU-L |
| IF/WB | CCM3 | PDCD10 | C-8 | (anti-mouse) | Santa Cruz | sc-365586 |
| IF/WB | CCM3-AF647 | PDCD10 | C-8 | (anti-mouse) | Santa Cruz | sc-365586 |
| IF/WB | CCM3-AF546 | PDCD10 | C-8 | (anti-mouse) | Santa Cruz | sc-365586 |
| IF/WB | CCM3 | PDCD10 | F-12 | (anti-mouse) | Santa Cruz | sc-365587 |
| WB | PAQR5 | mPRγ | | (Anti-Rabbit) | Aviva | OASG04642 |
| WB | PAQR5/6 | mPRδ/γ | B-8 | (anti-mouse) | Santa Cruz | sc-514273 |
| WB | PAQR7 | mPRα | | (Anti-Rabbit) | Aviva | OASG04641 |
| WB | PAQR7 | mPRα | | (Anti-Rabbit) | Aviva | ARP67727 |
| IF | PAQR8 | mPRβ | | (Anti-Rabbit) | Aviva | OAAB11180 |
| WB | PAQR8 | mPRβ | | (Anti-Rabbit) | Aviva | ARP66903 |
| WB | β-actin | | C-2 | (anti-mouse) | Santa Cruz | sc-8432 |
| WB | α-actinin | | H-2 | (anti-mouse) | Santa Cruz | sc-17829 |
| WB | α-actinin-AF488 | | H-2 | | Santa Cruz | sc-17829 |
| WB | Rabbit-HRP | | | | Santa Cruz | sc-2357 |
| WB | Mouse-HRP | | m-IgGκ BP | | Santa Cruz | sc-516102 |
| WB | m-IgGκ BP IgG-CFL 488 | | m-IgGκ BP | | Santa Cruz | sc-516176 |
| WB | Rabbit IgG-CFL 488 | | | | Santa Cruz | sc-516248 |
| IF | mouse anti-rabbit IgG-CFL 488 | | | secondary | Santa Cruz | sc-516248 |
| IF | mouse anti-rabbit IgG-CFL 555 | | | secondary | Santa Cruz | sc-516249 |
| IHC/IF | DAPI | | | | Santa Cruz | sc-3598 |

Suppl. Table 1

| Target Gene | Primer | Sequence | Product Size (bp) | Anneling Temp (°C) |
|---|---|---|---|---|
| CCM1 | CCM1p1-F1 | AACTGGGAAGAAGCTGCAAA | 156 | 60 |
| | CCM1p1-R1 | AGAGAGACGCATTCCTTCCA | | |
| CCM3 | PDCD102F1 | ATGAGGATGACAATGGAAGAGATGAAG | 307 | 65 |
| | PDCD101R2 | GGTCTTGGAATTCTGGCTCTGGTCGTTC | | |
| CCM2-5 | MGC-EXB1-2-F1 | GGCAAGAAGCCTGGAATTGT | 98 | 65 |
| | MGC-EX2-R1 | CGCCTCTCTGTCACCTTCTC | | |
| CCM2-6 | MGC-EXB1-3-F1 | GGGCAAGAAGTATTTAGGTCAG | 103 | 60 |
| | MGC-EXB3-4-R1 | GTGGGCTCTCTTTGCATTGT | | |
| CCM2-17 | MGC-EXB6-7as01-F1 | TGCACAGCGATGACTCTTCT | 89 | 65 |
| | MGC-EX10as04-R1 | GCCAAAGCTGTCAGTGATGA | | |
| CCM2-37 | MGC-EXB1A-2-F1 | GGAGAATGAGCCTGGAATTG | 120 | 65 |
| | MGC-EX2-R2 | AACACCACAGTGTGCAGAGG | | |
| CCM2-48 | MGC-EXB6A-7-F2 | CTCCCTATTTCAAGATGACTCTTCTAC | 103 | 60 |
| | MGC-EX8-R2 | ACCACCCACATCCACAGATT | | |
| mPRα/PAQR7 | PAQR7-F1 | cgctcttctggaagccgtacatctatg | 122 | 65 |
| | PAQR7-R1 | cagcaggtgggtccagacattcac | | |
| mPRβ/PAQR8 | PAQR8-F1 | agcctcctacata gatgctgccc | 194 | 65 |
| | PAQR8-R1 | ggtgcctggttcacatgttcttca | | |
| mPRγ/PAQR5 | PAQR5-F1 | cagctgtttcacgtgtgtgtgatcctg | 144 | 65 |
| | PAQR5-R1 | gcacagaag tatggctccagctatctgag | | |
| mPRδ/PAQR6 | PAQR6-F2 | gttgacccaccagcttagga | 176 | 65 |
| | PAQR6-R2 | atgccatcttcccagaacac | | |
| mPRε/PAQR9 | PAQR9-F1 | TGCTACAAAGGGATCCCAAC | 202 | 65 |
| | PAQR9-R1 | TGGCACAGATGATTGGAAAA | | |
| PGRMC1 | PGRMC1-F1 | CTGCATGATTTCTGTTTTATCTACCTCTA | 86 | 65 |
| | PGRMC1-R1 | TGTTACTGGACAGCGCTTAATCC | | |
| AR | AR-F1 | cct ggc ttc cgc aac tta cac | 168 | 65 |
| | AR-R1 | gga ctt gtg cat gcg gta ctc a | | |
| GR | GR-F1 | tc aaa aga gca gtg gaa gg | 260 | 65 |
| | GR-R1 | ggt agg ggt gag ttg tgg taa cg | | |
